# Gene and allele specific expression underlying the electric signal divergence in African weakly electric fish

**DOI:** 10.1101/2023.10.30.564736

**Authors:** Feng Cheng, Alice B. Dennis, Otto Baumann, Frank Kirschbaum, Salim Abdelilah-Seyfried, Ralph Tiedemann

## Abstract

In the African weakly electric fish genus *Campylomormyrus*, electric organ discharge (EOD) signals are strikingly different in shape and duration among closely related species, contribute to pre-zygotic isolation and may have triggered an adaptive radiation. We performed mRNA sequencing on electric organs (EOs) and skeletal muscles (SMs; from which the EOs derive) from three species with short (0.4 ms), medium (5 ms), and long (40 ms) EODs and two different cross-species hybrids. We identified 1,444 up-regulated genes in EO shared by all five species/hybrids cohorts, rendering them candidate genes for EO-specific properties in *Campylomormyrus*. We further identified several candidate genes, including *KCNJ2* and *KLF5*, their up-regulation may contribute to increased EOD duration. Hybrids between a short (*C. compressirostris*) and a long (*C. rhynchophorus*) discharging species exhibit EODs of intermediate duration and showed imbalanced expression of *KCNJ2* alleles, pointing towards a cis-regulatory difference at this locus, relative to EOD duration. *KLF5* is a transcription factor potentially balancing potassium channel gene expression, a crucial process for the formation of an EOD. Unraveling the genetic basis of the species-specific modulation of the EOD in *Campylomormyrus* is crucial for understanding the adaptive radiation of this emerging model taxon of ecological (perhaps even sympatric) speciation.

## Introduction

Electric fish have independently evolved six times (Darwin 1859; Bass 1986; Kirschbaum and Formicki 2020). They possess a specific myogenic electric organ (EO) derived from skeletal muscle (SM) fibers except for Apteronotidae which possess an EO derived from nervous tissue (Smith 2013). Comparative genomics have unraveled this convergent phenotypic evolution to originate in part also from convergence on the molecular level: both voltage-dependent sodium and potassium channels are involved in the electric organ development and physiology. Because of the teleost-specific whole genome duplication (Glasauer and Neuhauss 2014), these fish possess two copies of most genes and subfunctionalization among paralogs and differential expression between EO and SM seem to play a major role in the transition of myocytes to electrocytes. A prominent example is the voltage-gated sodium (*Na_v_*) channel gene (*SCN4a*): convergently in three electrogenic taxa (Mormyridea, Siluriformes and Gymnotiformes), only one paralog (*SCN4ab*) is still expressed in SM, but the other one (*SCN4aa*) is exclusively expressed in the EO, indicating a crucial role for electrogenesis (Zakon 2012; Wang and Yang 2021; LaPotin et al. 2022). The *Na_v_* channel (*SCN4aa*) is regulated by *FGF13a* in the three electric fish lineages Siluriformes, Gymnotiformes, and Mormyroidea (Gallant et al. 2014).

Differential expression of multiple isoforms of α and β subunits of sodium/potassium ATPase is important in the EO as well (Gallant et al. 2012; Gallant et al. 2014; Lamanna et al. 2015). In addition, several transcription factors, *HEY1*, *MEF2a*, *SIX2a*, are convergently up-regulated in the EOs of those electric fishes lineages (Kim et al. 2004; Gallant et al. 2012; Gallant et al. 2014). EOs hence comprise a prime example of convergent evolution in both genotype and phenotype.

One of the electric fish clades, mormyrid fish, contains about 200 described species that are endemic to Africa. This outstanding adaptive radiation within the otherwise species-poor basal lineage of osteoglossiforms is putatively due to their species-specific weak electric signals, which is used for both electrolocation and electrocommunication (Feulner et al. 2009a; Feulner et al. 2009b). Divergence in electric organ discharge (EOD) is considered a major driver in the ecological (and possibly sympatric) speciation in the mormyrid genus *Campylomormyrus*, which is mainly distributed in the Congo River (Tiedemann et al. 2010).

The genus *Campylomormyrus* comprises 15 described species, which have profoundly diverged in their electric organ discharge (EOD) with regard to signal duration and waveform (Feulner et al. 2009a). Those species possess either long or short, biphasic or triphasic, but always species-specific EODs, that function as a pre-zygotic reproductive isolation mechanism and are supposed to have arisen via divergent selection among closely related species (Feulner et al. 2009a). In adult *Campylomormyrus*, the electric organ, confined to the caudal peduncle (Fig. 1a), is composed of specialized electrocytes (Paul et al. 2015). They have a flat, disk-shaped appearance with a clear orientation toward the longitudinal body axis (Fig. 1b). Unlike skeletal muscle myocytes, electrocytes possess a number of special evaginations, called stalks, mostly on the posterior face (Paul et al. 2015). These stalks are either fused into major stalks on the posterior face (Fig. 1c left) or they penetrate the electrocyte and merge at the anterior face to constitute to major stalks (Fig. 1c right). A branch of the spinal nerve forms numerous synapses with the major stalk, whether on the posterior or on the anterior face of the electrocyte, and the action potentials are propagated along the stalk system to the disc-like part of the electrocyte (Paul et al. 2015). The externally measurable EOD is formed by simultaneous action potentials of all electrocytes. The shape of the EOD in *Campylomormyrus* is often associated with the penetration of the stalks (Gallant et al. 2011), while the structural basis of the EOD duration, which can vary 100-fold across species, is still only partially understood. A very elongated EOD (∼40 ms) is produced by *C. rhynchophorus* and *C. numenius* which exhibit large foldings or evaginations on the anterior face of the electrocytes, so called papillae (Kirschbaum et al. 2016; Korniienko et al. 2021). In two species with relatively short EOD (Fig. 2a), *C. compressirostris* (0.4 ms) and *C. tamandua* (0.4 ms), many small stalks fuse into one major stalk of large diameter after their origin (Paul et al. 2015). In contrast, the stalk system in species with an EOD of medium (e.g. *C. tshokwe*, 5 ms), or long duration (e.g. *C. numenius*, 40 ms) is more branched (Paul et al. 2015). Apart from these difference in the stalk system, species with highly diverged EOD waveforms still show similar electrocyte geometry suggesting further core mechanisms to contribute to the observed EOD variations. Since the electrocytes generate action potentials for EOD, the distribution and repertoire of ion currents have long been considered to play a key role in EOD formation (Lamanna et al. 2014; Lamanna et al. 2015; Paul et al. 2016; Nagel et al. 2017; Swapna et al. 2018).

**Fig. 1.**
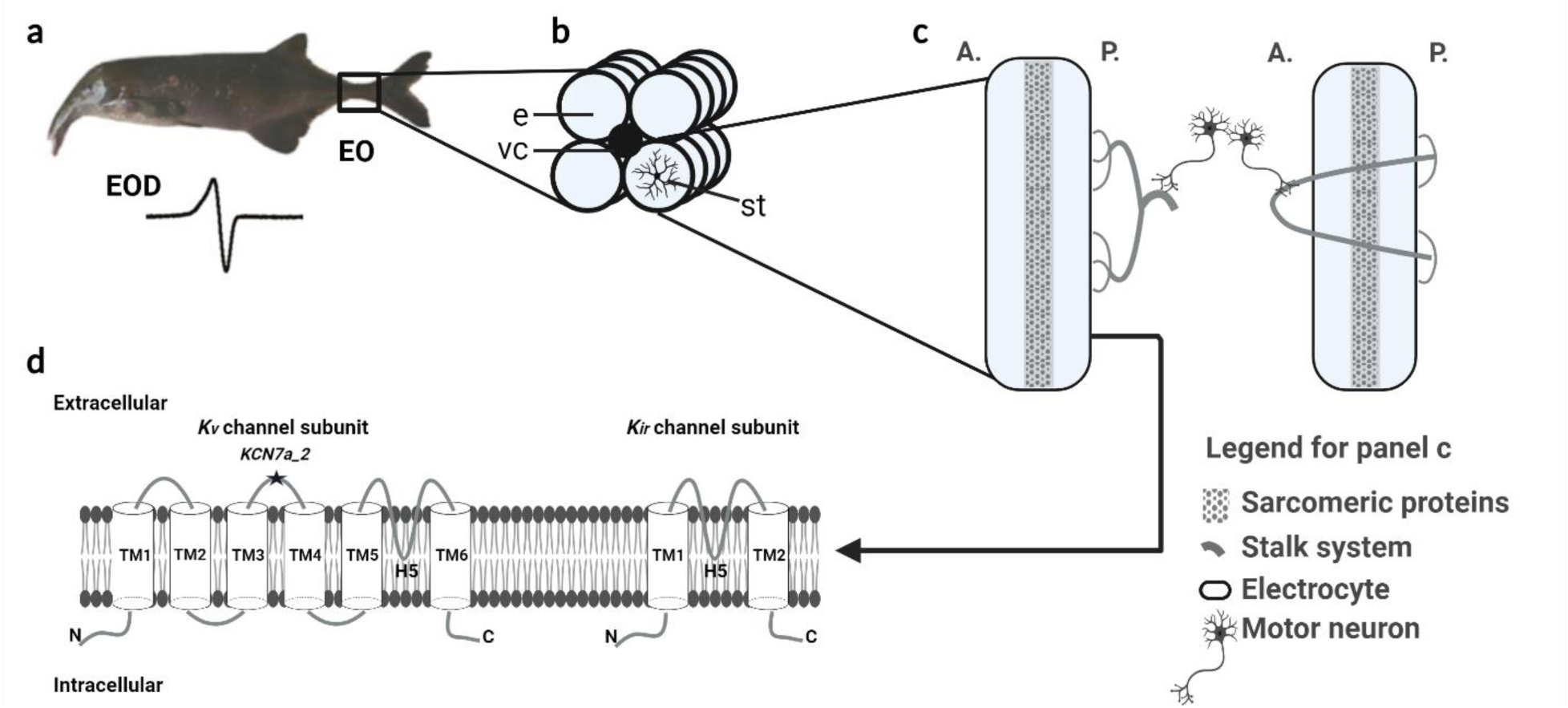
Electric organ and electrocyte structure in *Campylomormyrus*, and schematic illustration for potassium channels. **a** Electric organ (EO) and electric organ discharge (EOD) in an adult *Campylomormyrus* fish. **b** The EO consists of four columns of electrocytes (e) which surround the vertebral column (vc), the stalk system (st) is connected to the posterior face of the electrocyte. **c** Anterior (A.) and posterior (P.) faces of electrocytes with two types of stalk system. Panel c is modified from *Gallant et al.* 2012. **d** Schematic illustration of voltage-gated potassium (*K_v_*) channel and inwardly rectifying (*K_ir_*) channel subunit. *K_v_* channel subunit contains six transmembrane (TM) helices, a pore-forming (H5) loop, and cytosolic NH_2_ (N) and COOH (C) termini. The gene *KCN7A_2* was inferred to be under positive selection and the mutation encodes the loop between TM3-4. *K_ir_* channel subunit contains only two TMs.

**Fig. 2.**
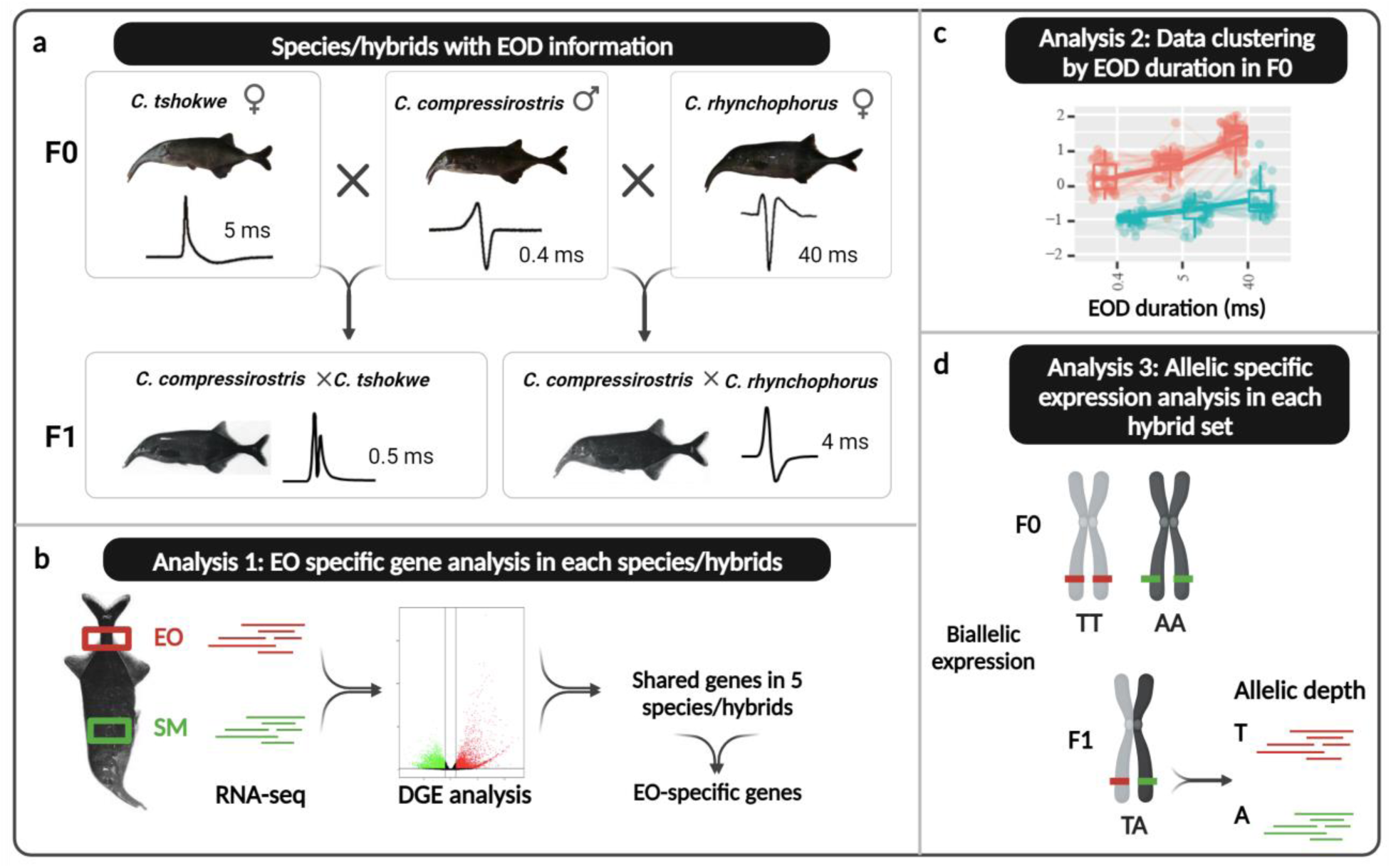
Electric organ discharge (EOD) shape and duration of *Campylomormyrus* species and hybrids, and the working flow of this study. **a** Species/hybrids samples used in the study and their EOD pattern. **b** Differential gene expression (DGE) analysis between electric organ (EO) and skeletal muscle (SM) for each species/hybrid to identify genes with EO-specific expression. **c** RNA-seq data clustering based on EOD duration change EO (red) and SM (blue) in F0 species. **d** Allele specific expression analysis in each hybrid set.

Sodium and potassium fluxes are considered the most important ion currents in controlling the EOD (Stoddard and Markham 2008). They are the basic requirements for generating an action potential (Mehaffey et al. 2006). Consequently, abundance and properties of sodium and potassium channels are likely to profoundly influence the EOD. The potassium channels can be classified into different classes based on their structure and function: voltage-gated (*K_v_*, includes subfamilies e.g. *shaker*-related *KCNA*, *shab*-related *KCNB*), inwardly rectifying (*K_ir_*), tandem pore domain channels (*K_2p_*), ligand-gated channels and calcium-activated channels (*K_ca_*) (Kuang et al. 2015). Two paralogs of the *KCNA7* channel gene originate from the whole genome duplication event in teleost fish and these paralogs might have undergone subfunctionalization or neofunctionalization in mormyrids: one of them *KCNA7a* is predominantly expressed in the EO of mormyrids, while *KCNA7b* is preferentially expressed in SM (Swapna et al. 2018). The *K_v_* channel contains six transmembrane helices (Fig. 1d). In *KCNA7a*, a non-synonymous substitution was observed in the transmembrane helices 3-4 linker and the encoded amino acid substitution might relate to the EOD duration difference among the mormyrid taxa *Brienomyrus* and *Gymnarchus* (Swapna et al. 2018).

This study focusses on potential molecular mechanisms underlying the divergent EOD among *Campylomormyrus* species as a potential major driver of their adaptive radiation. This study takes further advantage of artificially bred hybrid electric fish. *Campylomormyrus* species hybrids often exhibit an adult EOD which is similar to the juvenile EOD from one of the parental species, and the adult EOD duration in hybrids is usually intermediate between the two parental species (Kirschbaum et al. 2016). Gene expression analyses in hybrids further enable assessment of allelic specific expression, relative to the expressed trait of interest (here, EOD duration). To enhance our understanding of the genetic regulation of EOD divergence among *Campylomormyrus* species, especially for the EOD duration divergence, we: 1) compared the gene expression pattern between electric organ (EO) and skeletal muscle (SM) in the three F0 species *C. compressirostris* (*com*, short and biphasic EOD), *C. tshokwe* (*tsh*, medium and biphasic EOD), *C. rhynchophorus* (*rhy*, long and triphasic EOD), and two F1 hybrids *C. compressirostris* ♂ x *C. tshokwe* ♀ (*com* x *tsh*, short and biphasic EOD), and *C. compressirostris* ♂ x *C. rhynchophorus* ♀ (*com* x *rhy*, medium and biphasic EOD); 2) clustered RNA-seq data relative to the EOD duration in three F0 species to infer genes with duration-specific expression; 3) assessed biallelic specific expression for two hybrid sets (each set includes two F0 parental species and their hybrid; Fig. 2).

## Results

We examined overall patterns in gene expression using a principal component analysis (PCA) based on all expressed genes (Fig. 3a). Expression profiles of SMs and EOs were broadly separated along PC1, which explained 74% of variance. The SMs profiles from all species/hybrids clustered together; however, species/hybrid-specific EOs’ expression profiles were stratified along PC2 (explained 6% of variance), relative to EOD duration (Fig. 3a). The PCA hence indicates that gene expression in *Campylomormyrus* is (1) EO specific, compared with SM; (2) relates to EOD duration, enabling inference of underlying candidate genes.

**Fig. 3.**
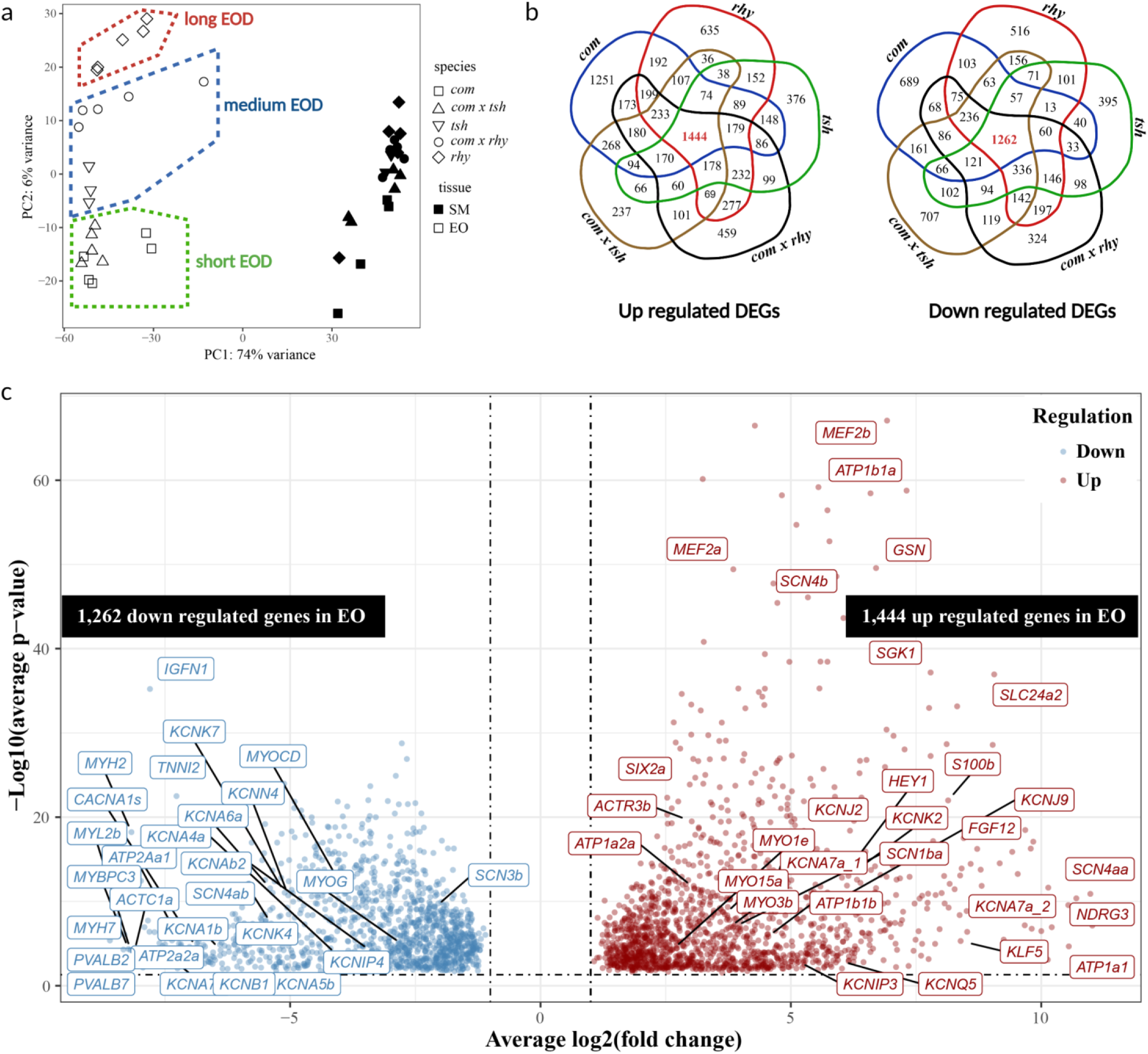
Differential gene expression between electric organ (EO) and skeletal muscle (SM) in *C. compressirostris* (*com*), *C. rhynchophorus* (*rhy*), *C. tshokwe* (*tsh*) and hybrids *C. compressirostris* ♂ × *C. rhynchophorus* ♀ (*com* × *rhy*), *C. compressirostris* ♂ × *C. tshokwe* ♀ (*com* × *tsh*). **a** Principal component analysis (PCA) of gene expression levels between EO and SM in 5 species/hybrids. **b** Venn Diagram graph for up (left) and down (right) regulated genes shared in 5 species/hybrids. All differentially expressed genes (DEGs) have **|**log2(fold change)**|** >1 and a p-value < 0.05. Many of the DEGs are related to “membrane” and “plasma membrane” (see Supplementary file 3). **c** Volcano plot showing genes differentially expressed in EO (relative to SM) in all 5 species/hybrids. X-axis is the average log_2_(fold change) among 5 species/hybrids, and y-axis is the associated –log_10_ (average p-value for 5 species/hybrids). Potential candidate genes and genes with low p-value or high fold change are labeled with their name.

### Genes with EO-specific expression pattern

Differential gene expression analysis was used for pairwise comparisons between EO and SM for each species and hybrid (Fig. 2b). We identified significantly differentially expressed genes (DEGs) based on a |log_2_ folder change (log_2_FC)|> 1 and a p-value < 0.05. We specifically identified genes with an EO-specific expression pattern shared among all *Campylomormyrus* species/hybrids. There were 1,444 up-regulated and 1,262 down-regulated DEGs that were shared in the comparison of EO and SM in all species/hybrids (Fig. 3b, c).

Among the DEGs up-regulated in the EO, 54 genes were related to transmembrane ion transport (Fig. 3c, Table 1, Supplementary Table 1). We identified four genes encoding sodium/potassium-ATPase α and β subunit (*ATP1a1*, *ATP1a2a*, *ATP1b1a* and *ATP1b1b*), and three *Na_v_* channel genes (*SCN4aa*, *SCN4b* and *SCN1ba*). Several genes encoding for different types of potassium channels were also identified: four *K_v_* channel genes (*KCNA7a_1*, *KCNA7a_2*, *KCNIP3* and *KCNQ5*), two *K_ir_* channel genes (*KCNJ2* and *KCNJ9*), and one *K_2p_* channel gene (*KCNK2*). Further transmembrane ion transport DEGs were chloride, calcium and other cation channel genes (Supplementary Table 1). Several solute carrier family genes were also up-regulated in the EO, in particular *SLC24a2* (Table 1).

**Table 1.**
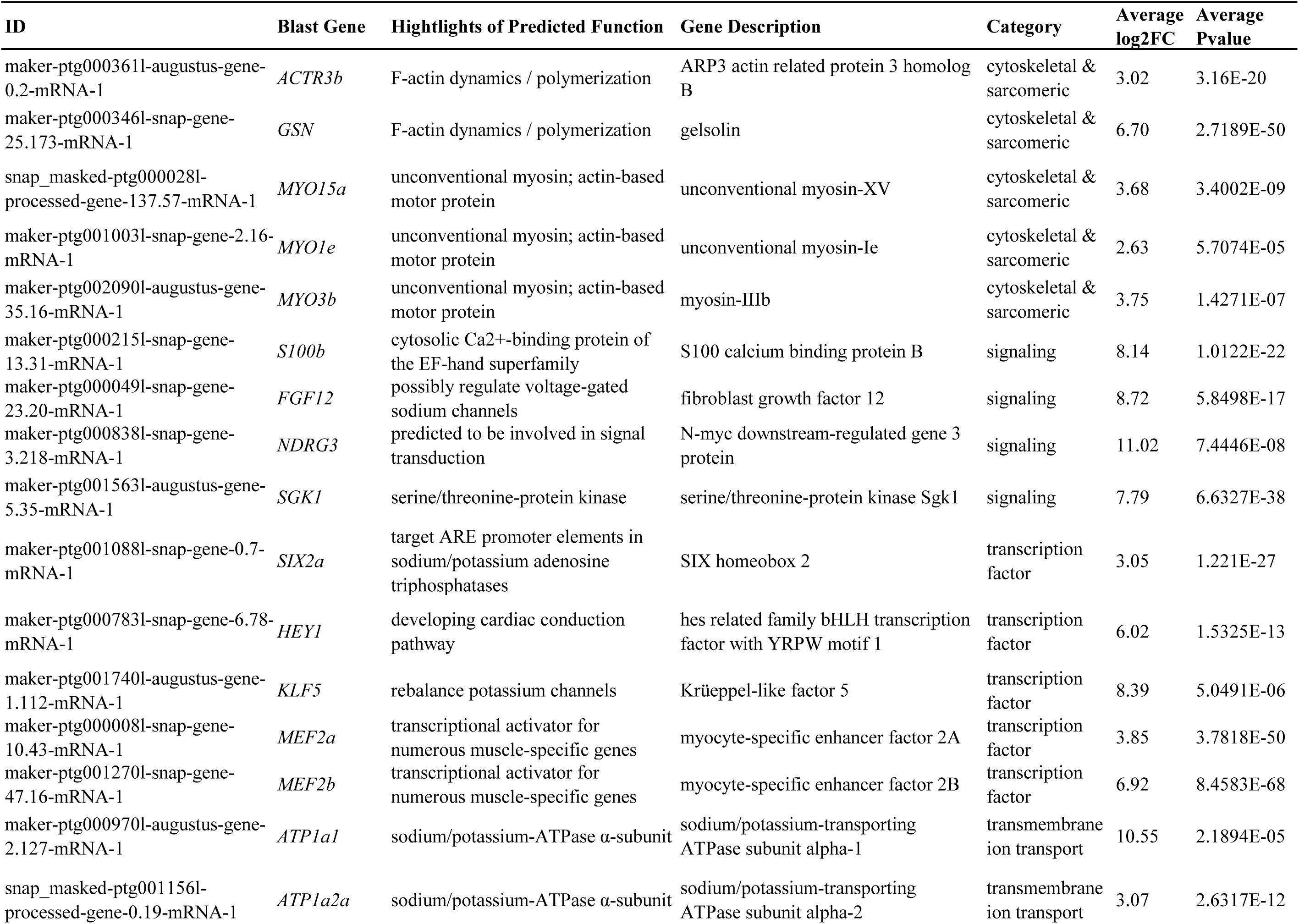
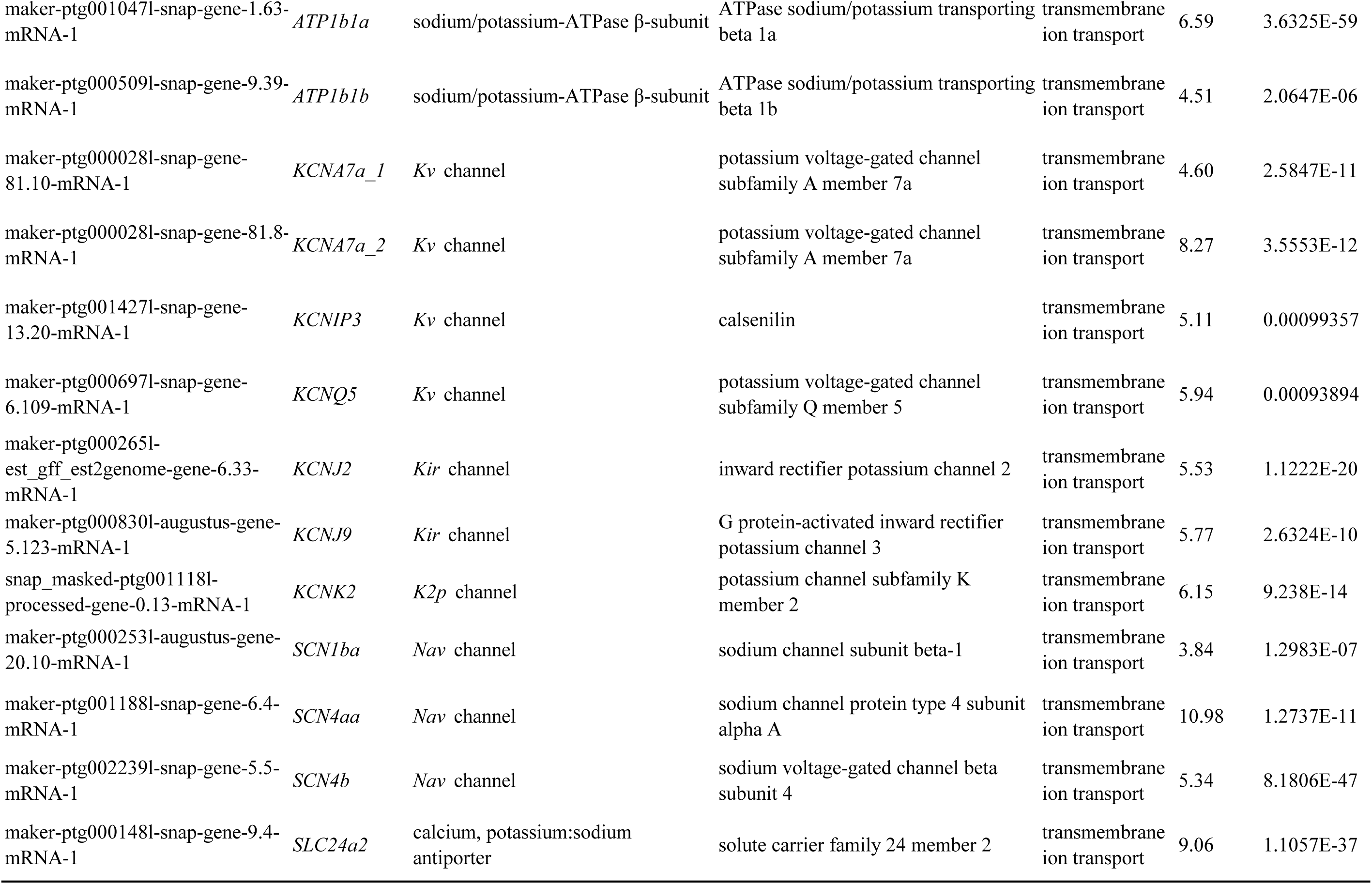
Candidate genes up-regulated in all species/hybrids in the electric organ relative to sceletal muscle.

18 genes up-regulated in the EO were associated with cytoskeletal and sarcomeric protein (Supplementary Table 1). The predicted function of those genes were mainly related to F-actin dynamics and unconventional myosin activity (Table 1). A signaling gene *NDRG3* showed very high overexpression in EO (log_2_FC=11.02), as well as the genes *SGK2, S100b* and *FGF12*. The up-regulated transcription factors in the EO included *KLF5*, *FOXL2*, *SIX2a*, *HEY1* and two myocyte-specific enhancer factors (*MEF2a* and *MEF2b*).

In the down-regulated DEGs in EO (or up-regulated in SM), 44 genes were classified into the category “cytoskeletal & sarcomeric” (Fig. 3c, Supplementary Table 2). There were 37 transmembrane ion transport genes down-regulated in EO, which were related to the ions potassium, sodium, and calcium. In contrast to the expression pattern of the two *KCNA7a* copies, five *K_v_1* subfamily genes (*KCNA1b*, *KCNA4a*, *KCNA5b*, *KCNA6a* and *KCNA7b*) were down- regulated in the EO. This was also the case for other potassium and sodium channel genes, e.g. *K_v_* subfamily genes (*KCNB1*, *KCNE4*, *KCNIP4*), *K_2p_* subfamily genes (*KCNK4*, *KCNK7*), a *K_ca_* subfamily gene (*KCNN4*), and *Na_v_* channel genes (*SCN3b*, *SCN4ab*). Two muscle-specific transcription factors, *MYOCD* and *MYOG*, were also down-regulated in EO (Supplementary Table 2).

We applied a Gene Ontology (GO) enrichment analysis to further examine the function of all the up- and down-regulated DEGs in EO respectively (Dennis et al. 2003). Among the up-regulated DEGs in the EO, there were 44 significantly enriched GO terms (Fisher’s exact test p-value<0.01, Supplementary Fig. 1, Supplementary Table 3). Among them, the three GO terms with the highest number of DEGs were all related to the cell membrane: membrane (464 DEGs), integral component of membrane (309 DEGs) and plasma membrane (237 DEGs). There were 47 DEGs assigned to the enriched GO term “ion transport”. 62 and 35 DEGs were assigned to the enriched Golgi-related GO terms “Golgi membrane” and “Golgi apparatus”, respectively. In addition, there were 23 DEGs assigned to the enriched GO term “actin filament binding”. There were 73 GO terms significantly enriched for DEGs down-regulated in the EO (up-regulated in SM, Supplementary Fig. 2, Supplementary Table 4). They were associated with skeletal and cardiac muscle tissue related GO terms.

### Genes with expression levels related to EOD duration

The PCA plot from transcriptome-wide gene expression showed a significant association between overall gene expression and EOD duration in all species/hybrids (PC2 in Fig. 3a; accounting for 6% of the variance in gene expression). DESeq2 provides a Likelihood Ratio Test (LRT) that compares how well a gene’s read count data fit a “full model” (with independent variables) compared to a “reduced model” (without those variables). Therefore, it is well suited to explore whether there are any significant associations of gene expression levels across a series of values of an independent variable (here, EOD duration) (Love et al. 2015). Specifically, we used this approach to test whether a gene’s expression fits a pattern of increasing or decreasing over the different durations in two different tissues, EO and SM (Bendjilali et al. 2017). In order to avoid any bias potentially stemming from distorted expression pattern in the hybrids, we only used the quantification data from the parental pure-bred (F0) species. The LRT analysis returned 1,874 significant genes using a threshold of padj < 0.05. Those genes were further sorted into groups using the degPatterns function. Each such group contained genes following a specific pattern of expression across the different duration values in the analyzed tissues EO and SM (L Pantano 2019).

The degPatterns function generated 27 groups of different expression pattern in EO and SM, relative to EOD duration (Supplementary Fig. 3). To identify EOD duration-specific genes, we focused on the groups meeting the following criteria: 1) the gene expression level in EO is higher than SM in all F0 species (i.e., the gene is consistently up-regulated in the EO); 2) the gene expression level in the EO shows an increasing or decreasing pattern, relative to EOD duration. Two groups showed a consistent increasing expression pattern (groups 5, 41 genes; and 6, 239 genes) and one a decreasing expression pattern (group 3, 405 genes), relative to the EOD duration (Fig. 4).

**Fig. 4.**
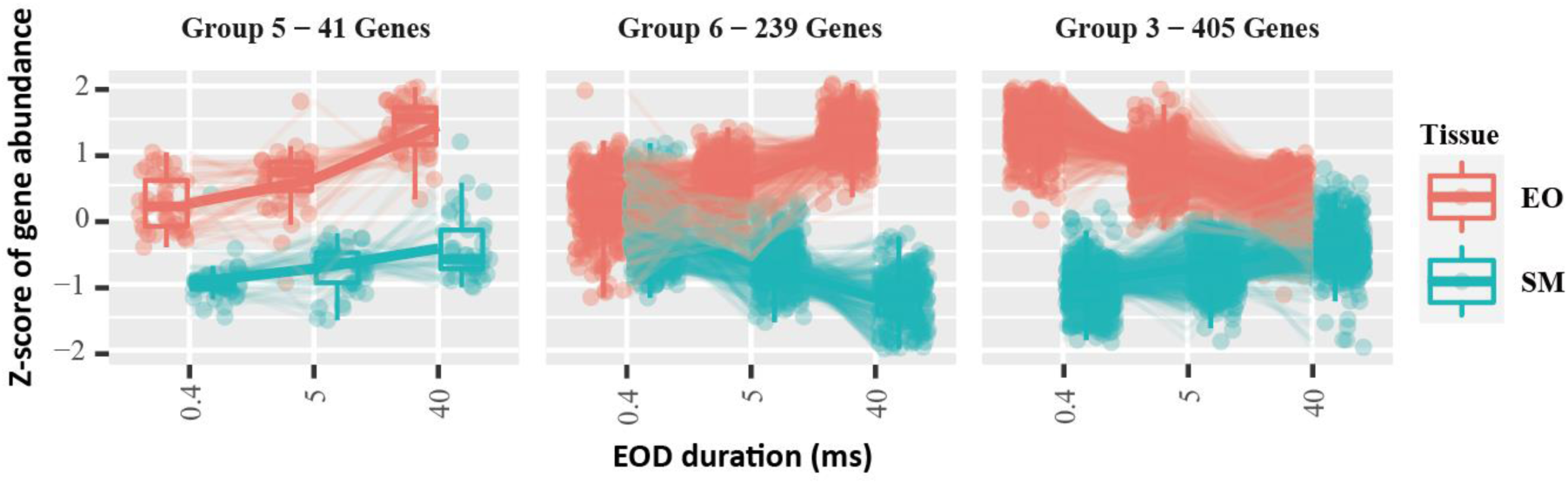
RNA-seq data clustered by EOD duration (only for the 3 pure-bred species). Increasing (**Group 5** and **6**) and decreasing (**Group 3**) expression patterns over EOD duration among electric organ (EO) and skeletal muscle (SM). The x-axis for each group represents the duration of the analyzed species: *C. compressirostris* (0.4ms), *C. tshokwe* (5ms) and *C. rhynchophorus* (40ms).

In the increased expression pattern groups (5 and 6), we found *K_ir_* subfamily gene *KCNJ2* and the transcription factor Krüppel-like fator 5 (*KLF5*), both were found among the genes with EO-specific expression as well (Table 2). In the decreased expression group 3, there were two transmembrane ion transport genes (*KCNK6* and *KCNQ5*) and two cytoskeletal and sarcomeric genes (*ACTR3b* and *NHS*, Table 2).

**Table 2.** Genes with expression correlated to EOD duration.

| Group | Gene ID in annotation | Gene | Highlights of Predicted Function | Gene Description | Category |
| --- | --- | --- | --- | --- | --- |
| 3 | maker-ptg000361l-augustus-gene-0.2-mRNA-1 | *ACTR3b* | F-actin dynamics / polymerization | ARP3 actin related protein 3 homolog B | cytoskeletal & sarcomeric |
| 3 | maker-ptg000509l-snap-gene-4.4-mRNA-1 | *NHS* | regulator of actin remodelling | Nance-Horan syndrome protein | cytoskeletal & sarcomeric |
| 3 | maker-ptg001966l-augustus-gene-18.42-mRNA-1 | *KCNK6* | outward rectification in a physiological potassium gradient and mild inward rectification in symmetrical potassium conditions | potassium channel subfamily K member 6 | transmembrane ion transport |
| 3 | maker-ptg000697l-snap-gene-6.109-mRNA-1 | *KCNQ5* | voltage-gated potassium channel | potassium voltage-gated channel subfamily Q member 5 | transmembrane ion transport |
| 5 | maker-ptg000265l-est_gff_est2genome-gene-6.33-mRNA-1 | *KCNJ2* | inwardly rectifying potassium channel | inward rectifier potassium channel 2 | transmembrane transport |
| 5 | maker-ptg001740l-augustus-gene-1.112-mRNA-1 | *KLF5* | transcription factor which might regulates potassium channel genes | krueppel-like factor 5 | transcription factor |

Assigning DEGs with increasing expression pattern to GO terms revealed 19 significantly enriched (Fisher’s exact p-value < 0.05) GO terms (Supplementary Fig. 4, Supplementary Table 5). 18 genes were assigned to the enriched GO term “Golgi apparatus”, 13 to “ion transport”. Among the genes with decreasing expression pattern (group 3), 41 GO terms were significantly enriched (Supplementary Fig. 5, Supplementary Table 6). 12 of the genes were assigned to the GO term “axon guidance” which yielded the lowest p-value. There were also several enriched terms which might be functionally related to the EOD, e.g. membrane, Golgi membrane and apparatus, calcium ion binding, and ATP binding.

### Allele specific expression in F1 hybrids

Two cohorts of F1 hybrids with one short duration EOD (*com* × *tsh*) and one medium duration EOD (*com* × *rhy*) were analyzed in our study. In total, we identified fixed SNPs (homozygous in parental species) in 177 genes differentially expressed in EO and in 77 differentially expressed in SM in the hybrid *com* × *rhy*. For the hybrid *com* × *tsh*, the respective SNP numbers were 52 in genes differentially expressed in the EO and 36 in genes differentially expressed in SM (Fig. 5a). For each of these genes, we calculated the allelic read proportion of the allele stemming from the parental species *com* (as identified by the fixed SNPs), averaged over the specimens of the respective hybrid cohort. In general, most genes exhibit an equal expression of both parental alleles, with more genes have a *com* proportion near 0.5 (Fig. 5a). Among the genes with differentially expressed alleles, alleles stemming from *com* had an overall tendency towards higher expression, compared to the alleles from *rhy* or *tsh*, in both EO and SM from two hybrid cohorts (Fig. 5a).

**Fig. 5.**
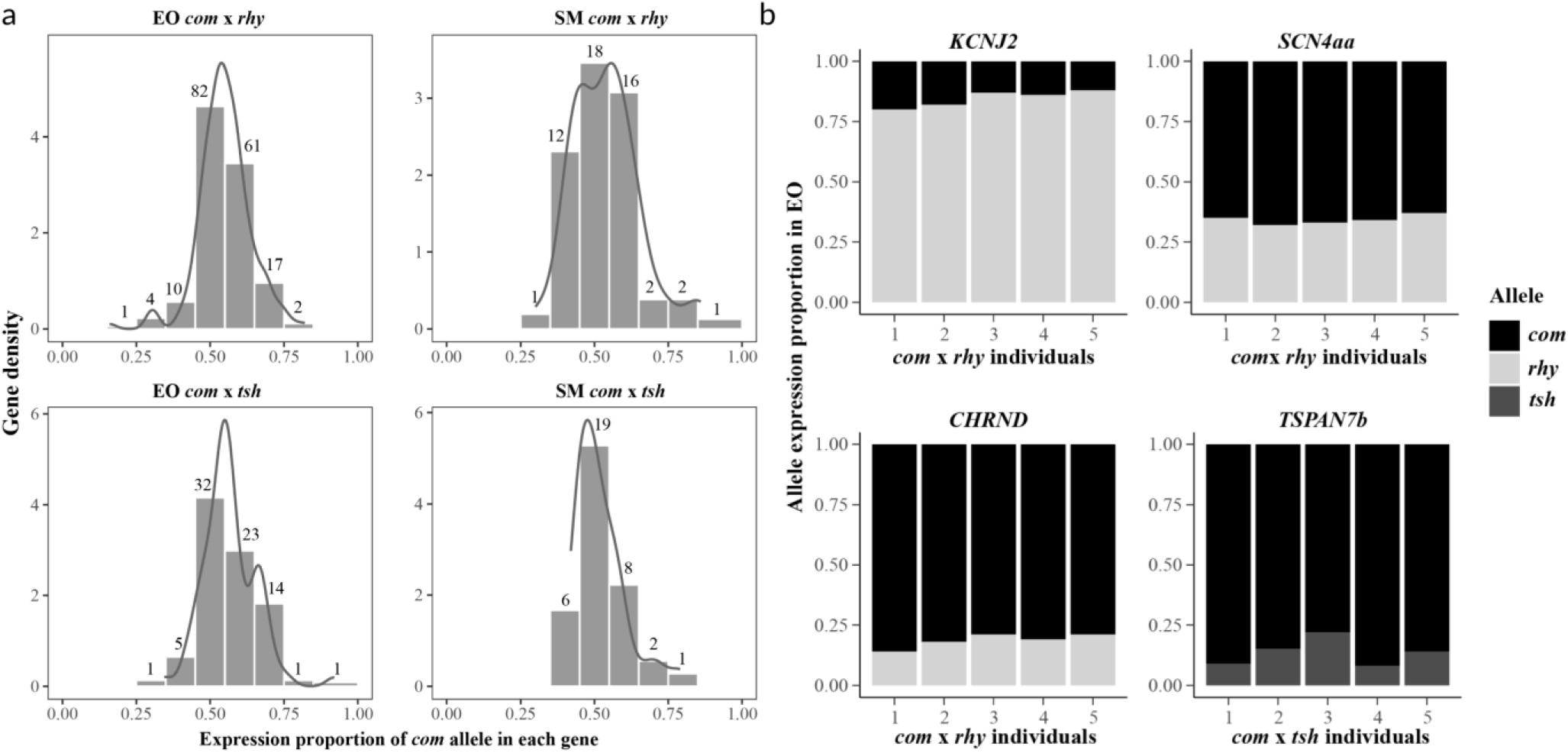
Allele specific expression in electric organ (EO) and skeletal muscle (SM) among two hybrid cohorts *C. compressirostris* (*com*) × *C. rhynchophorus* (*rhy*) and *C. compressirostris* (*com*) × *C. tshokwe* (*tsh*). **a** Gene density (y-axis) from two different tissues in each hybrids. The x-axis shows the expression proportion of the allele stemming from one parental species (*com*). Numbers above the bars represent the number of genes in the respective proportion ranges. **b** Proportion of two alleles from parental species in four genes related to ion transport and membrane (*KCNJ2*, *SCN4AA*, *CHRND* and *TSPAN7b*) in the EO. The x-axis represents the different individual samples of the corresponding hybrid cohort.

In order to understand the allelic expression imbalance (AEI), we counted the number of genes with more than 0.6 proportion of one parental allele proportion for all individuals in each analyzed hybrid set (Chang et al. 2002). In total, we identified 17 and 7 genes with AEI in EO and SM of the hybrid *com* × *rhy*, respectively; 2 and 1 such genes were identified in EO and SM of the hybrid *com* × *tsh*, respectively (Supplementary Table 7). In all the genes with AEI, the allele from *com* showed a higher expression proportion, except for the gene *KCNJ2* in the hybrid *com* × *rhy* where allele expression was biased in the opposite direction (average proportion of *com* allele was 0.16, Fig. 5b). We inferred amino acid sequences from the transcript sequences of the *KNCJ2* gene from *com*, *tsh* and *rhy*. The inferred protein sequence between *com* and *tsh* were identical, but *rhy* showed two amino acid substitutions at sites 60 (corresponding to the fixed SNP we identified in hybrids) and 198 (Supplementary Table 8). The amino acid substitution at site 60 was considered benign in the Polyphen2 analysis (Adzhubei et al. 2010), while the substitution at site 198 may have changed the protein function in *rhy* (inferred as probably damaging; Supplementary Table 9).

We also identified AEI in the gene *SCN4aa* in the EO of *com* × *rhy* hybrids (average proportion of *com* allele was 0.66, Fig. 5b). This proportion of the *com* allele is much higher than in the hybrid *com* × *tsh*, where the *com* allele of the *SCN4aa* only had a proportion of 0.46 in the EO. In the EO of hybrid *com* × *rhy*, AEI was identified in the gene *CHRND*, which might relate to ion channel gating (Fig. 5b). The *TSPAN6b* gene, encoding for an integral component of the plasma membrane, was also identified to exhibit a significant AEI in the EO of hybrid *com* × *tsh* (average proportion of *com* allele was 0.86, Fig. 5b).

## Discussion

### Convergent gene expression in different electric fish lineages

The myogenic EO has convergently evolved six times in fishes. Even though the EOs show great differences in electrocyte morphology among independently evolved electric fish lineages, particular genes exhibit similar transcriptional expression patterns in the EO, relative to skeletal muscle (Gallant et al. 2014).

Several EO-specific candidate genes that we identified in *Campylomormyrus* were also overexpressed in the EO of other electric fish lineages, possibly indicating convergent expression pattern evolution in electric fish. This is particularly apparent in genes related to sodium and potassium currents. For instance, the *Na_v_* channel gene *SCN4aa*, considered to be very important in regulating the sodium current to electrocytes, has been previously found overexpressed in the EO of different electric fish lineages, i.e., Siluriformes, Gymnotiformes, and Mormyroidea other than *Campylomormyrus* (Wang and Yang 2021). The *FGF13a* that regulates this channel was consistently overexpressed in those electric fish species as well (Gallant et al. 2014). Interestingly, we identified another up-regulated ortholog (*FGF12*) in the EO which may have a similar function. In addition, multiple isoforms of sodium/potassium ATPase α and β subunits and several transcription factors (*SIX2a*, *HEY1*) were found to be convergently up-regulated in the EO among these electric fish lineages (Gallant et al. 2014), a pattern confirmed for *Campylomormyrus* in our study.

Overexpression of another transcription factor (*MEF2a*) and of the calcium binding gene *S100b* is characteristic for mormyrid EOs, i.e., *Paramormyrops*, *Brienomyrus* (Gallant et al. 2012; Gallant et al. 2017), and *Campylomormyrus* (this study). We recently found *KCNA7a* to be tandemly duplicated in *Campylomormyrus* (Cheng et al. 2023) and *Paramormyrops* (by re- analysis of the genome provided in Gallant et al. 2017). This tandem duplication might be exclusive to mormyrid fishes, as we did not find it in available genomes neither of other electric fishes nor in *Scleropages* (a non-electric fish closely related to mormyrids; data not shown). In our study, both gene copies *KCNA7a_1* and *KCNA7a_2* were consistently up-regulated in the EO (*KCNA7a_2* showed even higher expression than *KCNA7a_1*). *KCNA7a* was inferred to be under positive selection in the transmembrane helices 3-4 linker, and is considered to relate to the differences in EOD duration among *Brienomyrus* and *Gymnarchus* (Swapna et al. 2018).

The *NDRG3* gene (N-Myc Downstream-Regulated Gene 3) exhibited a remarkable overexpression in the EO of *Campylomormyrus*. Interestingly, the phosphopeptides encoded by an ortholog (*NDRG4*) were highly enriched in the EO of the strongly discharging gymnotiform electric eel (*Electrophorus electricus*) (Traeger et al. 2017), indicating high expression level of this gene. In addition, NDRG4 has been identified in zebrafish as a novel neuronal factor essential for sodium channel clustering at the nodes of Ranvier, the only places where action potentials are regenerated (Fontenas et al. 2016). The function of *NDRG3* in the nervous system has rarely been investigated. The NDRG3 protein can interact with extracellular signal-regulated kinases (ERK1/2) (Lee et al. 2015), which regulate *K_v_4.2* in the dendrites of hippocampal CA1 pyramidal neurons (Schrader et al. 2006; Gupte et al. 2016) as well as the *Na_v_1.7* channel (Stamboulian et al. 2010). In porcine as well as human lens, ERK1/2 is activated by the TRPV1 ion channel (Mandal et al. 2019), which was also overexpressed in EOs in *Campylomormyrus* (Supplementary Table 1).

### Gene expression specificity in *Campylomormyrus*

The EO in mormyrids is derived from myogenic tissue, which transitions from a motoric/sarcomeric organization of muscle fibers to a continuous tube of electrocytes parallel to the spinal cord (Denizot et al. 1982). This transition process during the ontogeny of the EO involves cell size, morphology and physiology, and is still only partially understood. Some genes encoding for sarcomeric proteins, e.g. troponin I isoforms, myosin heavy chain and tropomyosin, are overexpressed in the EO of the mormyrid *Brienomyrus brachyistius* (Gallant et al. 2012), providing a preliminary insight into the developmental transition from SM to EO. In the EO of *Campylomormyrus*, however, we rarely found those genes up-regulated. Instead, the up- regulated actin-related genes in *Campylomormyrus* were more related to F-actin dynamics and included several unconventional myosins (Table 1, Supplementary Table 1). The four paralogous transcription factors *MEF2a-d* are responsible for the transcriptional activation of muscle- specific genes in the early specification of skeletal muscle (Black and Olson 1998). Whereas *MEF2a* was overexpressed in the EOs of both *Brienomyrus* (Gallant et al. 2012) and *Campylomoryrus*, a further paralog *MEF2b* was overexpressed only in *Campylomormyrus* (our study). The difference in expression of F-actin-related/sarcomeric proteins and *MEF2* transcription factors between two mormyrids genera suggests that the developmental transition in the EO might be different or, in other words, that the organization of the F-actin system in electrocytes may vary across mormyrids. It has to be analyzed further whether these differences in the organization of the F-actin cytoskeleton concern the sarcomeric structure, the stalks of electrocytes, or both.

In addition, two paralogs of inwardly rectifying potassium channel (*K_ir_*) genes *KCNJ2* and *KCNJ9* were overexpressed in EOs of *Campylomormyrus*, along with *KCNQ5* and *KCNK2*. The mechanisms regulating potassium channels expression in electric fish are still unknown. We identified one transcription factor, Krüppel-like factor 5 (*KLF5*), that showed a high overexpression in EOs. In *Drosophila*, Krüppel is involved in the regulation of potassium channel expression. In case of a loss of *Shal* (*KCND*) potassium channel in *Drosophila*, Krüppel expression is induced and up-regulates expression of *Shaker* (*KCNA*) and *slowpoke* (*K_ca_*) potassium channels (Parrish et al. 2014). Remarkably, *Shal* (*KCND*) potassium channel is also not expressed in our studied *Campylomormyrus* species/hybrids (Parrish et al. 2014). We thus suppose that the EO differs from SM by the expression of a unique set of potassium channels that may contribute to the shape of the EÓs action potential and thus the shape of the EOD signal. Moreover, we propose the *KLF5* gene to represent a transcription factor that drives the expression of regulating potassium channels in EO.

Our study has further revealed that different paralogs from the solute carrier family are active in EOs. Solute carriers form a group of membrane transport proteins located in various cellular membrane systems which transport diverse substrates including amino acids, oligopeptides, inorganic cations and anions (He et al. 2009). We have found several genes of this gene family overexpressed in the EO that transport inorganic cations and anions, e.g. sodium, calcium, chloride. Especially the gene *SLC24a2* was highly overexpressed in *Campylomormyrus* EOs. This is a calcium/cation antiporter localized in the plasma membrane that mediates the extrusion of one calcium ion and one potassium ion in exchange for four sodium ions (Wang et al. 2017). Overexpression of a calcium-extruding transporter in the EO indicates that regulation of the cytosolic calcium level in electrocyte differs from that in SM. Unfortunately, we have no information yet on the distribution the *SLC24a2* protein within the electrocytes, and whether this calcium transporter is confined to a distinct region of the cell to mediate local regulation of the calcium level.

### Differential gene expression with respect to EOD duration divergence among *Campylomormyrus* species

*Campylomormyrus* species produce species-specific EODs; their duration varies in a 100-fold range across species. The EOD is assumed to be mediated by sodium and potassium currents across the plasma membrane (Stoddard and Markham 2008). The depolarizing phase of an action potential is primarily produced by sodium influx. The repolarization phase is - along with a gradual decreasing sodium influx - affected by the orchestrated activities of delayed rectifier and inward rectifier potassium channels (Nass et al. 2008; Stoddard and Markham 2008). We suppose that species producing EODs of different duration may be equipped with different channel types or channel orthologs with different properties. However, certainly other mechanisms, such as different cell morphology, may also contribute to the EOD duration diversification.

The PCA from RNA-seq data showed a clear association between the overall gene expression and the EOD duration pattern (Fig. 3a). Based on the preliminary PCA and LRT result, we identified several genes which might contribute to EOD duration diversification in *Campylomormyrus*, including the potassium channel genes (*KCNJ2*, *KCNK6* and *KCNQ5*), actin-related genes (*ACTR3b*, *NHS*), and transcription factor *KLF5* (Table 2).

The gene *KCNK5* was found to be up-regulated in *Paramormyrops* (producing a short EOD) compared to the species with an elongated EOD (Losilla et al. 2020). In *Campylomormyrus*, the expression of another paralog *KCNK6* was also higher in species with short EOD. Two-pore potassium channels (*K_2p_*) usually generate an outward potassium current and are also known as potassium leak channels. When silencing the *KCNK6* gene in the human heart, the action potential duration is prolongated (Chai et al. 2017). Another voltage-gated potassium channel gene *KCNQ5* was decreasingly expressed in elongated EOD *Campylomormyrus* species. It forms M-type potassium current, a slowly activating and deactivating potassium conductance that works in determining the subthreshold electrical excitability of neurons (Hibino et al. 2010). The lower expression of both potassium channels genes in elongated EOD species will probably decrease the outward potassium current and consequently prolongate EOD repolarization.

The gene *KCNJ2* was increasingly expressed in elongated EOD species. It encodes for an inwardly rectifying potassium channel, with the greater tendency to allow potassium ions to flow into a cell rather than out of a cell (Hibino et al. 2010). The inward potassium current stabilizes the resting membrane potential of the cell and modulates the cardiac repolarization processes (Hibino et al. 2010; Li et al. 2017). This inward rectifier channel-mediated potassium current is responsible for shaping the initial depolarization and final repolarization of the action potential in human cardiomyocytes (Dhamoon and Jalife 2005; Jeevaratnam et al. 2018).

Regarding allele specific expression, there was a tendency towards higher expression of *com* alleles in the EOs and SMs of two analyzed hybrid cohorts (Fig. 5a). However, the phenotype of the EOD waveform in each hybrid is closer to the other parental species. This points towards some genes playing key roles in regulating the EOD waveform in the hybrids. The gene *KCNJ2* showed allelic expression imbalance (AEI) in *com* × *rhy* hybrids, which was the only gene with AEI and for which the *rhy* allele was preferentially expressed (Fig. 5b). The EOD in the adult hybrids *com* × *rhy* was of intermediate duration (4 ms), and the shape and waveform resemble the subadults’ EOD in *rhy*. Both the EOD phenotype and the AEI in *KCNJ2* were hence closer to the parental species with the elongated EOD, i.e., *rhy*. The expression of *KCNJ2* in the EO among the purebred species also increased with increasing EOD duration, e.g. the expression in *rhy* is higher than in *com*. This suggests that the *KCNJ2* gene might be under cis-regulation, and it should be a powerful candidate gene involved in the regulation of EOD duration in *Campylomormyrus*.

In addition, the *KCNJ2* gene in the species *rhy* (very long EOD) exhibits two non-synonymous substitutions, one of which predicted to cause a functionally relevant to amino acid substitution (at site 198; Supplementary Table 8, 9). Interestingly, the same substitution at site 198 is present in another species with very long EOD (*C. numenius*, EOD duration 40 ms), while it is absent in other *Campylomormyrus* species with short or medium EOD which resemble the amino acid sequence of *com* and *tsh* (Cheri, Cheng & Tiedemann, unpubl. results). *C. numenius* and *rhy* are phylogenetically close (Lamanna et al. 2016), such that the shared amino acid substitution could also reflect phylogenetic affinity. Nonetheless, the found amino acid substitution with inferred functional relevance could relate to the evolution of very long EODs in *Campylomormyrus*. Then, the *KCNJ2* gene could modulate EOD duration by a combination of expression level and functional protein sequence alteration. In summary, this study identifies the *KCNJ2*, *KCNK6* and *KCNQ5* genes, possibly in combination with other genes (e.g. *KLF5, ACTR3b, NHS*) as strong candidates underlying EOD duration diversification in the weakly electric fish genus *Campylomormyrus*. The diverged EOD likely affect the food spectrum and are used for mate recognition. This potential dual function in disruptive natural selection and pre-zygotic reproductive isolation would rank the EOD as a “magic trait”^14^, which may have promoted the ecological (probably sympatric) speciation and radiation of *Campylomormyrus* in the Congo River.

## Methods

### Animals, RNA isolation, library preparation and sequencing

Three adult specimens of *C. tshokwe* were collected at Brazzaville/Republic of in the Congo River in 2012 and stored in RNAlater in -80°C. Five adult specimens from each of the other two species (*C. compressirostris*, *C. rhynchophorus*) and two hybrids (*C. compressirostris* ♂ × *C. rhynchophorus* ♀, *C. compressirostris* ♂ × *C. tshokwe* ♀) were artificially bred and raised at the University of Potsdam. All specimens except for *C. tshokwe* were anesthetized by a lethal dose of clove oil, and dissected on cold 99% ethanol. Electric organ (EO) and skeletal muscle (SM) tissue from each specimen were flash frozen in liquid nitrogen, and further preserved in -80°C. In total, we collected three samples of both EOs & SMs from *C. tshokwe*, five samples of both EOs & SMs from the other four species/hybrid cohorts in this study.

The RNA isolation was performed in all the EOs and SMs samples using QIAGEN RNeasy Fibrous Tissue Mini Kit. Total RNA concentration was estimated using a NanoDrop 1000 spectrophotometer (ThermoFischer Scientific, Germany), RNA quality was checked with an Agilent Bioanalyzer 2100 (Agilent Technologies, USA). mRNA enrichment was performed by poly (A) capture from isolated RNA using NEXTflex Poly (A) Beads. Strand-specific transcriptomic libraries were built using NEXTflex Rapid Directional RNA-Seq Kit (Bioo Scientific, USA) based on the manufacturer’s instructions.

Libraries were sequenced as 150 bp paired-end reads by Illumina HiSeq 4000 sequencing system at a commercial company (Novogene). Raw reads have been deposited in the National Center for Biotechnology Information (NCBI) Gene Expression Omnibus (accessions number:GSE240783). We trimmed the adapter sequences and low quality reads using a 4 bp sliding window with a mean quality threshold of 25, and a minimum read length of 36 bp by Trimmomatic v0.39 (Bolger et al. 2014). Read quality, before and after read filtering, was measured by FastQC v0.11.9 (Andrew 2010).

### Differential gene expression analysis

The quality-filtered reads from EOs and SMs were mapped to the *C. compressirostris* genome (Cheng et al. 2023) using RSEM (Li and Dewey 2011) for gene level-quantification estimation. The estimated counts were imported into R/Bioconductor with the tximport package, which produced count matrices from gene-level quantification by taking the effective gene length into account (Paraskevopoulou et al. 2020). Low count (≤ 10) and low frequency (not present in at least two replicates) genes were removed. We performed a principle component analysis (PCA) from filtered and log-transformed counts. One SM sample from *C. compressirostris* was removed from this study, as its overall gene expression showed a deviant unusual pattern in the PCA.

We forwarded the normalized count matrices to DESeq2 (M. I. Love et al. 2014) to infer expression differences among EO and SM in each species/hybrid cohort respectively. We used a false discovery rate threshold of 0.05 to correct for multiple testing. The differentially expressed genes (DEGs) were identified with |log_2_ folder change (log_2_FC)| > 1 and p-value < 0.05. In order to detect the EO specific gene expression pattern, we used Venn Diagrams (Chen and Boutros 2011) to visualize the shared DEGs (up- and down-regulated separately) among three purebred species and two hybrid cohorts.

The shared DEGs were annotated against the NCBI *nr* database by blastx with an e-value cutoff 1e^-10^. In addition, the up- and down-regulated DEGs in the EO were used to perform a Gene Ontology (GO) enrichment analysis (Dennis et al. 2003).

### RNA-seq data clustering by EOD duration

The PCA plotting from log-transformed count matrices showed a clear pattern by the length of EOD (Fig. 3a). To identify genes with an expression pattern associated to EOD duration, we used DESeq2 to perform a likelihood ratio test (Michael I. Love et al. 2014) (LRT in the DESeq2 package). This test compares how well a gene’s count data fit a “full model” compared to a “reduced model” (Bendjilali et al. 2017). Our full model was an equation: full = ∼ duration * tissue. The duration is the length of the EOD in each purebred species, and tissue is the type of sample (EO or SM). The reduced model excluded the interaction between duration and tissue: reduced = ∼ duration + tissue. Genes with adjust P- value (padj) < 0.05 were considered to fit the “full model”. We used the degPatterns function from the ‘DEGreport’ package to cluster different groups with particular expression pattern using those significant genes across samples (L Pantano 2019), with time = “duration”, col = “tissue”.

The generated groups of different gene expression pattern across EOD duration were analyzed to identify genes with an expression pattern association with EOD duration. We hence focused on those groups fulfilling the following two criteria: 1) the gene expression level in EO is higher than SM in all F0 species; 2) the gene expression level in EO across EOD duration showed a consistent increasing or decreasing pattern.

The identified genes with increased and decreased expression relative to EOD duration were blasted against *nr* database using an e-value cutoff 1e^-10^. In addition, a GO term analysis was also performed for these genes.

### Allelic specific expression analysis

The F1 hybrid contains two sets of subgenome from two parental species. Examination of allele specific expression can be applied to detect the allelic imbalance in transcription in heterozygous F1 hybrids. We only focused on transcripts of genes with biallelic SNPs fixed among the respective F0 parental species (hence, heterozygous only in F1 hybrids, homozygous in parental species).

We mapped the trimmed and filtered RNA-seq from five species/hybrids (in EOs and SMs, respectively) to the *C. compressirostris* genome using STAR v2.7.7 (Dobin and Gingeras 2015). The generated bam files were sorted according to the coordinates by SAMtools v1.15 (Danecek et al. 2021). Variant calling was performed by BCFtools v1.9 (Danecek et al. 2021) in EOs and SMs respectively, using the command “bcftools mpileup –f REFERENCE LIST_OF_BAM –Ou | bcftools call –mv –Ob –o BCFFILE”, where the REFERENCE, LIST_OF_BAM, BCFFILE were the CDS sequence name of the *C. compressirostris* genome, the list of bam files, and the output bcf file name, respectively.

After the variant calling, we performed the following steps to identify the fixed parental biallelic SNPs for each hybrid set. Firstly, we excluded the uncalled variants and only preserved biallelic SNPs using the command “bcftools view --exclude-uncalled -m2 -M2 BCFFILE > CALLING_AD”, where the CALLING_AD was the allelic depth for the final biallelic SNPs. Secondly, we discarded SNPs where the variant calling score at QUAL field was lower than 70, and allelic depth was lower than 10 in both alleles. Finally, we obtained high quality SNPs, at which both parental species were homozygous and fixed for a different allele.

For each hybrid, we calculated the expression proportion of the allele from *C. compressirostris* in EO and SM, respectively. Genes with *C. compressirostris* allele proportions <0.4 or >0.6 in the transcriptomes were considered exhibiting an imbalanced expression^31^.

## Data availability

Sequence data have been deposited at NCBI Gene Expression Omnibus under accession GSE240783.

## Acknowledgements

We thank Dr. Linh Nguyen for fish breeding and raising. This project is funded by University of Potsdam.

## Author contributions

R.T. conceived and supervised this study, and provided finicial and logistical support. F.C. performed lab work, analysis and manuscript writing with the input from R.T., D.A.B., B.O., K.F., and S.A.S..

