## Supplementary Material for "Gene and allele specific expression underlying the electric signal divergence in African weakly electric fish"

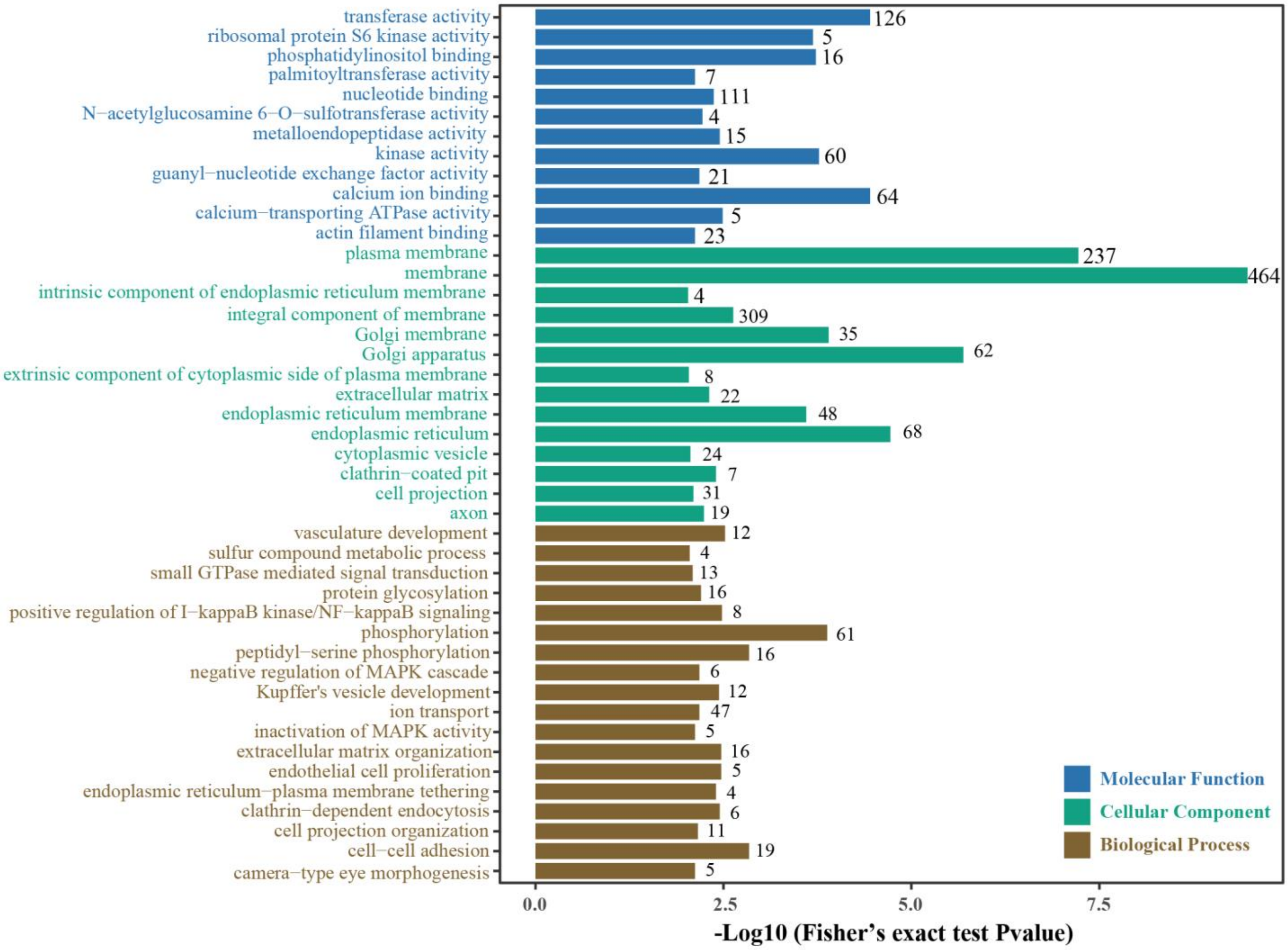

**Supplementary Fig. 1**

44 significantly enriched Gene Ontology (GO) terms with Fisher's exact test p-value < 0.01 in genes up-regulated in electric organ. The number of genes is plotted for each term. The GO terms are colored by their assignment to molecular function, cellular component, or biological process.

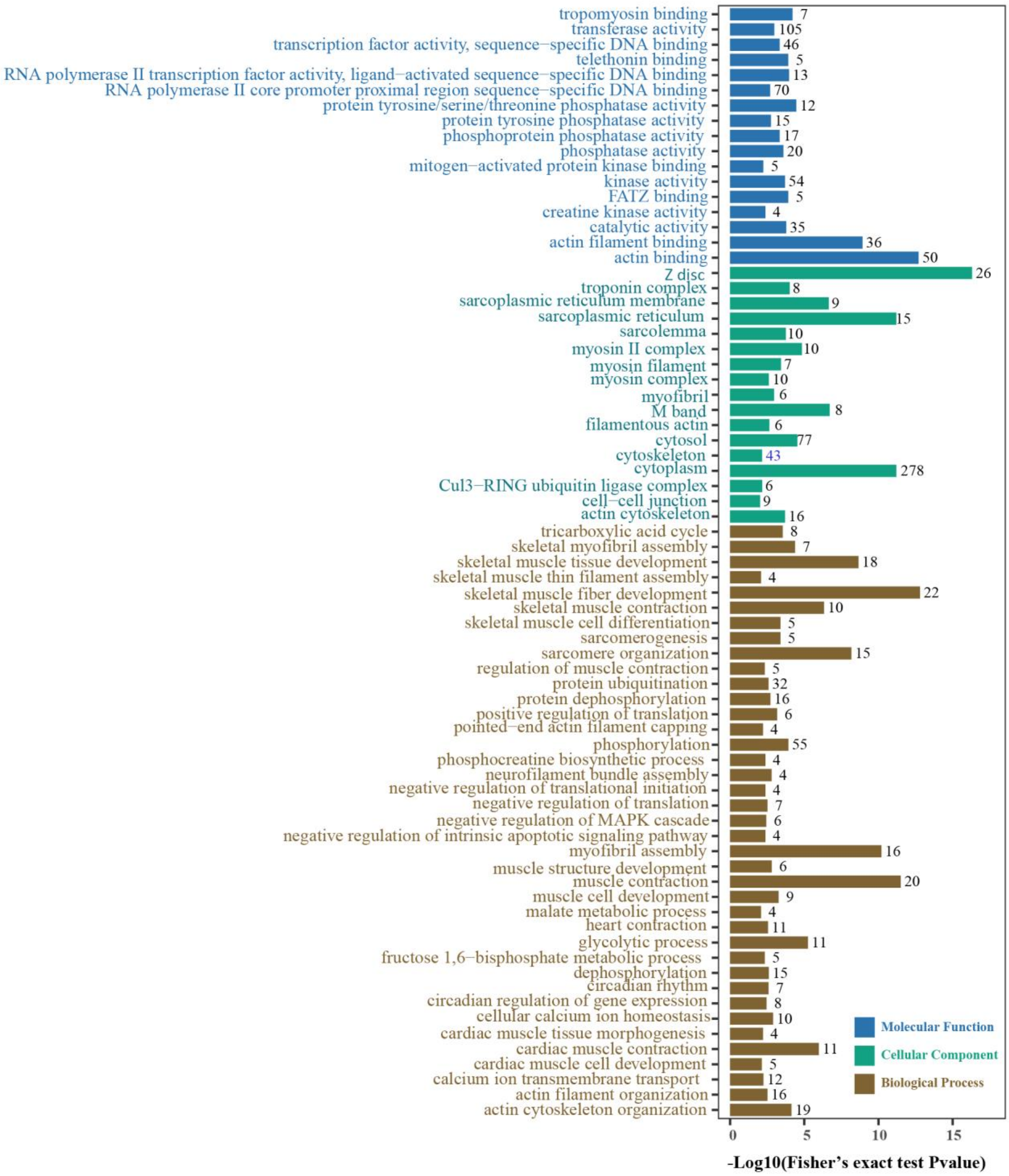

**Supplementary Fig. 2**

Significantly enriched Gene Ontology (GO) terms with Fisher's exact test Pvalue < 0.01 of genes down-regulated in electric organ (up regulated in skeleton muscle). The number of genes is plotted for each term. The GO terms are colored by their assignment to molecular function, cellular component, or biological process.

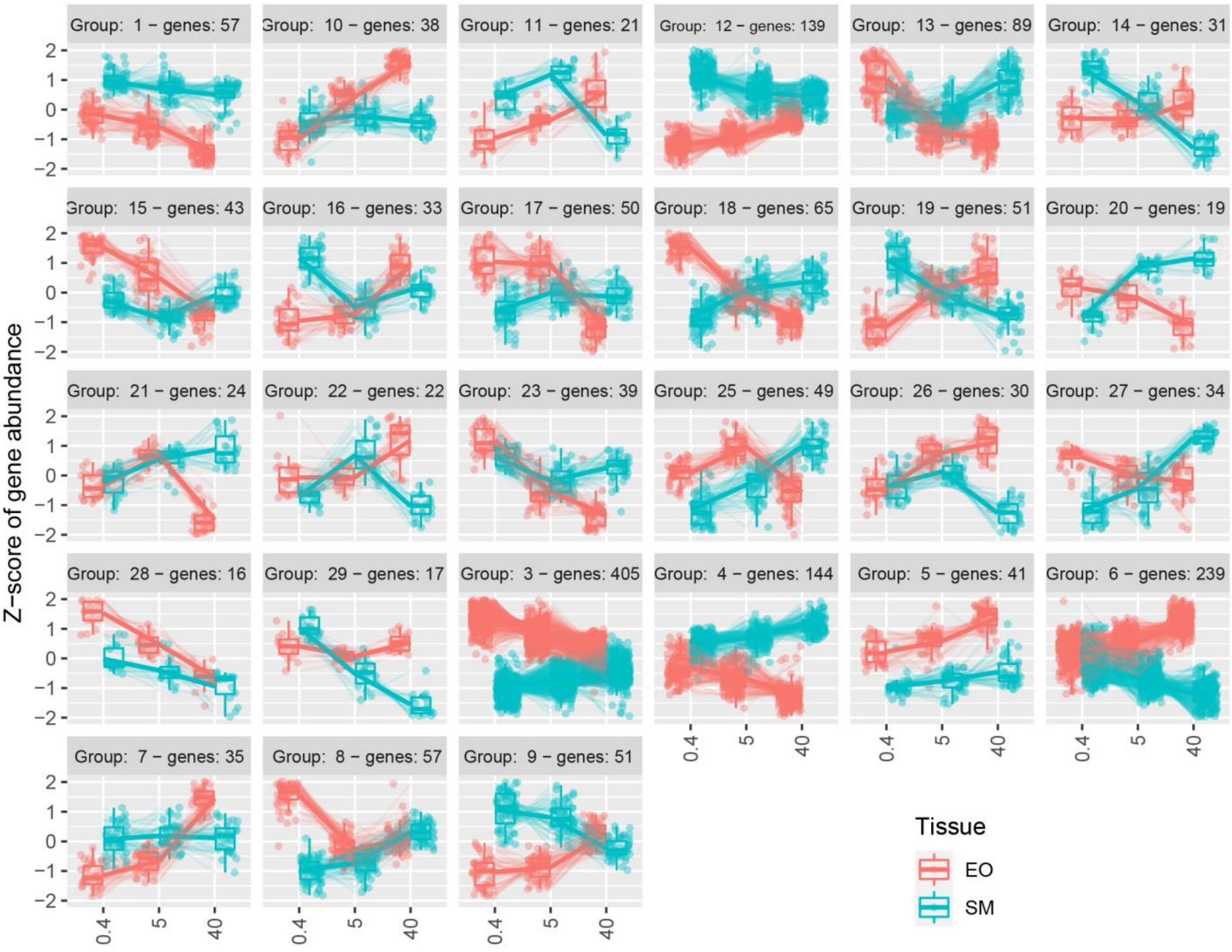

**Supplementary Fig. 3**

RNA-seq data clustering analysis based on EOD duration in electric organ (EO) and skeleton muscle (SM) of three F0 species. The x-axis for each group represents the EOD duration of the respective species: *C. compressirostris* (0.4ms), *C. tshokwe* (5ms) and *C. rhynchophorus* (40ms).

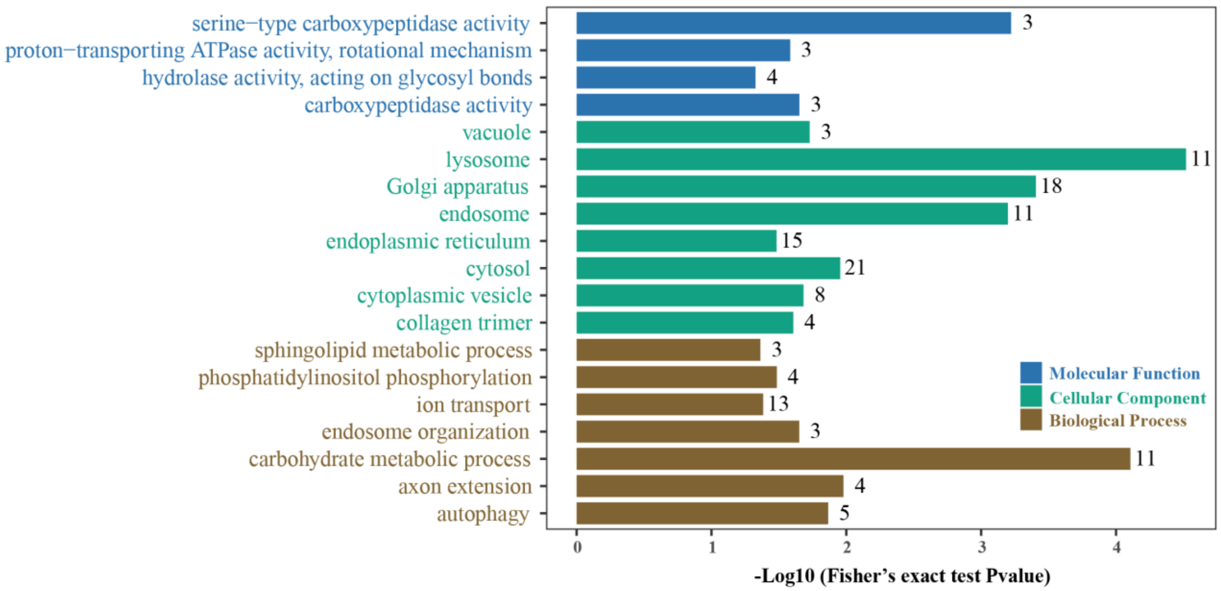

**Supplementary Fig. 4**

19 significantly enriched Gene Ontology (GO) terms with Fisher's exact test Pvalue < 0.05 for genes with increasing expression pattern over EOD duration change (Group 5 and 6). The number of genes is plotted for each term. The GO terms are colored by their assignment to molecular function, cellular component, or biological process.

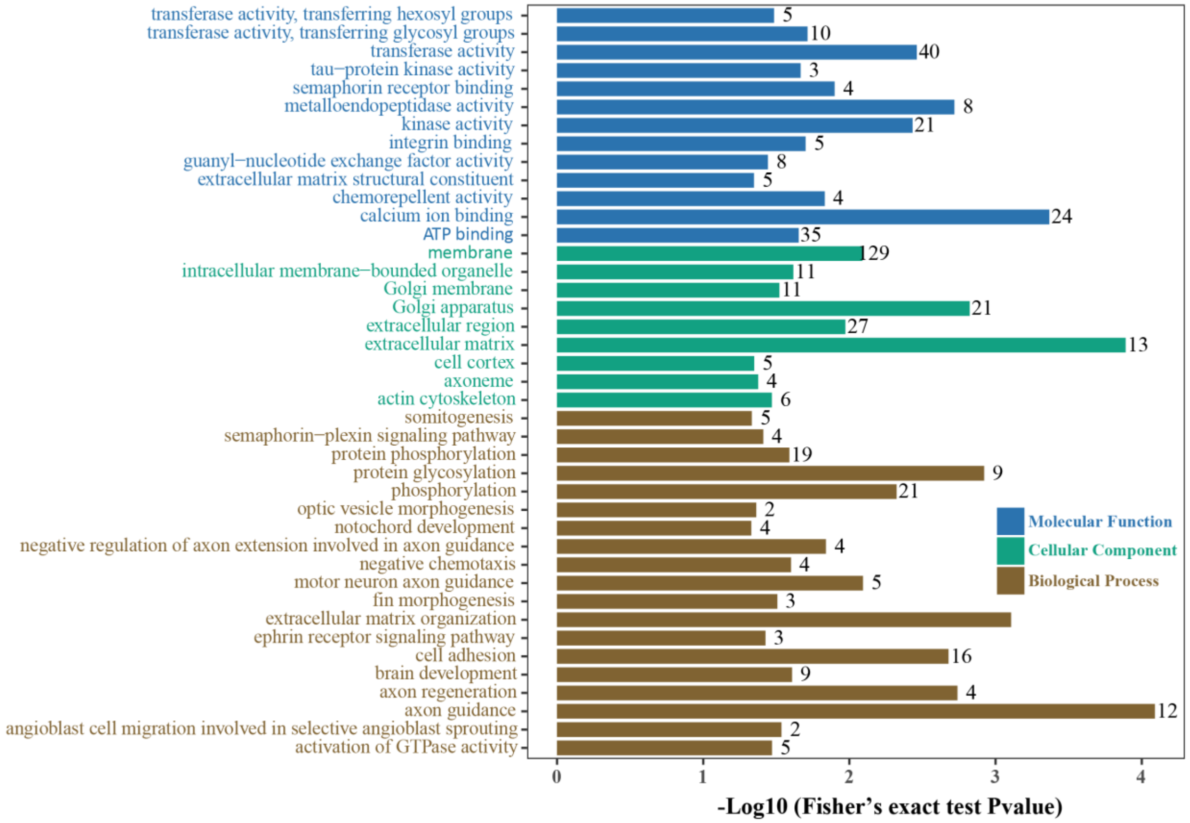

**Supplementary Fig. 5**

Significantly enriched Gene Ontology (GO) terms with Fisher's exact test Pvalue < 0.05 in genes with decreasing expression pattern over EOD duration change (Group 3). The number of genes is plotted for each term. The GO terms are colored by their assignment to molecular function, cellular component, or biological process.

**Supplementary Table 1** Genes up-regulated in all species/hybrids in the electric organ relative to sceletal muscle.

| ID | Blast Gene | Hightlighs of Predicted Function | Gene Description | Category | Average log2FC | Average Pvalue |
|---|---|---|---|---|---|---|
| maker-ptg000361l-augustus-gene-0.2-mRNA-1 | *ACTR3b* | F-actin dynamics / polymerization | ARP3 actin related protein 3 homolog B | cytoskeletal & sarcomeric | 3,02 | 3,1635E-20 |
| snap_masked-ptg000904l-processed-gene-5.5-mRNA-1 | *ARPC3* | F-actin dynamics / polymerization | actin related protein 2/3 complex subunit 3 | cytoskeletal & sarcomeric | 2,01 | 9,68947E-05 |
| maker-ptg001180l-augustus-gene-13.21-mRNA-1 | *ARPC5* | F-actin dynamics / polymerization | actin-related protein 2/3 complex subunit 5 | cytoskeletal & sarcomeric | 1,99 | 0,003541395 |
| maker-ptg002072l-snap-gene-6.44-mRNA-1 | *CARMIL1* | F-actin dynamics / polymerization | capping protein regulator and myosin 1 linker 1 | cytoskeletal & sarcomeric | 1,91 | 0,001040325 |
| maker-ptg000650l-snap-gene-7.49-mRNA-1 | *EPPK1* | controls reorganization of intermediate filaments | epiplakin 1 | cytoskeletal & sarcomeric | 7,74 | 1,21807E-11 |
| maker-ptg000346l-snap-gene-25.173-mRNA-1 | *GSN* | F-actin dynamics / polymerization | gelsolin | cytoskeletal & sarcomeric | 6,70 | 2,7189E-50 |
| maker-ptg001291l-augustus-gene-1.16-mRNA-1 | *MYO1d* | unconventional myosin; actin-based motor protein | unconventional myosin-Id | cytoskeletal & sarcomeric | 2,16 | 0,013895309 |
| maker-ptg001182l-snap-gene-5.74-mRNA-1 | *MTSS1* | regulation of F-actin dynamics | metastasis suppressor protein 1 | cytoskeletal & sarcomeric | 4,71 | 1,09085E-10 |
| maker-ptg000190l-snap-gene-4.78-mRNA-1 | *MYH10* | unconventional myosin; actin-based motor protein | myosin-10 | cytoskeletal & sarcomeric | 1,57 | 2,93467E-10 |
| maker-ptg001188l-snap-gene-5.24-mRNA-1 | *MYL4* | regulatory light chain of myosin | myosin light chain 4 | cytoskeletal & sarcomeric | 7,12 | 5,64592E-05 |
| snap_masked-ptg000028l-processed-gene-137.57-mRNA-1 | *MYO15a* | unconventional myosin; actin-based motor protein | unconventional myosin-XV | cytoskeletal & sarcomeric | 3,68 | 3,40016E-09 |
| maker-ptg001291l-snap-gene-1.34-mRNA-1 | *MYO1d* | unconventional myosin; actin-based motor protein | myosin ID | cytoskeletal & sarcomeric | 1,93 | 0,003762165 |
| maker-ptg001003l-snap-gene-2.16-mRNA-1 | *MYO1e* | unconventional myosin; actin-based motor protein | unconventional myosin-Ie | cytoskeletal & sarcomeric | 2,63 | 5,70736E-05 |
| maker-ptg002090l-augustus-gene-35.16-mRNA-1 | *MYO3b* | unconventional myosin; actin-based motor protein | myosin-IIIb | cytoskeletal & sarcomeric | 3,75 | 1,42712E-07 |
| maker-ptg000509l-snap-gene-4.4-mRNA-1 | *NHS* | regulator of actin remodelling | Nance-Horan syndrome protein | cytoskeletal & sarcomeric | 5,30 | 3,10465E-11 |
| maker-ptg002114l-augustus-gene-0.11-mRNA-1 | *PARVG* | regulation of cytoskeleton organization | parvin gamma | cytoskeletal & sarcomeric | 3,70 | 7,51203E-19 |
| snap_masked-ptg000650l-processed-gene-9.23-mRNA-1 | *PLEC* | cross-linking and stabilization of cytoskeletal intermediate filaments | plectin | cytoskeletal & sarcomeric | 2,62 | 8,77122E-22 |
| maker-ptg000014l-snap-gene-7.139-mRNA-1 | *MAP7d1* | microtubule cytoskeleton organization | MAP7 domain-containing protein 1 | cytoskeletal & sarcomeric | 8,32 | 6,90805E-34 |
| snap_masked-ptg000056l-processed-gene-18.17-mRNA-1 | *ATP10d* | catalytic component of a P4-ATPase flippase complex | ATPase phospholipid transporting 10D | membrane organization | 1,74 | 0,001111433 |
| maker-ptg000298l-snap-gene-2.25-mRNA-1 | *ESYT1* | ER-plasma membrane contact sites | extended synaptotagmin | membrane organization | 2,30 | 3,56639E-09 |
| maker-ptg000317l-snap-gene-0.11-mRNA-1 | *CALR3b* | Ca2+-binding chaperone in ER | calreticulin | other | 1,38 | 0,000700663 |
| maker-ptg000316l-snap-gene-4.241-mRNA-1 | *CAPN1a* | Ca2+-dependent, non-lysosomal cysteine protease | calpain-1 catalytic subunit | other | 3,26 | 1,5809E-41 |
| maker-ptg000214l-augustus-gene-8.178-mRNA-1 | *CAPN5* | Ca2+-dependent, non-lysosomal cysteine protease | calpain 5 | other | 4,07 | 3,13077E-28 |
| maker-ptg000886l-augustus-gene-0.134-mRNA-1 | *CARHSP1* | regulation of mRNA stability | calcium regulated heat stable protein 1 | other | 3,01 | 2,85152E-08 |
| maker-ptg000316l-snap-gene-7.47-mRNA-1 | *CRIP3* | metal ion binding activity | cysteine rich protein 3 | other | 4,90 | 1,31827E-14 |
| maker-ptg000644l-snap-gene-0.4-mRNA-1 | *GDPD4* | enable metal ion binding activity and phosphoric diester hydrolase acti | glycerophosphodiester phosphodiesterase domain containing 4 | other | 4,98 | 3,97129E-12 |
| maker-ptg000405l-snap-gene-38.13-mRNA-1 | *SMOC2* | secreted calcium-binding protein | SPARC related modular calcium binding 2 | other | 5,40 | 1,01506E-17 |
| maker-ptg000070l-snap-gene-5.18-mRNA-1 | *FLOT2a* | scaffolding protein within caveolar membranes | flotillin-2a | other | 5,77 | 1,81884E-53 |
| maker-ptg000040l-snap-gene-14.53-mRNA-1 | *NAT8l* | catalyzes the synthesis of N-acetylaspartate acid | N-acetyltransferase 8 | other | 7,97 | 1,96623E-07 |
| maker-ptg000102l-snap-gene-52.7-mRNA-1 | *ST8SIA5* | glycosphingolipid biosynthetic process | ST8 alpha-N-acetyl-neuraminide alpha-2,8-sialyltransferase 5 | other | 9,67 | 6,85106E-08 |
| maker-ptg000120l-snap-gene-2.54-mRNA-1 | *SYNGR3* | synaptic vesicule membrane protein | synaptogyrin-3 | other | 10,00 | 3,33741E-07 |
| maker-ptg000152l-snap-gene-4.203-mRNA-1 | *ZDHHC23* | palmitoyltransferase | zinc finger DHHC-type containing 23 | other | 6,43 | 5,59154E-07 |
| maker-ptg001827l-augustus-gene-0.22-mRNA-1 | *FAT3* | cell-cell adhesion | FAT atypical cadherin 3 | other | 2,72 | 0,001238961 |
| snap_masked-ptg000148l-processed-gene-1.18-mRNA-1 | *FAT4* | cell adhesion molecule | FAT atypical cadherin 4 | other | 3,97 | 3,45897E-11 |
| maker-ptg001421l-augustus-gene-5.0-mRNA-1 | *JCAD* | cell adhesion | junctional protein associated with coronary artery disease | other | 4,28 | 3,35791E-67 |
| maker-ptg001427l-augustus-gene-38.97-mRNA-1 | *KCTD9* | Substrate-specific adapter of a cullin-based E3 ubiquitin-protein ligase c | potassium channel tetramerization domain containing 9 | other | 1,77 | 3,05442E-05 |
| maker-ptg000361l-snap-gene-40.32-mRNA-1 | *LAMA1* | extracellular matrix protein | laminin subunit alpha-1 | other | 5,73 | 3,75043E-57 |
| maker-ptg000187l-augustus-gene-16.29-mRNA-1 | *MMP28* | extracellular matrix protein | matrix metallopeptidase 28 | other | 4,82 | 6,22492E-59 |
| maker-ptg000964l-snap-gene-1.24-mRNA-1 | *NCAM1* | cell adhesion molecule | neuronal cell adhesion molecule | other | 4,84 | 2,16487E-24 |
| maker-ptg000718l-snap-gene-13.14-mRNA-1 | *NDFIP2* | activates HECT domain-containing E3 ubiquitin-protein ligases | NEDD4 family-interacting protein 2 | other | 8,81 | 1,42211E-12 |
| maker-ptg000879l-snap-gene-1.65-mRNA-1 | *TMIGD1* | cell adhesion molecule | transmembrane and immunoglobulin domain containing 1 | other | 9,81 | 1,75298E-07 |
| maker-ptg001236l-augustus-gene-24.16-mRNA-1 | *TRH* | tripeptide hypothalamic regulatory hormone | thyrotropin releasing hormone | other | 9,79 | 5,36476E-17 |
| maker-ptg000774l-augustus-gene-4.234-mRNA-1 | *CTNNAL1* | modulation the Rho pathway signaling | catenin alpha like 1 | signaling | 5,91 | 2,6796E-49 |
| maker-ptg000049l-snap-gene-23.20-mRNA-1 | *FGF12* | regulation of voltage-gated sodium channels | fibroblast growth factor 12 | signaling | 8,72 | 5,84977E-17 |
| maker-ptg000028l-augustus-gene-28.23-mRNA-1 | *HEG1* | calcium ion binding activity | protein HEG homolog 1 | signaling | 8,38 | 8,41625E-14 |
| maker-ptg000159l-augustus-gene-6.0-mRNA-1 | *PCP4* | modulator of calcium-binding by calmodulin | calmodulin regulator protein PCP4 | signaling | 6,64 | 4,01068E-05 |
| maker-ptg000869l-snap-gene-20.89-mRNA-1 | *PVALB9* | cytosolic Ca2+-binding protein of the EF-hand superfamily | parvalbumin, thymic | signaling | 8,51 | 1,38958E-09 |
| maker-ptg000215l-snap-gene-13.31-mRNA-1 | *S100b* | cytosolic Ca2+-binding protein of the EF-hand superfamily | S100 calcium binding protein B | signaling | 8,14 | 1,01224E-22 |
| snap_masked-ptg001241l-processed-gene-1.62-mRNA-1 | *KL* | may be involved in the regulation of calcium and phosphorus homeosta | klotho | signaling | 5,39 | 0,000196154 |
| maker-ptg000830l-snap-gene-5.84-mRNA-1 | *SEMA5a* | ligand for receptor PLXNB3 | semaphorin-5A | signaling | 9,17 | 2,9895E-15 |
| maker-ptg000408l-augustus-gene-0.0-mRNA-1 | *ANXA4* | annexin family of calcium-dependent phospholipid binding proteins | annexin A4 | signaling | 5,55 | 6,72209E-60 |
| maker-ptg000084l-augustus-gene-28.11-mRNA-1 | *CAMK1d* | Ca2+/calmodulin-dependent protein kinase | calcium/calmodulin dependent protein kinase ID | signaling | 3,18 | 0,000145331 |
| maker-ptg000237l-snap-gene-9.30-mRNA-1 | *CAMK1g* | Ca2+/calmodulin-dependent protein kinase | calcium/calmodulin-dependent protein kinase type 1D | signaling | 4,56 | 0,000111371 |
| maker-ptg000008l-augustus-gene-0.0-mRNA-1 | *CHRNB4* | neuronal acetylcholine receptor subunit alpha; nonselective cation char | neuronal acetylcholine receptor subunit beta-4 | signaling | 6,28 | 2,08134E-05 |
| maker-ptg001610l-snap-gene-4.137-mRNA-1 | *GRIK3* | ionotropic glutamate receptor | glutamate ionotropic receptor kainate type subunit 3 | signaling | 6,15 | 5,72433E-05 |
| maker-ptg001029l-snap-gene-17.22-mRNA-1 | *GRIN2a* | ionotropic glutamate receptor | glutamate receptor ionotropic, NMDA 2A | signaling | 6,25 | 0,001464578 |
| maker-ptg000897l-snap-gene-2.61-mRNA-1 | *GRINA* | negative regulation of apoptotic signaling pathway | glutamate ionotropic receptor NMDA type subunit associated protein 1 | signaling | 3,24 | 7,33209E-61 |
| maker-ptg000068l-augustus-gene-18.3-mRNA-1 | *ITPR1* | InsP3-dependent ER Ca2+ channel | inositol 1,4,5-trisphosphate receptor type 1 | signaling | 4,07 | 9,87654E-18 |
| maker-ptg000838l-snap-gene-3.218-mRNA-1 | *NDRG3* | predicted to be involved in signal transduction | N-myc downstream-regulated gene 3 protein | signaling | 11,02 | 7,44456E-08 |
| snap_masked-ptg001427l-processed-gene-2.9-mRNA-1 | *P2RY2* | receptor for ATP and UTP coupled to G-proteins that activate a phospha | P2Y purinoceptor 2 | signaling | 3,78 | 0,001473877 |
| maker-ptg000267l-snap-gene-2.92-mRNA-1 | *PIEZO2* | mechanosensitive ion channel | piezo type mechanosensitive ion channel component 2 | signaling | 1,89 | 1,03736E-06 |
| maker-ptg001424l-augustus-gene-5.145-mRNA-1 | *RET* | receptor tyrosine-protein kinase | ret proto-oncogene | signaling | 10,14 | 2,32382E-12 |
| maker-ptg001563l-augustus-gene-5.35-mRNA-1 | *SGK1* | serine/threonine-protein kinase | serine/threonine-protein kinase Sgk1 | signaling | 7,79 | 6,63267E-38 |

| | | | | | | |
|---|---|---|---|---|---|---|
| maker-ptg000926l-snap-gene-6.19-mRNA-1 | *TRPV1* | transient receptor potential family of ion channels; nociception | transient receptor potential cation channel subfamily V member 1 | signaling | 2,43 | 0,005919225 |
| maker-ptg001088l-snap-gene-0.7-mRNA-1 | *SIX2a* | target ARE promoter elements in Na+/K+ adenosine triphosphatases | SIX homeobox 2 | transcription factor | 3,05 | 1,22103E-27 |
| maker-ptg000783l-snap-gene-6.78-mRNA-1 | *HEY1* | developing cardiac conduction pathway | hes related family bHLH transcription factor with YRPW motif 1 | transcription factor | 6,02 | 1,5325E-13 |
| maker-ptg000149l-augustus-gene-2.87-mRNA-1 | *ETV5* | transcription factor | ETS translocation variant 5 | transcription factor | 5,11 | 1,99403E-55 |
| snap_masked-ptg000737l-processed-gene-6.55-mRNA-1 | *FOXL2* | regulate distinct female sex determining pathways | forkhead box protein L2 | transcription factor | 8,52 | 4,14855E-07 |
| maker-ptg001740l-augustus-gene-1.112-mRNA-1 | *KLF5* | rebalance potassium channels | Krueppel-like factor 5 | transcription factor | 8,39 | 5,04913E-06 |
| maker-ptg000008l-snap-gene-10.43-mRNA-1 | *MEF2a* | transcriptional activator for numerous muscle-specific genes | myocyte-specific enhancer factor 2A | transcription factor | 3,85 | 3,78175E-50 |
| maker-ptg001270l-snap-gene-47.16-mRNA-1 | *MEF2b* | transcriptional activator for numerous muscle-specific genes | myocyte-specific enhancer factor 2B | transcription factor | 6,92 | 8,45833E-68 |
| maker-ptg000345l-snap-gene-4.29-mRNA-1 | *ANO10* | calcium-activated chloride channel | anoctamin 10 | transmembrane ion transport | 2,37 | 1,21792E-13 |
| maker-ptg000406l-snap-gene-3.4-mRNA-1 | *ANO5* | calcium-activated chloride channel | anoctamin 5 | transmembrane ion transport | 2,76 | 3,56495E-17 |
| maker-ptg000979l-snap-gene-12.179-mRNA-1 | *ANO6* | calcium-activated nonselective cation (SCAN) channel which acts as a re | anoctamin 6 | transmembrane ion transport | 1,82 | 0,000445473 |
| maker-ptg000970l-augustus-gene-2.127-mRNA-1 | *ATP1a1* | Na/K-ATPase α-subunit | sodium/potassium-transporting ATPase subunit alpha-1 | transmembrane ion transport | 10,55 | 2,18943E-05 |
| snap_masked-ptg001156l-processed-gene-0.19-mRNA-1 | *ATP1a2a* | Na/K-ATPase α-subunit | sodium/potassium-transporting ATPase subunit alpha-2 | transmembrane ion transport | 3,07 | 2,63171E-12 |
| maker-ptg001047l-snap-gene-1.63-mRNA-1 | *ATP1b1a* | Na/K-ATPase β-subunit | ATPase NA+/K+ transporting beta 1a | transmembrane ion transport | 6,59 | 3,63255E-59 |
| maker-ptg000509l-snap-gene-9.39-mRNA-1 | *ATP1b1b* | Na/K-ATPase β-subunit | ATPase NA+/K+ transporting beta 1b | transmembrane ion transport | 4,51 | 2,06472E-06 |
| maker-ptg000993l-snap-gene-3.19-mRNA-1 | *ATP2a2* | sarcoplasmic/endoplasmic reticulum calcium ATPase 2 | sarcoplasmic/endoplasmic reticulum calcium ATPase 2 | transmembrane ion transport | 1,90 | 5,79784E-08 |
| snap_masked-ptg000904l-processed-gene-1.73-mRNA-1 | *ATP2a2b* | sarcoplasmic/endoplasmic reticulum calcium ATPase 2 | sarcoplasmic/endoplasmic reticulum calcium ATPase 2b | transmembrane ion transport | 2,72 | 5,65072E-08 |
| maker-ptg000084l-augustus-gene-36.7-mRNA-1 | *ATP2b1a* | plasma membrane calcium-transporting ATPase 1 | plasma membrane calcium-transporting ATPase 1a | transmembrane ion transport | 2,96 | 4,08993E-15 |
| maker-ptg000084l-snap-gene-36.24-mRNA-1 | *ATP2b1b* | plasma membrane calcium-transporting ATPase 1 | plasma membrane calcium-transporting ATPase 1b | transmembrane ion transport | 3,66 | 1,07062E-27 |
| maker-ptg000135l-augustus-gene-78.38-mRNA-1 | *ATP2b3* | ATPase plasma membrane Ca2+ transporting 3 | ATPase plasma membrane Ca2+ transporting 3 | transmembrane ion transport | 5,65 | 1,02377E-05 |
| maker-ptg000347l-snap-gene-10.66-mRNA-1 | *ATP2c1* | secretory pathway Ca2+-ATPase | ATPase secretory pathway Ca2+ transporting 1 | transmembrane ion transport | 2,14 | 0,000146327 |
| maker-ptg000032l-augustus-gene-1.26-mRNA-1 | *ATP6ap2* | subunit of vacuolar-type H+-ATPase | ATPase H+ transporting accessory protein 2 | transmembrane ion transport | 1,78 | 2,28333E-09 |
| maker-ptg000501l-augustus-gene-3.39-mRNA-1 | *ATP6v1f* | subunit of vacuolar-type H+-ATPase | ATPase H+ transporting V1 subunit F | transmembrane ion transport | 1,48 | 1,99856E-09 |
| maker-ptg001427l-snap-gene-4.23-mRNA-1 | *CACNA1b* | calcium channel subunit | voltage-dependent N-type calcium channel subunit alpha-1B | transmembrane ion transport | 5,19 | 2,70467E-08 |
| maker-ptg001182l-snap-gene-8.79-mRNA-1 | *CNGB1* | nonselective cation channel | cyclic nucleotide-gated cation channel beta-1 | transmembrane ion transport | 5,76 | 2,75381E-05 |
| snap_masked-ptg000084l-processed-gene-43.23-mRNA-1 | *CRACR2aa* | Ca2+ binding protein that is a key regulator of CRAC channel-mediated | calcium release activated channel regulator 2Aa | transmembrane ion transport | 3,67 | 0,001092994 |
| maker-ptg001069l-augustus-gene-3.65-mRNA-1 | *GABRA1* | subunit of ligand-gated chloride channel | gamma-aminobutyric acid receptor subunit alpha-1 | transmembrane ion transport | 5,44 | 1,17436E-05 |
| snap_masked-ptg000509l-processed-gene-7.11-mRNA-1 | *GABRG3* | subunit of ligand-gated chloride channel | gamma-aminobutyric acid receptor subunit gamma-3 | transmembrane ion transport | 2,78 | 0,000289984 |
| snap_masked-ptg002179l-processed-gene-1.70-mRNA-1 | *GLRBB* | ligand-gated chloride channel | glycine receptor, beta b | transmembrane ion transport | 9,03 | 2,58552E-29 |
| maker-ptg000559l-snap-gene-2.76-mRNA-1 | *HCN2* | hyperpolarization-activated ion channel | hyperpolarization activated cyclic nucleotide gated potassium and sodium channel 2 | transmembrane ion transport | 3,65 | 1,79576E-09 |
| maker-ptg000028l-snap-gene-81.10-mRNA-1 | *KCNA7a_1* | voltage-gated potassium channel | potassium voltage-gated channel subfamily A member 7a | transmembrane ion transport | 4,60 | 2,58472E-11 |
| maker-ptg000028l-snap-gene-81.8-mRNA-1 | *KCNA7a_2* | voltage-gated potassium channel | potassium voltage-gated channel subfamily A member 7a | transmembrane ion transport | 8,27 | 3,55527E-12 |
| maker-ptg001427l-snap-gene-13.20-mRNA-1 | *KCNIP3* | voltage-gated potassium channel | calsenilin | transmembrane ion transport | 5,11 | 0,000993571 |
| maker-ptg000265l-est_gff_est2genome-gene-6.33-mRNA-1 | *KCNJ2* | inwardly rectifying potassium channel | inward rectifier potassium channel 2 | transmembrane ion transport | 5,53 | 1,12219E-20 |
| maker-ptg000830l-augustus-gene-5.123-mRNA-1 | *KCNJ9* | inwardly rectifying potassium channel | G protein-activated inward rectifier potassium channel 3 | transmembrane ion transport | 5,77 | 2,63245E-10 |
| snap_masked-ptg001118l-processed-gene-0.13-mRNA-1 | *KCNK2* | potassium two pore domain channel | potassium channel subfamily K member 2 | transmembrane ion transport | 6,15 | 9,23801E-14 |
| maker-ptg000697l-snap-gene-6.109-mRNA-1 | *KCNQ5* | voltage-gated potassium channel | potassium voltage-gated channel subfamily Q member 5 | transmembrane ion transport | 5,94 | 0,000938938 |
| maker-ptg001106l-augustus-gene-0.48-mRNA-1 | *MCOLN1* | intracellular cation channel | mucolipin-1 | transmembrane ion transport | 6,37 | 6,01855E-26 |
| maker-ptg001057l-snap-gene-3.24-mRNA-1 | *MCOLN3* | intracellular cation channel | mucolipin-3 | transmembrane ion transport | 5,68 | 4,30182E-06 |
| maker-ptg000253l-augustus-gene-20.10-mRNA-1 | *SCN1ba* | voltage-gated sodium channel beta-subunit (regulatory) | sodium channel subunit beta-1 | transmembrane ion transport | 3,84 | 1,29832E-07 |
| maker-ptg001188l-snap-gene-6.4-mRNA-1 | *SCN4aa* | voltage-gated sodium channel alpha-subunit | sodium channel protein type 4 subunit alpha A | transmembrane ion transport | 10,98 | 1,27373E-11 |
| maker-ptg002239l-snap-gene-5.5-mRNA-1 | *SCN4b* | voltage-gated sodium channel | sodium voltage-gated channel beta subunit 4 | transmembrane ion transport | 5,34 | 8,18062E-47 |
| maker-ptg000477l-augustus-gene-1.10-mRNA-1 | *VDAC1* | ion channel in the outer mitochondrial membrane and also the outer c | voltage-dependent anion-selective channel protein 1 | transmembrane ion transport | 2,69 | 2,46626E-06 |
| maker-ptg000051l-snap-gene-132.157-mRNA-1 | *TMEM206* | proton-activated chloride channel | transmembrane protein 206 | transmembrane ion transport | 1,98 | 6,00534E-10 |
| maker-ptg001300l-snap-gene-4.198-mRNA-1 | *TMEM63c* | Ca2+ permeable cation channel at ER/mitochondria contact sites | transmembrane protein 63C | transmembrane ion transport | 4,00 | 0,001858763 |
| maker-ptg000187l-augustus-gene-15.2-mRNA-1 | *TMEM120a* | mechanosensing ion channel | transmembrane protein 120A | transmembrane ion transport | 6,05 | 2,33605E-44 |
| maker-ptg000148l-snap-gene-9.4-mRNA-1 | *SLC24a2* | calcium, potassium:sodium antiporter | solute carrier family 24 member 2 | transmembrane ion transport | 9,06 | 1,1057E-37 |
| snap_masked-ptg000102l-processed-gene-12.46-mRNA-1 | *SLC4a4* | sodium bicarbonate cotransporter | solute carrier family 4 member 4 | transmembrane ion transport | 5,67 | 1,3753E-11 |
| snap_masked-ptg000004l-processed-gene-3.9-mRNA-1 | *SLC8a1a* | sodium/calcium exchanger | solute carrier family 8 member 1a | transmembrane ion transport | 5,01 | 3,00748E-13 |
| maker-ptg002207l-snap-gene-0.59-mRNA-1 | *ABCA2* | probable lipid transporter that modulates cholesterol sequestration in t | ATP binding cassette subfamily A member 2 | transmembrane ion transport | 2,21 | 1,89872E-10 |
| maker-ptg001844l-snap-gene-4.2-mRNA-1 | *ABCB1* | ABC transporter; translocates drugs and phospholipids across the mem | ATP binding cassette subfamily B member 1 | transmembrane ion transport | 4,72 | 0,000481822 |
| maker-ptg001436l-snap-gene-0.52-mRNA-1 | *ABCG2* | ABC transporter of broad substrate specificity | ATP-binding cassette sub-family G member 2 | transmembrane ion transport | 2,24 | 0,008846064 |
| maker-ptg000326l-snap-gene-3.64-mRNA-1 | *SLC13a1* | sodium:sulfate symporter | solute carrier family 13 member 1 | transmembrane ion transport | 3,41 | 3,99214E-05 |
| snap_masked-ptg000878l-processed-gene-2.22-mRNA-1 | *SLC17a5* | membrane transporter that exports free sialic acids | solute carrier family 17 member 5 | transmembrane ion transport | 3,88 | 1,23265E-08 |
| maker-ptg001427l-snap-gene-14.26-mRNA-1 | *SLC23a2* | sodium/ascorbate cotransporter | solute carrier family 23 member 2 | transmembrane ion transport | 2,45 | 3,35477E-05 |
| snap_masked-ptg001837l-processed-gene-1.59-mRNA-1 | *SLC25a23* | calcium-dependent mitochondrial solute carrier on the inner mitochon | solute carrier family 25 member 23 | transmembrane ion transport | 5,13 | 0,000247848 |
| maker-ptg000102l-augustus-gene-64.1-mRNA-1 | *SLC25a25b* | calcium-binding mitochondrial carrier on the inner mitochondrial mem | solute carrier family 25 member 25b | transmembrane ion transport | 2,65 | 0,003639241 |
| maker-ptg000184l-augustus-gene-2.1-mRNA-1 | *SLC5a7* | sodium ion- and chloride ion-dependent high-affinity transporter that r | solute carrier family 5 member 7 | transmembrane ion transport | 5,91 | 0,000293221 |
| maker-ptg000175l-augustus-gene-7.91-mRNA-1 | *SLC5a9* | sodium/glucose cotransporter 4 | solute carrier family 5 member 9 | transmembrane ion transport | 2,80 | 6,19949E-11 |
| maker-ptg001587l-snap-gene-9.16-mRNA-1 | *SLC6a17* | sodium-dependent vesicular transporter selective for proline, glycine, le | solute carrier family 6 member 17 | transmembrane ion transport | 8,15 | 5,51716E-10 |
| maker-ptg000800l-augustus-gene-2.11-mRNA-1 | *SLC6a2* | sodium- and chloride-dependent transport of norepinephrine | solute carrier family 6 member 2 | transmembrane ion transport | 5,50 | 3,30148E-15 |
| maker-ptg001004l-snap-gene-4.176-mRNA-1 | *SLC6a6* | sodium- and chloride-dependent transport of taurine and beta-alanine | solute carrier family 6 member 6 | transmembrane ion transport | 4,10 | 7,32652E-09 |

**Supplementary Table 2** Genes down-regulated in all species/hybrids in the electric organ relative to sceletal muscle.

| ID | Blast Gene | Highlights of Predicted Function | Gene Description | Category | Average log2FC | Average Pvalue |
|---|---|---|---|---|---|---|
| snap_masked-ptg000308l-processed-gene-8.27-mRNA-1 | *ABI2a* | regulator of actin cytoskeleton dynamics | abl interactor 2a | cytoskeletal & sarcomeric | -1,94 | 0,004180823 |
| maker-ptg000512l-snap-gene-2.123-mRNA-1 | *ABLIM1b* | binds to actin filaments and mediates interactions between actin and cytoplasmic targets | actin-binding LIM protein 1 | cytoskeletal & sarcomeric | -2,88 | 1,69993E-09 |
| maker-ptg001458l-snap-gene-2.2-mRNA-1 | *ABRaa* | muscle specific actin-binding protein; involved in skeletal muscle hypertrophy and atrophy | actin-binding Rho-activating protein | cytoskeletal & sarcomeric | -5,52 | 1,11159E-06 |
| maker-ptg001416l-augustus-gene-5.37-mRNA-1 | *ABRab* | muscle specific actin-binding protein; involved in skeletal muscle hypertrophy and atrophy | actin binding Rho activating protein | cytoskeletal & sarcomeric | -5,37 | 5,05532E-07 |
| maker-ptg000306l-augustus-gene-0.39-mRNA-1 | *ACTA1* | actin isoform, striated muscle | actin, alpha 1, skeletal muscle | cytoskeletal & sarcomeric | -5,17 | 9,36728E-19 |
| maker-ptg001070l-augustus-gene-5.71-mRNA-1 | *ACTB1* | actin isoform, cytoplasmic | actin, cytoplasmic 1 | cytoskeletal & sarcomeric | -1,95 | 0,00022477 |
| maker-ptg000171l-snap-gene-9.31-mRNA-1 | *ACTC1a* | actin isoform, striated muscle | actin, alpha cardiac muscle 1a | cytoskeletal & sarcomeric | -8,11 | 8,1603E-05 |
| snap_masked-ptg002519l-processed-gene-0.5-mRNA-1 | *ACTC1b* | actin isoform, striated muscle | actin, alpha cardiac muscle 1b | cytoskeletal & sarcomeric | -4,70 | 0,004466918 |
| maker-ptg000067l-snap-gene-17.2-mRNA-1 | *ACTC2* | actin isoform, striated muscle | actin, alpha, cardiac muscle 2 | cytoskeletal & sarcomeric | -7,94 | 0,006969056 |
| maker-ptg000501l-snap-gene-3.13-mRNA-1 | *CAPZA2* | F-actin binding protein | F-actin-capping protein subunit alpha-2 | cytoskeletal & sarcomeric | -2,35 | 1,10876E-11 |
| maker-ptg000693l-snap-gene-21.128-mRNA-1 | *CAPZB* | F-actin binding protein | F-actin-capping protein subunit beta | cytoskeletal & sarcomeric | -1,32 | 9,88642E-06 |
| maker-ptg000196l-augustus-gene-4.0-mRNA-1 | *KY* | cytoskeleton-associated protease required for normal muscle growth | kyphoscoliosis peptidase | cytoskeletal & sarcomeric | -7,80 | 0,000264434 |
| maker-ptg000007l-snap-gene-2.65-mRNA-1 | *MYBPC1* | thick filament-associated protein located in the crossbridge region of vertebrate striated muscle A bands | myosin binding protein C, slow type | cytoskeletal & sarcomeric | -6,60 | 0,000728724 |
| snap_masked-ptg000028l-processed-gene-83.4-mRNA-1 | *MYBPC2* | thick filament-associated protein located in the crossbridge region of vertebrate striated muscle A bands | myosin-binding protein C, fast-type | cytoskeletal & sarcomeric | -6,01 | 0,001321102 |
| maker-ptg000333l-augustus-gene-6.128-mRNA-1 | *MYBPC3* | thick filament-associated protein located in the crossbridge region of vertebrate striated muscle A bands | myosin-binding protein C, cardiac-type | cytoskeletal & sarcomeric | -8,23 | 4,50551E-05 |
| maker-ptg000962l-augustus-gene-12.1-mRNA-1 | *MYBPH* | binds to myosin; probably involved in interaction with thick myofilaments in the A-band | myosin-binding protein H | cytoskeletal & sarcomeric | -4,84 | 0,003814769 |
| maker-ptg000085l-augustus-gene-17.7-mRNA-1 | *MYH2* | unconventional myosin; actin-based motor protein | myosin heavy chain, fast skeletal muscle | cytoskeletal & sarcomeric | -8,17 | 5,93451E-19 |
| snap_masked-ptg000407l-processed-gene-2.145-mRNA-1 | *MYH1* | myosin heavy chain beta isoform expressed primarily in the heart, but also in skeletal muscles | myosin 1 | cytoskeletal & sarcomeric | -8,60 | 4,34569E-06 |
| maker-ptg000133l-augustus-gene-18.18-mRNA-1 | *MYL1* | non-regulatory myosin light chain | myosin light chain 1 | cytoskeletal & sarcomeric | -5,57 | 1,66255E-08 |
| maker-ptg001175l-snap-gene-0.158-mRNA-1 | *MYL2* | myosin regulatory light chain 2 | myosin regulatory light chain 2, cardiac muscle | cytoskeletal & sarcomeric | -6,12 | 0,000536607 |
| maker-ptg000187l-snap-gene-16.83-mRNA-1 | *MYL2b* | myosin regulatory light chain 2 | myosin regulatory light chain 2B, cardiac muscle | cytoskeletal & sarcomeric | -8,19 | 5,84804E-05 |
| maker-ptg000100l-snap-gene-11.24-mRNA-1 | *MYL4* | myosin regulatory light chain | myosin light chain 4 | cytoskeletal & sarcomeric | -7,66 | 0,001195604 |
| maker-ptg000135l-snap-gene-52.42-mRNA-1 | *MYLK2* | myosin light chain kinase | myosin light chain kinase 2 | cytoskeletal & sarcomeric | -4,77 | 0,000103916 |
| maker-ptg000441l-augustus-gene-6.2-mRNA-1 | *MYO16* | unconventional myosin; actin-based motor protein | unconventional myosin-XVI | cytoskeletal & sarcomeric | -4,85 | 0,006179819 |
| maker-ptg000256l-snap-gene-3.25-mRNA-1 | *MYO18a* | unconventional myosin; actin-based motor protein | unconventional myosin-XVIIIa | cytoskeletal & sarcomeric | -4,19 | 1,94298E-13 |
| maker-ptg000960l-augustus-gene-0.14-mRNA-1 | *MYO18b* | unconventional myosin; actin-based motor protein | myosin XVIIIB | cytoskeletal & sarcomeric | -3,94 | 8,08729E-15 |
| maker-ptg000193l-snap-gene-2.47-mRNA-1 | *MYOM1* | major component of the vertebrate myofibrillar M band | myomesin 1 | cytoskeletal & sarcomeric | -7,91 | 1,13806E-07 |
| maker-ptg000110l-augustus-gene-2.12-mRNA-1 | *MYOM2* | major component of the vertebrate myofibrillar M band | myomesin 2 | cytoskeletal & sarcomeric | -4,02 | 3,46432E-14 |
| maker-ptg000405l-snap-gene-31.60-mRNA-1 | *MYOZ1* | involved in linking Z-disk proteins | myozenin 1 | cytoskeletal & sarcomeric | -6,71 | 6,15331E-07 |
| maker-ptg000314l-augustus-gene-4.2-mRNA-1 | *MYOZ2* | involved in linking Z-disk proteins | myozenin 2 | cytoskeletal & sarcomeric | -7,19 | 0,001878985 |
| maker-ptg000182l-snap-gene-3.82-mRNA-1 | *MYOZ3* | involved in linking Z-disk proteins | myozenin 3 | cytoskeletal & sarcomeric | -6,59 | 0,001769106 |
| maker-ptg001112l-snap-gene-5.13-mRNA-1 | *NEB* | binds and stabilize F-actin in the sarcomere | nebulin | cytoskeletal & sarcomeric | -7,40 | 0,000736895 |
| maker-ptg000512l-snap-gene-3.14-mRNA-1 | *NRAP* | may be involved in anchoring the terminal actin filaments in the myofibril to the membrane | nebulin related anchoring protein | cytoskeletal & sarcomeric | -3,92 | 1,29313E-19 |
| maker-ptg002114l-snap-gene-0.37-mRNA-1 | *PARVB* | adapter protein involved in the reorganization of the actin cytoskeleton | parvin beta | cytoskeletal & sarcomeric | -5,41 | 3,02009E-05 |
| maker-ptg000572l-snap-gene-14.41-mRNA-1 | *SMYHC1* | myosin heavy chain; actin-based motor protein | slow myosin heavy chain 1 | cytoskeletal & sarcomeric | -7,57 | 0,004522565 |
| maker-ptg000572l-augustus-gene-14.18-mRNA-1 | *SMYHC2* | myosin heavy chain; actin-based motor protein | slow myosin heavy chain 2 | cytoskeletal & sarcomeric | -5,20 | 0,001635808 |
| maker-ptg000135l-augustus-gene-79.30-mRNA-1 | *TNNC1* | component of troponin complex; regulation of muscle contraction | troponin C, slow skeletal and cardiac muscles | cytoskeletal & sarcomeric | -7,24 | 0,000307439 |
| maker-ptg000135l-snap-gene-52.7-mRNA-1 | *TNNC2* | component of troponin complex; regulation of muscle contraction | troponin C, skeletal muscle | cytoskeletal & sarcomeric | -4,81 | 5,01121E-11 |
| maker-ptg000223l-snap-gene-2.48-mRNA-1 | *TNNI2* | component of troponin complex; regulation of muscle contraction | troponin I, fast skeletal muscle | cytoskeletal & sarcomeric | -7,27 | 3,13868E-23 |
| maker-ptg000445l-snap-gene-11.25-mRNA-1 | *TPM1* | actin-binding protein; in association with the troponin complex involved in the striated muscle contraction | tropomyosin alpha-1 chain | cytoskeletal & sarcomeric | -4,81 | 6,55146E-15 |
| maker-ptg001553l-augustus-gene-0.54-mRNA-1 | *TPM2* | actin-binding protein; in association with the troponin complex involved in the striated muscle contraction | tropomyosin beta chain | cytoskeletal & sarcomeric | -6,67 | 1,53935E-06 |
| maker-ptg001563l-snap-gene-12.53-mRNA-1 | *TPM3* | actin-binding protein; in association with the troponin complex involved in the striated muscle contraction | tropomyosin alpha-3 chain | cytoskeletal & sarcomeric | -3,70 | 4,35071E-06 |
| maker-ptg000555l-augustus-gene-11.55-mRNA-1 | *TTN* | sarcomere organization | titin | cytoskeletal & sarcomeric | -2,95 | 3,70566E-07 |
| snap_masked-ptg000176l-processed-gene-4.104-mRNA-1 | *HSPB11* | disassembly of the sarcomeres | heat shock protein beta-11 | cytoskeletal & sarcomeric | -4,89 | 5,99473E-07 |
| maker-ptg001400l-snap-gene-3.115-mRNA-1 | *CAPN3* | muscle-specific calpain; calcium-activated non-lysosomal thiol-protease | calpain 3 | other | -4,51 | 3,46551E-17 |
| maker-ptg000222l-snap-gene-9.11-mRNA-1 | *CDH20* | calcium-dependent cell adhesion protein | cadherin-20 | other | -3,77 | 0,000127412 |
| maker-ptg000061l-augustus-gene-13.21-mRNA-1 | *CDH26* | calcium-dependent cell adhesion protein | cadherin-like protein 26 | other | -2,83 | 0,005372135 |
| maker-ptg000534l-snap-gene-5.169-mRNA-1 | *KCMF1* | E3 ubiquitin-protein ligase | potassium channel modulatory factor 1 | other | -1,41 | 7,97951E-07 |
| snap_masked-ptg002090l-processed-gene-16.20-mRNA-1 | *ACOT12* | fatty acid metabolic process | acyl-CoA thioesterase 12 | other | -7,79 | 0,00346148 |
| maker-ptg001966l-augustus-gene-10.17-mRNA-1 | *CKM* | reversibly catalyzes the transfer of phosphate between ATP and various phosphogens | creatine kinase M-type | other | -8,43 | 5,01097E-09 |
| maker-ptg000393l-augustus-gene-30.5-mRNA-1 | *CKMT1a* | reversibly catalyzes the transfer of phosphate between ATP and various phosphogens | creatine kinase U-type, mitochondrial | other | -7,81 | 4,06172E-07 |
| maker-ptg000323l-snap-gene-1.83-mRNA-1 | *EPN2* | interacts with clathrin | epsin-2 | other | -7,50 | 1,26567E-05 |
| maker-ptg000299l-snap-gene-6.20-mRNA-1 | *GLS2B* | glutaminase activity | glutaminase kidney isoform, mitochondrial | other | -7,18 | 1,23041E-05 |
| snap_masked-ptg001143l-processed-gene-2.56-mRNA-1 | *IGFN1* | cell adhesion | immunoglobulin-like and fibronectin type III domain-containing protein 1 | other | -7,80 | 5,98234E-36 |
| maker-ptg000487l-snap-gene-5.15-mRNA-1 | *LDHA* | catalyzes the conversion of L-lactate and NAD to pyruvate and NADH in the final step of anaerobic glycolysis | L-lactate dehydrogenase A chain | other | -7,47 | 0,000293053 |
| maker-ptg000254l-augustus-gene-6.0-mRNA-1 | *TECR* | involved in both the production of very long-chain fatty acids | very-long-chain enoyl-CoA reductase | other | -7,82 | 0,000632574 |
| maker-ptg000665l-snap-gene-7.139-mRNA-1 | *MYOC* | secreted glycoprotein regulating the activation of different signaling pathways in adjacent cells | myocilin | signaling | -2,10 | 0,00200062 |
| maker-ptg000558l-snap-gene-4.101-mRNA-1 | *CALM3* | calcium-binding EF-hand protein | calmodulin 3 | signaling | -2,84 | 8,29055E-05 |
| snap_masked-ptg000869l-processed-gene-20.110-mRNA-1 | *PVALB2* | Ca2+-binding protein of the EF-hand superfamily | parvalbumin-2 | signaling | -8,61 | 0,000478414 |
| maker-ptg000974l-augustus-gene-10.59-mRNA-1 | *PVALB4* | Ca2+-binding protein of the EF-hand superfamily | parvalbumin 4 | signaling | -6,32 | 0,000158436 |
| maker-ptg002802l-snap-gene-0.11-mRNA-1 | *PVALB7* | Ca2+-binding protein of the EF-hand superfamily | parvalbumin 7 | signaling | -8,33 | 5,15658E-05 |
| maker-ptg000481l-snap-gene-2.8-mRNA-1 | *ADRB2* | beta-2-adrenergic receptor | adrenoceptor beta 2 | signaling | -4,35 | 1,00955E-12 |
| maker-ptg000135l-augustus-gene-82.84-mRNA-1 | *GPR173* | G-protein coupled receptor | probable G-protein coupled receptor 173 | signaling | -8,04 | 0,00013504 |
| maker-ptg001004l-augustus-gene-9.10-mRNA-1 | *SBK1* | serine/threonine kinase activity | serine/threonine-protein kinase SBK1 | signaling | -6,93 | 0,00015683 |
| maker-ptg000253l-augustus-gene-58.60-mRNA-1 | *MYOCD* | smooth muscle cells and cardiac muscle cells-specific transcriptional factor | myocardin | transcription factor | -3,89 | 1,03704E-11 |
| maker-ptg000085l-snap-gene-26.0-mRNA-1 | *MYOG* | muscle-specific transcription factor | myogenin | transcription factor | -2,73 | 1,67587E-05 |
| maker-ptg000171l-augustus-gene-6.26-mRNA-1 | *ANO1* | calcium-activated chloride channel | anoctamin 1 | transmembrane ion transport | -3,40 | 1,225E-17 |
| maker-ptg000852l-snap-gene-16.100-mRNA-1 | *ATP1b3a* | Na/K-ATPase β-subunit | ATPase Na+/K+ transporting subunit beta 3a | transmembrane ion transport | -4,10 | 1,54943E-19 |
| maker-ptg000600l-augustus-gene-1.19-mRNA-1 | *ATP2a1* | sarcoplasmic/endoplasmic reticulum calcium ATPase 1 | sarcoplasmic/endoplasmic reticulum calcium ATPase 1 | transmembrane ion transport | -7,30 | 3,1585E-07 |
| maker-ptg000474l-snap-gene-6.56-mRNA-1 | *ATP2a2a* | sarcoplasmic/endoplasmic reticulum calcium ATPase 2 | sarcoplasmic/endoplasmic reticulum calcium ATPase 2a | transmembrane ion transport | -7,66 | 0,000724467 |
| maker-ptg000085l-snap-gene-5.119-mRNA-1 | *ATP2b2* | plasma membrane Ca2+ transporting ATPase 2 | ATPase plasma membrane Ca2+ transporting 2 | transmembrane ion transport | -2,46 | 0,002679005 |
| maker-ptg000718l-snap-gene-53.80-mRNA-1 | *ATP5f1b* | subunit of mitochondrial ATP synthase | ATP synthase subunit beta, mitochondrial | transmembrane ion transport | -1,81 | 2,19442E-06 |
| maker-ptg000237l-snap-gene-5.34-mRNA-1 | *CACNA1s* | voltage-gated calcium channel | dihydropyridine-sensitive L-type skeletal muscle calcium channel subunit alpha-1 | transmembrane ion transport | -6,87 | 3,16021E-05 |
| maker-ptg000085l-augustus-gene-14.68-mRNA-1 | *CACNA2d1* | voltage-gated calcium channel | voltage-dependent calcium channel subunit alpha-2/delta-2 | transmembrane ion transport | -4,46 | 1,86424E-11 |
| maker-ptg000202l-snap-gene-1.0-mRNA-1 | *CACNG1a* | voltage-gated calcium channel | voltage-dependent calcium channel gamma-1 subunit | transmembrane ion transport | -4,04 | 4,43468E-11 |

| | | | | | | |
|---|---|---|---|---|---|---|
| maker-ptg000763l-augustus-gene-3.165-mRNA-1 | *CALHM6* | pore-forming subunit of a voltage-gated ion channel | calcium homeostasis modulator family member 6 | transmembrane ion transport | -3,12 | 1,19758E-11 |
| maker-ptg001156l-augustus-gene-8.0-mRNA-1 | *CASQ1a* | calcium-binding protein in SR | calsequestrin-1a | transmembrane ion transport | -7,32 | 4,86051E-08 |
| maker-ptg000021l-augustus-gene-6.30-mRNA-1 | *CASQ1b* | calcium-binding protein in SR | calsequestrin-1b | transmembrane ion transport | -3,71 | 9,95212E-15 |
| maker-ptg000650l-augustus-gene-22.51-mRNA-1 | *CLCN1* | voltage-dependent chloride channel; important for repolarization of skeletal muscle cells after muscle contraction | chloride channel protein 1 | transmembrane ion transport | -7,56 | 6,86007E-07 |
| snap_masked-ptg000633l-processed-gene-25.11-mRNA-1 | *KCNA1b* | voltage-gated potassium channel | potassium voltage-gated channel subfamily A member 1b | transmembrane ion transport | -6,39 | 6,20958E-05 |
| snap_masked-ptg000643l-processed-gene-7.150-mRNA-1 | *KCNA4a* | voltage-gated potassium channel | potassium voltage-gated channel subfamily A member 4a | transmembrane ion transport | -4,05 | 0,000266439 |
| snap_masked-ptg000633l-processed-gene-25.9-mRNA-1 | *KCNA5b* | voltage-gated potassium channel | potassium voltage-gated channel subfamily A member 5b | transmembrane ion transport | -4,65 | 0,001218785 |
| snap_masked-ptg000633l-processed-gene-26.11-mRNA-1 | *KCNA6a* | voltage-gated potassium channel | potassium voltage-gated channel subfamily A member 6a | transmembrane ion transport | -5,40 | 2,72077E-12 |
| maker-ptg000600l-augustus-gene-13.28-mRNA-1 | *KCNA7b* | voltage-gated potassium channel | potassium voltage-gated channel subfamily A member 7b | transmembrane ion transport | -7,39 | 0,000445521 |
| maker-ptg000072l-snap-gene-0.6-mRNA-1 | *KCNAB2* | voltage-gated potassium channel | voltage-gated potassium channel subunit beta-2 | transmembrane ion transport | -3,37 | 9,48977E-05 |
| maker-ptg000068l-snap-gene-8.7-mRNA-1 | *KCNB1* | voltage-gated potassium channel | potassium voltage-gated channel subfamily B member 1 | transmembrane ion transport | -5,58 | 0,006990731 |
| maker-ptg001248l-est_gff_est2genome-gene-0.0-mRNA-1 | *KCNE4* | voltage-gated potassium channel | potassium voltage-gated channel subfamily E regulatory subunit 4 | transmembrane ion transport | -1,99 | 0,002653689 |
| maker-ptg000314l-augustus-gene-2.9-mRNA-1 | *KCNIP4* | voltage-gated potassium channel | Kv channel-interacting protein 4 | transmembrane ion transport | -3,20 | 0,000884639 |
| maker-ptg001348l-snap-gene-2.39-mRNA-1 | *KCNK4* | potassium two pore domain channel | potassium channel subfamily K member 4 | transmembrane ion transport | -4,94 | 0,00032563 |
| maker-ptg000170l-snap-gene-11.9-mRNA-1 | *KCNK7* | potassium two pore domain channel | potassium channel subfamily K member 1 | transmembrane ion transport | -5,11 | 9,18843E-12 |
| maker-ptg000253l-snap-gene-49.55-mRNA-1 | *KCNN4* | calcium-activated potassium channel | potassium calcium-activated channel subfamily N member 4 | transmembrane ion transport | -5,04 | 1,88707E-11 |
| maker-ptg000721l-snap-gene-3.11-mRNA-1 | *SCN3b* | voltage-gated sodium channel | sodium voltage-gated channel beta subunit 3 | transmembrane ion transport | -2,11 | 5,45393E-10 |
| maker-ptg000974l-augustus-gene-6.41-mRNA-1 | *SCN4ab* | voltage-gated sodium channel | sodium channel protein type 4 subunit alpha b | transmembrane ion transport | -5,38 | 4,17695E-08 |
| maker-ptg000422l-augustus-gene-0.23-mRNA-1 | *SRL* | Ca2+-binding protein in SR; Ca2+ buffering | sarcalumenin | transmembrane ion transport | -6,02 | 5,45506E-07 |
| maker-ptg000562l-augustus-gene-8.257-mRNA-1 | *TMEM38a* | monovalent cation channel in the SR and nuclear membranes of skeletal muscle | transmembrane protein 38A | transmembrane ion transport | -1,77 | 3,0994E-08 |
| maker-ptg000600l-snap-gene-7.69-mRNA-1 | *TRPM4* | Ca2 +-activated nonselective monovalent cation channel | transient receptor potential cation channel subfamily M member 4 | transmembrane ion transport | -4,84 | 1,11494E-06 |
| snap_masked-ptg002114l-processed-gene-2.64-mRNA-1 | *ABCC9* | subunit of ATP-sensitive potassium channels | ATP-binding cassette sub-family C member 9 | transmembrane ion transport | -3,10 | 3,68287E-09 |
| maker-ptg000181l-snap-gene-1.66-mRNA-1 | *SLC4a7* | sodium bicarbonate cotransporter | solute carrier family 4 member 7 | transmembrane ion transport | -4,78 | 1,00112E-09 |
| maker-ptg000361l-snap-gene-18.27-mRNA-1 | *SLC22a23* | antiporter to transport organic ions across cell membranes | solute carrier family 22 member 23 | transmembrane ion transport | -4,24 | 0,004015017 |
| maker-ptg000718l-augustus-gene-55.59-mRNA-1 | *SLC25a12* | mitochondrial electrogenic aspartate/glutamate antiporter | solute carrier family 25 member 12 | transmembrane ion transport | -3,53 | 1,18714E-12 |
| maker-ptg000230l-snap-gene-5.12-mRNA-1 | *SLC41a1* | Na+/Mg2+ ion exchanger | solute carrier family 41 member 1 | transmembrane ion transport | -4,91 | 5,94485E-08 |
| maker-ptg000600l-augustus-gene-10.17-mRNA-1 | *SLC6a16* | Na(+)- and Cl(-)-dependent neurotransmitter transporter | solute carrier family 6 member 16 | transmembrane ion transport | -2,21 | 1,60618E-05 |
| maker-ptg000299l-snap-gene-9.30-mRNA-1 | *SLC6a6b* | taurine:sodium symporter | solute carrier family 6 member 6b | transmembrane ion transport | -2,58 | 1,66873E-11 |

**Supplementary Table 3** 44 Significantly enriched Gene Ontology terms with Fisher's exact test p-value < 0.01 in genes  up-regulated in electric organ.

| Term | GO terms | Category | Count | % | P-value | Genes | List Total | Pop Hits | Pop Total | Fold Enrichment | Bonferroni | Benjamini | FDR |
|---|---|---|---|---|---|---|---|---|---|---|---|---|---|
| GO:0016310 | phosphorylation | Biological Process | 61 | 5,407801 | 1,32E-04 | *RET, SI:CH211-195B13.1, PFKFB2B, MOB1BA, PIK3CG, HK2, STK24B, RPS6KA3A, RPS6KA2, PIP5KL1, RPS6KA1, RPS6KB1B, AKT1, SI:CH211-243J20.2, PIM3, ERBB4B, PDGFRA, PRKCI, MAP4K3A, PRKCBB, DCLK1A, SI:DKEY-172J4.3, EPHA4A, SHPK, ACVR1BB, MAPK14A, PI4K2A, GNE, IPPK, ITPK1B, UCK2A, LTK, PRKX, TTK, CITA, PIP4K2AA, PRKCZ, PFKPA, CKMT1, PIP5K1CA, GRK6, ERBB2, TRIOA, CAMK1GB, MAPK6, CHKB, YES1, TNK2B, LIMK2, PTK2AB, HIPK2, PRKACBB, ETNK2, JAK2B, RPS6KAL, PRKCHA, CMPK, PI4KB, PTK7B, CDK14, CAMK1DA* | 944 | 718 | 18397 | 1,66 | 0,201384509 | 0,224860832 | 0,224860832 |
| GO:0018105 | peptidyl-serine phosphorylation | Biological Process | 16 | 1,41844 | 0,001444283 | *PRKCI, SI:CH211-195B13.1, TTK, PRKX, TTBK1A, PRKCZ, HIPK2, PRKCBB, RPS6KA3A, RPS6KA2, RPS6KA1, RPS6KAL, RPS6KB1B, PRKCHA, CAMK1GB, CAMK1DA* | 944 | 122 | 18397 | 2,56 | 0,914560516 | 0,684527812 | 0,684527812 |
| GO:0098609 | cell-cell adhesion | Biological Process | 19 | 1,684397 | 0,001444379 | *DCHS1A, ARNT2, NEO1A, IGSF9BA, ITGA2B, CTNND1, NRCAMA, HSPG2, TMEM47, CNTN1A, DLG2, PERP, ITGA8, CDH13, ELMO2, ITGAV, PKP4, FAT4, CDH17* | 944 | 160 | 18397 | 2,31 | 0,91457447 | 0,684527812 | 0,684527812 |
| GO:0001944 | vasculature development | Biological Process | 12 | 1,06383 | 0,003025118 | *NOTCH2, YAP1, STN1, PANK2, ANGPT2B, CYP26C1, LGALS2A, AGAP2, ENPP2, ITGB8, RAB11A, MYCA* | 944 | 82 | 18397 | 2,85 | 0,994238484 | 0,684527812 | 0,684527812 |
| GO:0043123 | positive regulation of I-kappaB kinase/NF-kappaB signaling | Biological Process | 8 | 0,70922 | 0,003315885 | *PRKCBB, CD40, TNIP2, TNFRSF19, RBCK1, S100B, LURAP1, MAP3K14A* | 944 | 39 | 18397 | 4,00 | 0,996493066 | 0,684527812 | 0,684527812 |
| GO:0001935 | endothelial cell proliferation | Biological Process | 5 | 0,443262 | 0,003380098 | *ITGA2B, ITGB8, ITGAV, ARHGEF7B, HSPG2* | 944 | 13 | 18397 | 7,50 | 0,996857294 | 0,684527812 | 0,684527812 |
| GO:0030198 | extracellular matrix organization | Biological Process | 16 | 1,41844 | 0,003386468 | *MMP15B, MMP17A, FBLN2, COLQ, SMOC2, COL4A2, ADAMTSL4, SI:DKEY-6N6.1, COL1A1A, ADAMTSL3, ADAMTS17, MMP28, COL2A1B, MMP19, ADAMTSL7, ADAMTS9* | 944 | 133 | 18397 | 2,34 | 0,996891298 | 0,684527812 | 0,684527812 |
| GO:0072583 | clathrin-dependent endocytosis | Biological Process | 6 | 0,531915 | 0,003582283 | *AP2M1A, FCHO1, DNAJC6, SGIP1A, AP2A1, GPR107* | 944 | 21 | 18397 | 5,57 | 0,997774987 | 0,684527812 | 0,684527812 |
| GO:0070121 | Kupffer's vesicle development | Biological Process | 12 | 1,06383 | 0,003658342 | *YAP1, MBD3B, SNX10A, VGLL4B, ARL6, RAB3IP, DNMT3BB.1, ENPP2, ITGAV, GPR22A, RAB11A, ATP6V1F* | 944 | 84 | 18397 | 2,78 | 0,998046074 | 0,684527812 | 0,684527812 |
| GO:0061817 | endoplasmic reticulum-plasma membrane tethering | Biological Process | 4 | 0,35461 | 0,004021903 | *ESYT1A, ESYT2B, ESYT2A, GRAMD2AA* | 944 | 7 | 18397 | 11,14 | 0,998950121 | 0,684527812 | 0,684527812 |
| GO:0006486 | protein glycosylation | Biological Process | 16 | 1,41844 | 0,006246214 | *GALNT12, ST6GAL1, GALNT13, GALNT16, B3GAT2, EXT1B, FUT8A, ST6GALNAC5A, MGAT1B, B3GNT7, ST3GAL4, ST8SIA5, ST8SIA6, STT3B, LARGE2, ST3GAL2* | 944 | 142 | 18397 | 2,20 | 0,999976638 | 0,804864816 | 0,804864816 |
| GO:0043409 | negative regulation of MAPK cascade | Biological Process | 6 | 0,531915 | 0,006588046 | *DUSP4, DUSP5, SPRED2B, PPEF2A, SPRED2A, DUSP7* | 944 | 24 | 18397 | 4,87 | 0,999986992 | 0,804864816 | 0,804864816 |
| GO:0006811 | ion transport | Biological Process | 47 | 4,166667 | 0,006643522 | *SLC24A2, GLRBB, KCNG3, SCN4BA, SCN1BA, TTYH2, PACC1, TMEM63C, ITPR1B, ATP2C1, ATP1A3A, CHRNG, ATP1B1B, ATP2A2B, KCNQ5B, SLC39A7, GABRD, ATP1A1A.4, SI:CH211-225P5.8, ATP6V1F, SLC13A1, SCN4AA, HEPHL1B, GABRA1, CHRNB4, SLC8A1A, KCNJ9, SLC11A2, SLC39A10, TRPV1, SI:DKEY-28B4.8, CNGA3A, GRIN2AA, SLC5A9, ATP6V0A1A, KCNJ2A, KCNK2A, SLCO4A1, ATP1A2A, ATP2B1A, ATP2B3B, CACNA1BA, VDAC1, SLC4A4A, SLC4A4B, MCU, GRIA3B* | 944 | 614 | 18397 | 1,49 | 0,999988172 | 0,804864816 | 0,804864816 |
| GO:0030030 | cell projection organization | Biological Process | 11 | 0,975177 | 0,006953193 | *CATIP, SNX10A, INTU, ARL6, CFL1, IFT122, TMEM237A, GPR22A, CCDC61, GSNA, SDCCAG8* | 944 | 79 | 18397 | 2,71 | 0,999993042 | 0,804864816 | 0,804864816 |
| GO:0000188 | inactivation of MAPK activity | Biological Process | 5 | 0,443262 | 0,007609678 | *DUSP4, DUSP5, SPRED2B, SPRED2A, DUSP7* | 944 | 16 | 18397 | 6,09 | 0,999997742 | 0,804864816 | 0,804864816 |
| GO:0048593 | camera-type eye morphogenesis | Biological Process | 5 | 0,443262 | 0,007609678 | *SOX11A, LAMA1, LUM, ALDH1A2, IFT122* | 944 | 16 | 18397 | 6,09 | 0,999997742 | 0,804864816 | 0,804864816 |
| GO:0007264 | small GTPase mediated signal transduction | Biological Process | 13 | 1,152482 | 0,00803919 | *RAC3A, BCAR3, ARHGAP32A, RHOUB, DOCK4B, GDI1, DOCK7, TIAM1B, RHOF, RAPGEFL1, RHOBTB1, RHOCB, RASGEF1BA* | 944 | 106 | 18397 | 2,39 | 0,999998919 | 0,804864816 | 0,804864816 |
| GO:0006790 | sulfur compound metabolic process | Biological Process | 4 | 0,35461 | 0,008935331 | *CHST6, CHST7, CHST2B, SI:CH73-62B13.1* | 944 | 9 | 18397 | 8,66 | 0,999999768 | 0,844885165 | 0,844885165 |
| GO:0016020 | membrane | Cellular Component | 464 | 41,13475 | 3,42E-10 | *SI:DKEY-34D22.1, PGAP2, TMEM200A, OLFCS1, EXT1B, CORO2BA, ZFYVE28, LAMB1A, NSDHL, CHST2B, TIAM1B, BCR, BCAM, SLC5A9, GRAMD1BA, CACNA1BA, MCOLN1A, PI4K2A, GRAMD1BB, FREM3, CDC42SE1, RPN2, TTYH2, SGIP1A, ABCB5, MBOAT2B, HACD3, SI:DKEY-91M11.5, PDGFC, CHST10, ANO5B, ST3GAL4, GSG1L2B, ST3GAL2, PLXNB1B, PLXNB1A, ABCC6A, PRRT1, ABCA2, CADM3, ICMT, APCDD1L, SI:DKEY-15H8.17, SI:CH211-286O17.1, EIF5, NPTNB, MCOLN3A, TMEM237A, NMT2, SPIRE1A, PAM, RALAA, RET, ACHE, MTMR10, MTMR11, TMEM181, LYST, SLC6A2, PHEX, CYB561D2, TMEM47, CATIP, ADAMTSL3, SH3GLB2B, ATP1A1A.4, TMEM119B, CHST6, CHST7, RHBDF1A, ELOVL2, SLC39A10, GNL1, PRSS12, FAM234B, ECRG4A, SLCO4A1, ACVR1BB, TSPAN7B, ERGIC3, PIGF, MGLL, PACC1, SLC16A6B, TMEM72, ADCY7, PPP1R3AA, CHRNG, GDPD4A, MUC13B, SLC17A5, SC5D, GABRA1, SNX21, PLXDC1, TSPAN9A, GDPD5B, SH3GLB1A, CYB561, RIC3B, RCA2.1, TMEM229B, GPR146, SEMA5A, ADCY1A, GLRBB, KCNG3, NRROS, TUSC3, CLSTN1, ITGA2B, CXCL14, KDELR2B, AP2M1A, ABHD12, LAPTM4B, RNF19A, CFL1, TSPAN4A, SI:CH211-1E14.1, SI:DKEY-122A22.2, PLXNC1, SLC39A7, PHLDA2, IL13RA2, LARGE2, PHLDA3, SEMA6A, SLC25A23B, BSCL2, FRMD3, TOM1L2, TRIM101, MAG, MFSD12A, SYPL1, MADD, ROR2, SI:CH73-62B13.1, GRIA3B, NOTCH2, ABHD2A, ARL6, TMEM230A, LIN7C, PFKPA, MGAT1B, ADTRP1, SI:CH211-76L23.7, SI:CH211-225P5.8, DRD4B, KL, SLC37A2, OSBPL6, RAB4B, SDF2L1, GPR22A, SLC25A25B, CYP51, TMPRSS15, GRIN2AA, CNGA3A, G6PC3, TSPAN5A, CLTCB, GPR107, MCU, SPRED2B, DYSF, NENF, MRC1A, PIK3CG, B4GALNT3A, SPRED2A, SNX10A, SI:CH73-364H19.1, NAPBB, CERS3A, ITGB8, ITGAV, IL21R.1, CCDC51, SLC13A1, ADGRV1, CERS6, ZGC:110329, CHPF2, TNFRSF19, NRG1, SLC25A55A, ST6GALNAC5A, RRBP1A, SYNGR3A, PRKCBB, ZDHHC8B, KCNJ2A, SLC15A1A, ITGA8, EFHC1, TNFRSF21, CERS1, SLC24A2, NLGN1, NEO1A, ATP10D, LRP5, JAKMIP3, AP3M2, CKMT1, FCHO1, CYB5A, LMBRD2B, KCNJ9, SEMA4C, MBOAT1, MCOLN2, SI:DKEY-11F4.7, TMPPE, FRAS1, NECAP1, CERS2B, KCNK2A, DOLPP1, MGAT4A, CNGB1A, TRIM36, SLC4A4A, SLC4A4B, PTPN5, CNNM2B, SLC23A2, SLC35B4, ZGC:165507, SCN1BA, PLEKHB2, ITPR1B, TXNDC11, FRMD4BA, ZDHHC4, NPDC1A, STS, ERC1B, SLITRK3A, EHD1B, MFSD2B, ADRA1AA, ESYT2B, ESYT2A, ENTPD1, ST6GAL1, SLC6A17, SLC11A2, MMP15B, ADGRA3, ATP1A2A, EPHA4A, ATP2B1A, VDAC1, PEX11G, LRP1BB, CDS1, ZNF106A, ESYT1A, TMEM63C, RRAD, PEX11A, MYO6A, PCDH19, CRHR1, CD79B, LOXL3B, ATP1B1B, ST8SIA5, ST8SIA6, ANO10B, RHOCB, CARMIL2, TMEM86A, TMEM184C, YES1, CARMIL3, VASNB, NRXN3A, MACF1B, FLRT1B, AVPR1AA, NCAM1A, BAMBIA, MACF1A, AHCTF1, CD40, TENM4, AP2A1, SPPL3, ZCCHC14, ATP2A2B, SMPD2B, QSOX1, ERBB4B, DCHS1B, SI:CH211-264F5.2, DCHS1A, ZDHHC13, TMEM30AB, ANO6, MMEL1, ZDHHC17, CSGALNACT1A, ZDHHC14, SREBF2, RGMA, B3GNT7, AVPR2AA, GRINAA, SLC5A7A, SORBS2A, RPRMA, ZGC:92045, AMIGO1, HSD17B3, RFNG, NRN1A, GUCY2F, ATP1A3A, ORMDL2, PERP, ZGC:86609, MEP1A.1, REEP3B, KCNQ5B, GABRD, ZGC:162698, MVB12BA, ZGC:92275, YIPF5, STAC3, SCDB, ZDHHC23B, PLEKHA8, DPY19L3, ZDHHC9, ATP2B3B, EPPK1, ZGC:63972, SLC6A6B, GALNT12, SH3GL2B, GRM6A, GALNT13, GALNT16, ACSL4A, COX6A1, TMEM263, SELENOU1A, PALMDA, FAM174B, ANK3B, IGFLR1, SBF1, ADAM19B, PDGFRA, CHRNB4, PCSK5B, SLC8A1A, CCR12A, IGSF9BA, APLP2, SRD5A2A, CYBA, SHISA3, ACYP1, FUT8A, KCTD7, ALDH3A1, COL4A2, B3GLCTA, TMCC1B, FAM20CB, ABCG1, AQP1A.1, DIPK1B, DDX5, SCARB2A, PSEN2, LDLRB, SI:CH211-241B2.5, TMEM242, MAN2A2, CLDN11A, ATP6V1F, SI:DKEY-32E23.4, MFAP3L, OPN3, CAV3, ACSL3B, CRIM1, TRPV1, PLECA, IL17REL, DLG2, JAK2B, BAXA, CAPRIN2, PI4KB, SDK1A, STT3B, PTK7B, FLOT2A, NRCAMA, SI:CH73-269M14.2, HK2, SEC61A1, GRM4, SI:DKEY-112M2.1, CPT2, SI:DKEY-11F4.16, GRAMD2AA, HGSNAT, EGFL6, SI:CH211-153B23.3, LMF2A, ATP6AP2, BRINP3A.1, SI:DKEY-28B4.8, TMIGD1, ATP6V0A1A, CPTP, TMEM218, CDH13, NAT8L, CDH17, RNF128A, VAC14, LTK, SCN4BA, LAMA1, MOSPD2, PMP22B, ABHD17C, ATP2C1, CLN3, CNTN1A, RASD1, CHSY1, GNG5, SPOCK3, ERBB2, CYP4T8, SCN4AA, TMEM54A, TNK2B, LAMB2, BNIP3, B3GAT2, ACKR4A, PNKD, NDST1B, ABCG2A, AQP11, KRAS, CHMP6B, PCDH1G31, CADM2B* | 983 | 7112 | 18868 | 1,25 | 1,30E-07 | 1,30E-07 | 1,27E-07 |
| GO:0005886 | plasma membrane | Cellular Component | 237 | 21,01064 | 6,08E-08 | *ZGC:165507, SI:DKEY-34D22.1, SCN1BA, CPNE7, OLFCS1, ITPR1B, RAPGEFL1, ZDHHC4, EFR3A, MFSD2B, EHD1B, ESYT2B, ESYT2A, ENTPD1, PRKCI, SLC6A17, ACTN1, SLC11A2, TIAM1B, ADGRA3, HSPG2, BCAM, SLC5A9, CCNY, PRKAR1B, ATP1A2A, EPHA4A, ATP2B1A, GRAMD1BA, MCOLN1A, VDAC1, PI4K2A, GRAMD1BB, LRP1BB, CDC42SE1, ESYT1A, TTYH2, RRAD, TMEM63C, SGIP1A, MYO6A, PCDH19, CRHR1, CD79B, ATP1B1B, ANO5B, GSG1L2B, STXBP6, ANO10B, RHOCB, PLXNB1B, PLXNB1A, CARMIL2, YES1, CARMIL3, VASNB, APCDD1L, NPTNB, MCOLN3A, AVPR1AA, BAMBIA, FAT4, SPIRE1A, RALAA, RET, ACHE, DOCK4B, TENM4, SLC6A2, PHEX, CATIP, ERBB4B, ATP1A1A.4, TMEM119B, DCHS1B, HEPHL1B, SI:CH211-264F5.2, DCHS1A, SLC39A10, TMEM30AB, ANO6, MMEL1, RHOF, GMIP, PRSS12, ZDHHC17, RGMA, ECRG4A, SLCO4A1, ACVR1BB, AVPR2AA, TSPAN7B, DSCAML1, SLC5A7A, SORBS2A, EPS15L1A, SLC16A6B, PACC1, AMIGO1, NRN1A, GUCY2F, ADCY7, SEMA3AB, ATP1A3A, CHRNG, SLC17A5, ARF3A, GABRD, RASGEF1BA, SCCPDHA.1, GABRA1, STAC3, GDPD5B, TSPAN9A, ATP2B3B, SLC6A6B, RIC3B, UNC13BB, ADCY1A, TRHDE.1, GLRBB, GRM6A, NRROS, CTNND1, ACSL4A, AP2M1A, RAB44, LAPTM4B, FAM174B, IGFLR1, TSPAN4A, PLXNC1, PHLDA3, PDGFRA, CHRNB4, SLC8A1A, LMTK2, CYBA, TTC7A, KCTD7, MAG, MFSD12A, PSTPIP1A, PKP4, ROR2, ABCG1, GRIA3B, NOTCH2, ABHD2A, SCARB2A, ARL6, PSEN2, LDLRB, PIP4K2AA, LIN7C, RHOBTB1, PIP5K1CA, GRK6, CLDN11A, CNRIP1B, ATP6V1F, MFAP3L, DRD4B, OSBPL6, RAB4B, OPN3, CAV3, STXBP1A, ACSL3B, CRIM1, TRPV1, GPR22A, GRIN2AA, CNGA3A, DLG2, CAPRIN2, TSPAN5A, FLOT2A, HSP90AB1, SPRED2B, DYSF, NRCAMA, TRH, MRC1A, PIK3CG, RERG, GRM4, SEMA3GB, PIP5KL1, ADGRV1, ZGC:110329, TNFRSF19, ATP6AP2, ATP6V0A1A, KCNJ2A, NID1A, CPTP, SLC15A1A, CDH13, CDH17, TNFRSF21, SLC24A2, RAC3A, NLGN1, LTK, NEO1A, SCN4BA, ATP10D, LRP5, PMP22B, ABHD17C, ATP2C1, CNTN1A, GNG5, FCHO1, ERBB2, SH2B2, SI:DKEY-206P8.1, SCN4AA, PDZD7A, LMBRD2B, TNK2B, MYO1EA, KCNJ9, SEMA4C, MCOLN2, SI:DKEY-11F4.7, EEPD1, KCNK2A, CNGB1A, ABCG2A, SLC4A4A, KRAS, RGS11, SLC4A4B, PCDH1G31, CNNM2B* | 983 | 3299 | 18868 | 1,38 | 2,30E-05 | 1,15E-05 | 1,12E-05 |
| GO:0005794 | Golgi apparatus | Cellular Component | 62 | 5,496454 | 2,02E-06 | *GALNT12, SLC35B4, GALNT13, GALNT16, PGAP2, CLSTN1, EXT1B, B4GALNT3A, ZDHHC4, RAB44, STK24B, FAM174B, SLC39A7, LARGE2, ADAM19B, PDGFRA, ST6GAL1, RHBDF1A, CHPF2, ZDHHC13, TMEM30AB, FUT8A, ZDHHC17, ST6GALNAC5A, ZDHHC14, CSGALNACT1A, SREBF2, ZDHHC8B, B3GNT7, CPTP, ZGC:162200, ERGIC3, FAM20CB, GRINAA, PI4K2A, RNF128A, TMEM230A, MYO6A, PSEN2, RFNG, TMEM241, AP3M2, MGAT1B, CHSY1, CHST10, ST3GAL4, ST8SIA5, ST8SIA6, ZGC:162698, ST3GAL2, ARHGAP32A, CAV3, B3GAT2, YIPF5, CLASP1A, ZDHHC23B, RAB11A, PLEKHA8, ZDHHC9, NDST1B, SH3GLB1A, GPR107* | 983 | 628 | 18868 | 1,89 | 7,65E-04 | 2,55E-04 | 2,49E-04 |
| GO:0005783 | endoplasmic reticulum | Cellular Component | 68 | 6,028369 | 1,89E-05 | *NRROS, PGAP2, EXT1B, ITPR1B, ACSL4A, TXNDC12, PITPNBL, ZDHHC4, CYB561D2, KDELR2B, ABHD12, SEC61A1, NSDHL, CERS3A, SI:DKEY-122A22.2, SMPD2B, SLC39A7, ESYT2B, ESYT2A, CERS6, RHBDF1A, ELOVL2, ATP6AP2, LMF2A, BRINP3A.1, SRD5A2A, TMEM30AB, PDIA8, ZDHHC14, SREBF2, PDIA4, BSCL2, RCN2, ERGIC3, CRELD2, NAT8L, GRINAA, RNF128A, CERS1, NECAB3, PDXDC1, RPN2, PSEN2, HSD17B3, ATP2C1, HACD3, ORMDL2, REEP3B, CALR3A, KL, SLC37A2, ICMT, SDF2L1, BNIP3, YIPF5, ACSL3B, SERPINH1B, NCK2B, ZDHHC23B, CYP51, G6PC3, TBL2, CERS2B, ZDHHC9, MGAT4A, DOLPP1, PI4KB, RIC3B* | 983 | 763 | 18868 | 1,71 | 0,007155957 | 0,001795404 | 0,001752769 |
| GO:0000139 | Golgi membrane | Cellular Component | 35 | 3,102837 | 1,26E-04 | *GALNT12, SLC35B4, GALNT13, GALNT16, PGAP2, CLSTN1, PSEN2, RFNG, ATP2C1, ZDHHC4, KDELR2B, MAN2A2, CHST10, QSOX1, LARGE2, PCSK5B, CHST6,* | 983 | 331 | 18868 | 2,03 | 0,04670409 | 0,00956538 | 0,009338234 |

| | | | | | | | | | | | | | |
|---|---|---|---|---|---|---|---|---|---|---|---|---|---|
| GO:0005789 | endoplasmic reticulum membrane | Cellular Component | 48 | 4,255319 | 2,51E-04 | CDS1, ESYT1A, SI:DKEY-13N15.2, DIPK1B, NRROS, SLC35B4, PGAP2, EXT1B, PSEN2, FMN2B, ITPR1B, HACD3, ZDHHC4, KDELR2B, ABHD12, SEC61A1, CERS3A, ORMDL2, REEP3B, SC5D, SLC39A7, CALR3A, ESYT2B, ESYT2A, SLC37A2, ICMT, CERS6, RHBDF1A, SDF2L1, ELOVL2, LMF2A, ATP6AP2, YIPF5, SCDB, ZDHHC14, SREBF2, RRBP1A, CYP51, BSCL2, G6PC3, ZDHHC9, CERS2B, DOLPP1, GRAMD1BA, ERGIC3, SEC24D, PIGF, GRAMD1BB | 983 | 529 | 18868 | 1,74 | 0,090736226 | 0,015851352 | 0,015474934 |
| GO:0016021 | integral component of membrane | Cellular Component | 309 | 27,39362 | 0,00231972 | SLC23A2, SLC35B4, ZGC:165507, PGAP2, SI:DKEY-34D22.1, PLEKHB2, TMEM200A, EXT1B, TXNDC11, ITPR1B, FRMD4BA, ZDHHC4, NPDC1A, NSDHL, STS, SLITRK3A, ARL10, ESYT2B, ESYT2A, ENTPD1, ST6GAL1, CHST2B, SLC11A2, MMP15B, ADGRA3, BCAM, SLC5A9, ATP1A2A, EPHA4A, GRAMD1BA, MCOLN1A, GRAMD1BB, FREM3, LRP1BB, CDS1, ESYT1A, RPN2, TMEM63C, SGIP1A, ABCB5, MBOAT2B, PCDH19, HACD3, CRHR1, CD79B, CHST10, PDGFC, ANO5B, ST3GAL4, ST8SIA5, ST8SIA6, GSG1L2B, ANO10B, ST3GAL2, PRRT1, ABCC6A, ABCA2, CADM3, TMEM86A, TMEM184C, ICMT, VASNB, APCDD1L, SI:DKEY-15H8.17, NRXN3A, SI:CH211-286O17.1, FLRT1B, EIF5, GPR180, NPTNB, MCOLN3A, TMEM237A, NCAM1A, FAT4, PAM, AHCTF1, TENM4, TMEM181, PHEX, CYB561D2, ZCCHC14, TMEM47, ADAMTSL3, ATP2A2B, SMPD2B, ERBB4B, ATP1A1A.4, TMEM119B, SI:CH211-264F5.2, DCHS1A, CHST6, RHBDF1A, ZDHHC13, SLC39A10, TMEM30AB, ANO6, MMEL1, GNL1, ZDHHC17, CSGALNACT1A, ZDHHC14, SREBF2, RGMA, FAM234B, B3GNT7, SLCO4A1, CRELD1, ERGIC3, GRINAA, DSCAML1, PIGF, SLC5A7A, RPRMA, ZGC:92045, SLC16A6B, PACC1, HSD17B3, TMEM72, GUCY2F, PPP1R3AA, TMEM164, MEGF6A, ATP1A3A, GDPD4A, ORMDL2, PERP, MEP1A.1, MUC13B, REEP3B, SLC17A5, KCNQ5B, SC5D, ZGC:162698, ZGC:92275, YIPF5, KLHL2, ZDHHC23B, PLXDC1, DPY19L3, ZDHHC9, GDPD5B, CYB561, RCA2.1, RIC3B, TMEM229B, GPR146, GALNT12, SEMA5A, TRHDE.1, KCNG3, GALNT13, NRROS, GALNT16, TUSC3, CLSTN1, ITGA2B, ACSL4A, COX6A1, TMEM263, KDELR2B, ABHD12, LAPTM4B, TMEM268, FAM174B, RNF19A, IGFLR1, SI:CH211-1E14.1, SI:DKEY-122A22.2, SLC39A7, IL13RA2, LARGE2, ADAM19B, PDGFRA, SLC8A1A, SEMA6A, CCR12A, IGSF9BA, APLP2, SRD5A2A, SHISA3, ACYP1, SLC25A23B, FUT8A, FRMD3, TOM1L2, ALDH3A1, TRIM101, MAG, B3GLCTA, SYPL1, TMCC1B, FAM20CB, ABCG1, AQP1A.1, NOTCH2, DDX5, ABHD2A, DIPK1B, SCARB2A, TMEM230A, PSEN2, SI:CH211-241B2.5, LDLRB, TMEM241, TMEM242, ADTRP1, MGAT1B, MAN2A2, CLDN11A, MFAP3L, KL, SLC37A2, ACSL3B, CRIM1, PLECA, GPR22A, SLC25A25B, CYP51, TMPRSS15, PTPRD, CNGA3A, IL17REL, G6PC3, BAXA, GPR107, SDK1A, STT3B, PTK7B, MCU, PRR7, MTCL1, DYSF, NRCAMA, SI:CH73-269M14.2, MRC1A, HK2, B4GALNT3A, GRM4, SI:DKEY-112M2.1, SI:CH73-364H19.1, CERS3A, SI:DKEY-11F4.16, IL21R.1, CCDC51, GRAMD2AA, SLC13A1, HGSNAT, CERS6, ADGRV1, CHPF2, TNFRSF19, SI:CH211-153B23.3, ATP6AP2, LMF2A, BRINP3A.1, NRG1, SLC25A55A, ST6GALNAC5A, SYNGR3A, ZDHHC8B, SI:DKEY-28B4.8, TMIGD1, ATP6V0A1A, KCNJ2A, TMEM218, EFHC1, NAT8L, CDH17, RNF128A, CERS1, NEO1A, SCN4BA, ATP10D, LRP5, MOSPD2, PMP22B, ATP2C1, JAKMIP3, CLN3, CKMT1, CHSY1, ERBB2, CYP4T8, TMEM54A, CYB5A, LMBRD2B, KCNJ9, B3GAT2, BNIP3, ACKR4A, MBOAT1, MCOLN2, SI:DKEY-11F4.7, TMPPE, PNKD, FRAS1, CERS2B, MGAT4A, NDST1B, CNGB1A, AQP11, TRIM36, SI:CH211-152P11.8, SLC4A4A, SLC4A4B, PTPN5, CNNM2B, CADM2B | 983 | 5161 | 18868 | 1,15 | 0,585297993 | 0,125596253 | 0,122613757 |
| GO:0005905 | clathrin-coated pit | Cellular Component | 7 | 0,620567 | 0,003965195 | AP2M1A, HIP1RB, FCHO1, SGIP1A, MYO6A, CLTCB, AP2A1 | 983 | 30 | 18868 | 4,48 | 0,778159411 | 0,187851107 | 0,183390263 |
| GO:0031012 | extracellular matrix | Cellular Component | 22 | 1,950355 | 0,004951409 | NRROS, VASNB, MMP15B, MMP17A, TSKU, COLQ, VCANB, CCN4A, COL4A2, ADAMTSL4, SI:DKEY-6N6.1, COL1A1A, ADAMTSL3, ADAMTS17, SI:CH211-106H11.3, MMP28, COL2A1B, TIMP2A, MMP19, SI:DKEY-65B12.6, ADAMTSL7, ADAMTS9 | 983 | 218 | 18868 | 1,94 | 0,84759982 | 0,208509355 | 0,203557945 |
| GO:0030424 | axon | Cellular Component | 19 | 1,684397 | 0,005735502 | RET, SCN4AA, SYNM, SLC8A1A, SCN1BA, CLSTN1, ATP6AP2, ROGDI, ELAVL4, CCKA, NRCAMA, RAB11A, SI:DKEY-91M11.5, BCR, CNTN1A, DLG2, NPTNB, DSCAML1, SLC5A7A | 983 | 179 | 18868 | 2,04 | 0,886959935 | 0,217375523 | 0,212213572 |
| GO:0042995 | cell projection | Cellular Component | 31 | 2,748227 | 0,007956686 | RAC3A, TENM4, INTU, ARL6, CLSTN1, ROGDI, ELAVL4, MYO6A, ABHD17C, RHOBTB1, SI:DKEY-91M11.5, LAPTM4B, TEKT3, RHOCB, ACTR3, PDGFRA, ADGRV1, ACTN1, IFT122, PTK2AB, RHOF, ZDHHC17, BCR, DLG2, TMEM218, TMEM237A, PI4K2A, GSNA, CCDC61, MACF1A, SDCCAG8 | 983 | 361 | 18868 | 1,65 | 0,951570454 | 0,252320114 | 0,246328344 |
| GO:0031410 | cytoplasmic vesicle | Cellular Component | 24 | 2,12766 | 0,008805908 | RAC3A, SI:DKEY-13N15.2, DENND4C, ROGDI, SPRED2B, MYO6A, DYSF, ZDHHC13, RHOF, ZDHHC17, SREBF2, RHOBTB1, KDELR2B, AP3M2, CADPSA, AP2M1A, TBC1D7, CLTCB, SPIRE1A, SEC24D, FLOT2A, RHOCB, CYB561, PI4K2A | 983 | 259 | 18868 | 1,78 | 0,964993461 | 0,252320114 | 0,246328344 |
| GO:0031234 | extrinsic component of cytoplasmic side of plasma membrane | Cellular Component | 8 | 0,70922 | 0,009171801 | ESYT1A, ESYT2B, ESYT2A, YES1, TNK2B, STAC3, PTK2AB, GRAMD2AA | 983 | 46 | 18868 | 3,34 | 0,969564691 | 0,252320114 | 0,246328344 |
| GO:0031227 | intrinsic component of endoplasmic reticulum membrane | Cellular Component | 4 | 0,35461 | 0,009320532 | ESYT1A, ESYT2B, ESYT2A, GRAMD2AA | 983 | 9 | 18868 | 8,53 | 0,97124797 | 0,252320114 | 0,246328344 |
| GO:0016740 | transferase activity | Molecular Function | 126 | 11,17021 | 3,52E-05 | GALNT12, GALNT13, PRDM9, GALNT16, PFKFB2B, GTF2B, EXT1B, ZDHHC4, RPS6KA3A, RPS6KA2, RNF19A, RPS6KA1, AKT1, PIM3, LARGE2, PDGFRA, PRKCI, ST6GAL1, CHST2B, UBE2E3, MAP4K3A, FUT8A, RC3H2, DCLK1A, SI:CH211-256M1.8, B3GLCTA, EPHA4A, MAPK14A, RBCK1, PI4K2A, SI:CH73-62B13.1, GNE, CDS1, IPPK, UCK2A, ITPK1B, CRATA, UAP1, CITA, PIP4K2AA, PRKCZ, PFKPA, MGAT1B, PIP5K1CA, GRK6, CHST10, ST3GAL4, ST8SIA5, ST8SIA6, TRIOA, CAMK1GB, ST3GAL2, YES1, ICMT, CHKB, JAK2B, PRKCHA, NMT2, PI4KB, STT3B, PTK7B, CDK14, RET, PYGB, SI:CH211-195B13.1, MOB1BA, PIK3CG, HK2, B4GALNT3A, GYS1, STK24B, CPT2, CERS3A, PIP5KL1, RPS6KB1B, SI:CH211-243J20.2, ERBB4B, CHST6, CHST7, CERS6, CHPF2, ELOVL2, ZDHHC17, ZDHHC14, CSGALNACT1A, ST6GALNAC5A, APRT, PRKCBB, ZDHHC8B, KAT2A, MYLIPA, B3GNT7, UBE2R2, SI:DKEY-172J4.3, ACVR1BB, NAT8L, RNF213A, KMT2CA, LTK, UHRF1, PRKX, TTK, RFNG, METTL21A, CKMT1, CHSY1, ERBB2, MAPK6, RNF20, EHMT1B, TNK2B, B3GAT2, LIMK2, DNMT3BB.1, PTK2AB, ZDHHC23B, HIPK2, CERS2B, PRKACBB, ZDHHC9, ETNK2, BMT2, NDST1B, RPS6KAL, CMPK, CAMK1DA | 880 | 1743 | 17340 | 1,42 | 0,027705485 | 0,014324224 | 0,014288368 |
| GO:0005509 | calcium ion binding | Molecular Function | 64 | 5,673759 | 3,59E-05 | RET, SNED1, CAPN1A, CAPN1B, FKBP14, CETN2, PDCD6, CLSTN1, CETN3, DYSF, ANXA11B, ITPR1B, EFHD2, ENPP2, EFHD1, EHD1B, DCHS1B, ESYT2B, ESYT2A, DCHS1A, EGFL6, ACTN1, ANXA4, VWDE, SLC25A23B, HSPG2, MYL4, VCANB, RCN2, NID1A, CDH13, CRELD1, PPEF2A, CRELD2, CDH17, GSNA, LRP1BB, FBLN7, ESYT1A, NOTCH2, EPS15L1A, DIPK1B, NECAB3, SWAP70B, LDLRB, PCDH19, FBLN2, MEGF6A, SPOCK3, REPS2, SLIT2, CALR3A, S100B, SLC25A25B, EDIL3A, MACF1B, SMOC2, PVALB9, FAT4, PCDH1G31, KCNIP3A, UNC13BB, MACF1A, CRACR2AA | 880 | 739 | 17340 | 1,71 | 0,028242471 | 0,014324224 | 0,014288368 |
| GO:0016301 | kinase activity | Molecular Function | 60 | 5,319149 | 1,72E-04 | RET, SI:CH211-195B13.1, PFKFB2B, MOB1BA, PIK3CG, HK2, STK24B, RPS6KA3A, RPS6KA2, PIP5KL1, RPS6KA1, RPS6KB1B, AKT1, SI:CH211-243J20.2, PIM3, ERBB4B, PDGFRA, PRKCI, MAP4K3A, PRKCBB, DCLK1A, SI:DKEY-172J4.3, EPHA4A, ACVR1BB, MAPK14A, PI4K2A, GNE, IPPK, ITPK1B, UCK2A, LTK, PRKX, TTK, CITA, PIP4K2AA, PRKCZ, PFKPA, CKMT1, PIP5K1CA, GRK6, ERBB2, TRIOA, CAMK1GB, MAPK6, CHKB, YES1, TNK2B, LIMK2, PTK2AB, HIPK2, PRKACBB, ETNK2, JAK2B, RPS6KAL, PRKCHA, CMPK, PI4KB, PTK7B, CDK14, CAMK1DA | 880 | 718 | 17340 | 1,65 | 0,128144638 | 0,032432442 | 0,03235126 |
| GO:0035091 | phosphatidylinositol binding | Molecular Function | 16 | 1,41844 | 1,87E-04 | ESYT1A, ESYT2B, ESYT2A, ARHGAP32A, SI:CH211-195B13.1, ITPR1B, PITPNBL, SNX21, STAM, ZCCHC14, TOM1L2, SH3YL1, SNX10A, HIP1RB, SNX7 | 880 | 102 | 17340 | 3,09 | 0,138769321 | 0,032432442 | 0,03235126 |
| GO:0004711 | ribosomal protein S6 kinase activity | Molecular Function | 5 | 0,443262 | 2,03E-04 | RPS6KA3A, RPS6KA2, RPS6KA1, RPS6KAL, RPS6KB1B | 880 | 7 | 17340 | 14,07 | 0,149710731 | 0,032432442 | 0,03235126 |
| GO:0005388 | calcium-transporting ATPase activity | Molecular Function | 5 | 0,443262 | 0,003245591 | SI:DKEY-28B4.8, ATP2B1A, ATP2A2B, ATP2B3B, ATP2C1 | 880 | 13 | 17340 | 7,58 | 0,925536363 | 0,401975219 | 0,400969023 |
| GO:0004222 | metalloendopeptidase activity | Molecular Function | 15 | 1,329787 | 0,003521685 | MMP15B, MMEL1, PAPPAA, MMP17A, PHEX, PITRM1, ADAMTSL4, ADAMTSL3, ADAMTS17, MEP1A.1, MMP28, MMP19, ADAMTSL7, ADAMTS9, ADAM19B | 880 | 122 | 17340 | 2,42 | 0,940322064 | 0,401975219 | 0,400969023 |
| GO:0000166 | nucleotide binding | Molecular Function | 111 | 9,840426 | 0,004302395 | ADCY1A, PANK2, TAOK2A, NUBP2, RPS6KA3A, RPS6KA2, RPS6KA1, DHX58, AKT1, PIM3, EHD1B, PDGFRA, PRKCI, ENTPD1, UBE2E3, MAP4K3A, AARSD1, DCLK1A, SI:CH211-256M1.8, PRKAR1B, EPHA4A, ATP1A2A, ATP2B1A, MAPK14A, ROR2, PI4K2A, ABCG1, IPPK, UCK2A, ITPK1B, DDX5, RTEL1, DHX8, ARL6, RRAD, ABCB5, MYO6A, TUBA8L3, TUBA8L2, CITA, PIP4K2AA, PRKCZ, PFKPA, PIP5K1CA, GRK6, TRIOA, CAMK1GB, MYH10, RHOCB, SI:DKEY-32E23.4, ABCC6A, ABCA2, RAB4B, YES1, MYO15AA, EIF5, JAK2B, PRKCHA, PI4KB, CDK14, RALAA, KIF13BA, RET, SI:CH211-195B13.1, HSP90AB1, MCM7, HK2, KIF15, ARL5C, PIP5KL1, ATP2A2B, RPS6KB1B, SI:CH211-243J20.2, ERBB4B, ATP1A1A.4, MBD3B, GNL1, PRKCBB, SI:DKEY-28B4.8, UBE2R2, SI:DKEY-172J4.3, ACVR1BB, EEF1A1B, BLVRA, LTK, ATP10D, PRKX, TTK, ATP2C1, ADCY7, CKMT1, ATP1A3A, RASD1, SI:CH211-257P13.3, ERBB2, ARF3A, MAPK6, KIF26BA, MAP4K4, TNK2B, MYO1EA, PTK2AB, HIPK2, PRKACBB, ATP2B3B, RPS6KAL, ABCG2A, CMPK, KRAS, CAMK1DA, RAN | 880 | 1703 | 17340 | 1,28 | 0,968096199 | 0,429701734 | 0,428626135 |
| GO:0001517 | N-acetylglucosamine 6-O-sulfotransferase activity | Molecular Function | 4 | 0,35461 | 0,006000047 | CHST6, CHST7, CHST2B, SI:CH73-62B13.1 | 880 | 8 | 17340 | 9,85 | 0,991839725 | 0,510765439 | 0,509486928 |
| GO:0005085 | guanyl-nucleotide exchange factor activity | Molecular Function | 21 | 1,861702 | 0,006592485 | BCAR3, ARHGEF10, RIC1, DOCK4B, IQSEC3A, DENND4C, RAB3IP, DOCK7, TIAM1B, RAB3IL1, ARHGEF7A, ARHGEF9B, ARHGEF7B, RAPGEFL1, SI:DKEY-91M11.5, NET1, BCR, MADD, TRIOA, SBF1, RASGEF1BA | 880 | 215 | 17340 | 1,92 | 0,994932132 | 0,510765439 | 0,509486928 |
| GO:0051015 | actin filament binding | Molecular Function | 23 | 2,039007 | 0,007522058 | ACTR3, MBD3B, MYO1EA, FHOD1, ACTN1, MYO6A, MYO15AA, NEB, SHROOM1, ARPC5LA, CORO2BA, DUB, PSTPIP1A, SAMD14, HIP1RB, ARPC3, CFL2, CFL1, TWF1B, GAS2L1, CTNNAL1, MYH10, GSNA | 880 | 247 | 17340 | 1,83 | 0,997601334 | 0,510765439 | 0,509486928 |
| GO:0016409 | palmitoyltransferase activity | Molecular Function | 7 | 0,620567 | 0,00767107 | ZDHHC8B, ZDHHC9, ZDHHC13, ZDHHC23B, ZDHHC17, ZDHHC14, ZDHHC4 | 880 | 35 | 17340 | 3,94 | 0,997872515 | 0,510765439 | 0,509486928 |

**Supplementary Table 4** 76 Significantly enriched Gene Ontology terms with Fisher's exact test p-value < 0.01 in genes down-regulated in electric organ.

| Term | GO terms | Category | Count | % | P-value | Genes | List Total | Pop Hits | Pop Total | Fold Enrichment | Bonferroni | Benjamini | FDR |
|---|---|---|---|---|---|---|---|---|---|---|---|---|---|
| GO:0048741 | skeletal muscle fiber development | Biological Process | 22 | 2,263374 | 1,52E-13 | *CAVIN4B, KLHL41A, MYBPC1, PYGMA, RBFOX2, SMPX, MYO18AB, RBFOX1L, LGALS2A, MYO18AA, SIX1B, RYR3, ACTN2B, LMOD3, RYR1B, RYR1A, NFIXA, KLHL40B, KLHL40A, KLHL41B, SMYD1B, MYF5* | 820 | 63 | 18397 | 7,834572203 | 2,09E-10 | 2,09E-10 | 2,06E-10 |
| GO:0006936 | muscle contraction | Biological Process | 20 | 2,057613 | 3,14E-12 | *TMOD1, TNNI1C, MYHB, TNNI4A, TMOD4, TPM3, TPM1, TNNT3B, TNNT2A, MYOM2A, MYOM1A, MYOM1B, LMOD3, LMOD2B, SPEGB, TNNI2A.4, TNNT2E, TPMA, DESMA, TNNI2A.1* | 820 | 58 | 18397 | 7,736333053 | 4,33E-09 | 2,16E-09 | 2,13E-09 |
| GO:0030239 | myofibril assembly | Biological Process | 16 | 1,646091 | 6,69E-11 | *TMOD1, TMOD4, TNNT3B, ACTN2B, LMOD3, LMOD2B, TTN.2, TNNI2A.4, TTN.1, MEF2AA, MEF2AB, PROX1A, CRYABA, DESMA, PGM5, SMYD1B* | 820 | 40 | 18397 | 8,974146341 | 9,22E-08 | 3,07E-08 | 3,02E-08 |
| GO:0007519 | skeletal muscle tissue development | Biological Process | 18 | 1,851852 | 2,25E-09 | *MYOG, POPDC3, CDKN1A, DNAJB6A, MYLPFB, FHL1A, STAC3, FXR1, SYNPO2LA, SYNPO2LB, TTN.2, BAG3, TTN.1, NFIXA, CRYABA, DESMA, ITGA7, MYF5* | 820 | 65 | 18397 | 6,212870544 | 3,10E-06 | 7,75E-07 | 7,61E-07 |
| GO:0045214 | sarcomere organization | Biological Process | 15 | 1,54321 | 6,83E-09 | *KLHL41A, CAPN3A, LRRC39, TFPI2, TNNT3B, TNNT2A, ACTN2B, SMYHC2, TTN.2, TNNT2E, MYH7L, DESMA, FLNCB, KLHL41B, SMYD1B* | 820 | 46 | 18397 | 7,31588017 | 9,42E-06 | 1,88E-06 | 1,85E-06 |
| GO:0003009 | skeletal muscle contraction | Biological Process | 10 | 1,028807 | 4,70E-07 | *TNNI1C, RYR1B, TNNI4A, ZMP:0000000930, TNNI2A.4, TNNC1B, RYR1A, STAC3, TNNI2A.1, TCAP* | 820 | 24 | 18397 | 9,348069106 | 6,48E-04 | 1,08E-04 | 1,06E-04 |
| GO:0060048 | cardiac muscle contraction | Biological Process | 11 | 1,131687 | 1,05E-06 | *SMYHC2, TNNI1C, MYL13, TNNI4A, ZMP:0000000930, TNNI2A.4, TNNC1B, MYH7L, TNNI2A.1, TNNT2A, TCAP* | 820 | 33 | 18397 | 7,478455285 | 0,001451569 | 2,08E-04 | 2,04E-04 |
| GO:0006096 | glycolytic process | Biological Process | 11 | 1,131687 | 5,68E-06 | *PFKMB, GPIB, INSRA, PGAM2, PKMA, TPI1B, ALDOAA, ENO3, ALDOAB, GAPDH, ALDOCB* | 820 | 39 | 18397 | 6,327923702 | 0,007795445 | 9,78E-04 | 9,61E-04 |
| GO:0014866 | skeletal myofibril assembly | Biological Process | 7 | 0,720165 | 4,17E-05 | *MYO18AB, TPM3, MYO18AA, TMOD4, TTN.1, DUSP27, SMYD1B* | 820 | 16 | 18397 | 9,815472561 | 0,055863454 | 0,006387031 | 0,006271 |
| GO:0030036 | actin cytoskeleton organization | Biological Process | 19 | 1,954733 | 7,32E-05 | *PDLIM5B, PDLIM3B, PHACTR3B, ACTN3B, EHBP1L1A, EHBP1L1B, ACTN2B, SSH2A, ROCK2A, CAPZB, STARD13B, DAAM2, CAPZA1B, LDB3B, XIRP1, FLNA, CORO1CA, ZGC:162952, SMTNL1* | 820 | 144 | 18397 | 2,960221883 | 0,096053207 | 0,010098108 | 0,009915 |

| | | | | | | | | | | | | | |
|---|---|---|---|---|---|---|---|---|---|---|---|---|---|
| GO:0016310 | phosphorylation | Biological Process | 55 | 5,658436 | 1,18E-04 | *COQ8AA, MYLK2, DYRK4, CDKN1A, PRKAB1A, MAST2, CKMT2A, AKAP8L, CAMK2N1A, ROCK2A, EEF2K, ULK1B, MYLK4A, PLAUA, PIK3R3B, ADKB, GRK7A, PRKG1B, ADKA, ZGC:172076, HUNK, BMPR1AA, MAPKAPK3, PIK3CA, PRKCQ, MET, UCKL1B, VEGFAA, DAPK2A, RAF1A, AK1, CITA, PAK1, INSRA, ERBB2, ABL1, CDKN1CA, MAP2K6, SRPK3, PDK2A, CKMB, PFKMB, CKMA, NEK6, SI:CH211-220I18.4, NEK7, CAMK2B1, PKMA, ALPK3A, PTK2AA, AKT3A, SI:DKEY-8E10.3, MAPKAPK2A, PFKFB4B, SI:DKEY-96F10.1* | 820 | 718 | 18397 | 1,718586521 | 0,149904153 | 0,014763328 | 0,014496 |
| GO:0006099 | tricarboxylic acid cycle | Biological Process | 8 | 0,823045 | 2,80E-04 | *CS, FH, SUCLA2, MDH2, IDH2, DLST, ACO2, IDH3A* | 820 | 30 | 18397 | 5,982764228 | 0,320253974 | 0,032165168 | 0,031582 |
| GO:0048769 | sarcomerogenesis | Biological Process | 5 | 0,514403 | 4,11E-04 | *ZMP:0000000930, TTN.2, TTN.1, TCAP, SMYD1B* | 820 | 9 | 18397 | 12,46409214 | 0,432339201 | 0,040436785 | 0,039704 |
| GO:0035914 | skeletal muscle cell differentiation | Biological Process | 5 | 0,514403 | 4,11E-04 | *MYOG, CDKN1A, FHL1A, KLHL41B, MYF5* | 820 | 9 | 18397 | 12,46409214 | 0,432339201 | 0,040436785 | 0,039704 |
| GO:0055001 | muscle cell development | Biological Process | 9 | 0,925926 | 5,41E-04 | *FXR1, CAPZB, MYH7BA, NRAP, TCAP, ACTN3B, NEB, TGFBI, ACTN2B* | 820 | 43 | 18397 | 4,695774248 | 0,525580303 | 0,049697423 | 0,048796 |
| GO:0045727 | positive regulation of translation | Biological Process | 6 | 0,617284 | 6,83E-04 | *FXR1, FXR2, PCIF1, METTL5, LARP4B, LARP1B* | 820 | 17 | 18397 | 7,918364419 | 0,61028456 | 0,058876034 | 0,057809 |
| GO:0006874 | cellular calcium ion homeostasis | Biological Process | 10 | 1,028807 | 0,001264 | *ATP2A2A, RYR1B, RYR1A, ATP2B3B, HOMER1B, TNNT2A, ATP2A1, ATP2B2, RYR3, DHRS7CB* | 820 | 60 | 18397 | 3,739227642 | 0,825237001 | 0,102542457 | 0,100683 |
| GO:0061061 | muscle structure development | Biological Process | 6 | 0,617284 | 0,001532 | *PDLIM5B, PDLIM3B, LDB3B, HOMER1B, KLHL40B, KLHL40A* | 820 | 20 | 18397 | 6,730609756 | 0,879206058 | 0,115330509 | 0,11324 |
| GO:0033693 | neurofilament bundle assembly | Biological Process | 4 | 0,411523 | 0,001589 | *SYNM, SI:DKEY-33C12.3, NEFMA, NEFLA* | 820 | 6 | 18397 | 14,95691057 | 0,888421012 | 0,115330509 | 0,11324 |
| GO:0006470 | protein dephosphorylation | Biological Process | 16 | 1,646091 | 0,001923 | *EPM2A, SI:CH211-223P8.8, PTPN4A, PTPN21, DUSP27, DUSP16, CDC25B, SSH2A, PDP1, PTPRNA, DUSP10, SI:CH211-121A2.2, DUSP22A, SI:CH211-195B15.8, DUSP13A, DUSP22B* | 820 | 144 | 18397 | 2,492818428 | 0,929674738 | 0,132603536 | 0,1302 |
| GO:0016311 | dephosphorylation | Biological Process | 15 | 1,54321 | 0,002433 | *EPM2A, SI:CH211-223P8.8, PTPN4A, PTPN21, DUSP27, DUSP16, PTP4A3A, SSH2A, PTPRNA, DUSP10, SI:CH211-121A2.2, DUSP22A, SI:CH211-195B15.8, DUSP13A, DUSP22B* | 820 | 133 | 18397 | 2,53030442 | 0,965225892 | 0,153840334 | 0,151051 |
| GO:0016567 | protein ubiquitination | Biological Process | 32 | 3,292181 | 0,002548 | *VCP, ANAPC16, KLHL15, ASB5B, PDZRN3B, UBR3, NEDD4L, KLHL13, ASB18, SH3RF1, ASB16, TRIM35-31, CAND2, HERC2, ASB15B, ASB10, DCAF12, CUL3B, SI:CH73-54F23.4, SOCS3B, TRIM55B, FEM1A, KLHL21, ZBTB16A, FBXO32, UBAC1, FBXO31, SI:CH211-120G10.1, ASB2A.1, NEURL2, ASB4, TRIM54* | 820 | 406 | 18397 | 1,768304698 | 0,970353175 | 0,153840334 | 0,151051 |
| GO:0007623 | circadian rhythm | Biological Process | 7 | 0,720165 | 0,002566 | *NFIL3-6, NR0B2A, PER1B, CLOCKA, MITFA, NPAS2, ARNTL2* | 820 | 32 | 18397 | 4,90773628 | 0,971069964 | 0,153840334 | 0,151051 |
| GO:0060047 | heart contraction | Biological Process | 11 | 1,131687 | 0,002841 | *BAG3, TTN.2, LRRC39, CRYABA, DESMA, DLST, TNNT2A, TCAP, FBXO32, SMYD1B, LIMS1* | 820 | 80 | 18397 | 3,084862805 | 0,980211305 | 0,161969 | 0,159033 |

| | | | | | | | | | | | | | |
|---|---|---|---|---|---|---|---|---|---|---|---|---|---|
| GO:0017148 | negative regulation of translation | Biological Process | 7 | 0,720165 | 0,00302 | *FXR1, PAIP2B, FXR2, EIF4EBP1, CAPRIN1A, EIF4EBP3L, YBX1* | 820 | 33 | 18397 | 4,759016999 | 0,984558527 | 0,161969 | 0,159033 |
| GO:0007015 | actin filament organization | Biological Process | 16 | 1,646091 | 0,003054 | *TMOD1, MYO5AA, TMOD4, TPM3, TPM1, RHOBTB4, MYO16, LMOD3, TMSB, LMOD2B, TPMA, XIRP1, CORO6, BCL2L16, RHOAC, CORO1CA* | 820 | 151 | 18397 | 2,377257309 | 0,985266585 | 0,161969 | 0,159033 |
| GO:0032922 | circadian regulation of gene expression | Biological Process | 8 | 0,823045 | 0,003497 | *NFIL3-6, BHLHE40, KDM8, CRY2, PER1B, NR1D1, CLOCKA, NPAS2* | 820 | 45 | 18397 | 3,988509485 | 0,992015785 | 0,178433114 | 0,175198 |
| GO:0043409 | negative regulation of MAPK cascade | Biological Process | 6 | 0,617284 | 0,003623 | *DUSP10, SI:CH211-223P8.8, SI:CH211-121A2.2, SI:CH211-195B15.8, DUSP13A, DUSP16* | 820 | 24 | 18397 | 5,608841463 | 0,993296997 | 0,178433114 | 0,175198 |
| GO:0046314 | phosphocreatine biosynthetic process | Biological Process | 4 | 0,411523 | 0,00416 | *CKMB, CKMA, CKMT2A, ZGC:172076* | 820 | 8 | 18397 | 11,21768293 | 0,99681468 | 0,185072082 | 0,181717 |
| GO:0045947 | negative regulation of translational initiation | Biological Process | 4 | 0,411523 | 0,00416 | *PAIP2B, EIF4EBP1, EIF4EBP3L, YBX1* | 820 | 8 | 18397 | 11,21768293 | 0,99681468 | 0,185072082 | 0,181717 |
| GO:2001243 | negative regulation of intrinsic apoptotic signaling pathway | Biological Process | 4 | 0,411523 | 0,00416 | *MCL1A, MCL1B, BCL2L16, BCL2L1* | 820 | 8 | 18397 | 11,21768293 | 0,99681468 | 0,185072082 | 0,181717 |
| GO:0030388 | fructose 1,6-bisphosphate metabolic process | Biological Process | 5 | 0,514403 | 0,004621 | *PFKMB, ALDOAA, ALDOAB, FBP2, ALDOCB* | 820 | 16 | 18397 | 7,011051829 | 0,998315766 | 0,193081178 | 0,189581 |
| GO:0006937 | regulation of muscle contraction | Biological Process | 5 | 0,514403 | 0,004621 | *TNNT2E, TNNC1B, TNNT3B, TNNT2A, ATP2A1* | 820 | 16 | 18397 | 7,011051829 | 0,998315766 | 0,193081178 | 0,189581 |
| GO:0070588 | calcium ion transmembrane transport | Biological Process | 12 | 1,234568 | 0,005678 | *ATP2A2A, RYR1B, CACNA1SA, RYR1A, CACNA2D2B, ATP2B3B, TRPM4A, ITPR3, ATP2A1, ATP2B2, CACNG1B, RYR3* | 820 | 102 | 18397 | 2,639454806 | 0,999611057 | 0,230287196 | 0,226112 |
| GO:0051694 | pointed-end actin filament capping | Biological Process | 4 | 0,411523 | 0,006035 | *TMOD1, LMOD2B, TMOD4, LMOD3* | 820 | 9 | 18397 | 9,971273713 | 0,999763133 | 0,231188731 | 0,226997 |
| GO:0055008 | cardiac muscle tissue morphogenesis | Biological Process | 4 | 0,411523 | 0,006035 | *ZMP:0000000930, LRRC39, TCAP, FBXO32* | 820 | 9 | 18397 | 9,971273713 | 0,999763133 | 0,231188731 | 0,226997 |
| GO:0055013 | cardiac muscle cell development | Biological Process | 5 | 0,514403 | 0,007238 | *RBFOX2, ZMP:0000000930, RBFOX1L, TCAP, NR2F2* | 820 | 18 | 18397 | 6,23204607 | 0,999955381 | 0,269758239 | 0,264868 |
| GO:0030240 | skeletal muscle thin filament assembly | Biological Process | 4 | 0,411523 | 0,008339 | *ZMP:0000000930, TCAP, LMOD3, SMYD1B* | 820 | 10 | 18397 | 8,974146341 | 0,999990343 | 0,294860619 | 0,289515 |
| GO:0006108 | malate metabolic process | Biological Process | 4 | 0,411523 | 0,008339 | *FH, MDH2, ME1, ME3* | 820 | 10 | 18397 | 8,974146341 | 0,999990343 | 0,294860619 | 0,289515 |
| GO:0030018 | Z disc | Cellular Component | 26 | 2,674897 | 4,63E-17 | *FHL1A, ACTN3B, RYR3, SYNPO2LA, SYNPO2LB, ZMP:0000000930, BAG3, MYOZ2A, MYOZ2B, NRAP, DESMA, CASQ1B, PDLIM5B, PDLIM3B, NEB, PARVB, ACTN2B, RYR1B, MYOZ3A, RYR1A, LDB3B, MYOZ1A, MYOZ1B, TCAP, TRIM54, LIMS1* | 827 | 69 | 18868 | 8,596954244 | 1,43E-14 | 1,43E-14 | 1,37E-14 |
| GO:0016529 | sarcoplasmic reticulum | Cellular Component | 15 | 1,54321 | 6,45E-12 | *KLHL41A, CASQ1B, ITPR3, JPH1A, ATP2A1, JPH1B, RYR3, TRDN, ATP2A2A, RYR1B, JPH2, RYR1A, TMEM38A, KLHL41B, THBS4B* | 827 | 30 | 18868 | 11,40749698 | 1,99E-09 | 6,78E-10 | 6,52E-10 |

| | | | | | | | | | | | | | |
|---|---|---|---|---|---|---|---|---|---|---|---|---|---|
| GO:0005737 | cytoplasm | Cellular Component | 278 | 28,60082 | 6,61E-12 | *APOBEC2B, UGP2B, LRRC14B, PRKAB1A, AGLA, CALCOCO1A, DCAF6, ZFYVE28, NR0B2A, ROCK2A, HERC2, EIF2D, DUSP13A, CHAC1, KLHL41B, SMU1A, JMJD4, CAVIN4B, KLHL41A, RFX2, ARMC8, RNF123, RUFY3, ULK2, PRKCQ, TTLL12, HPRT1, SI:CH211-195B15.8, KLHL40B, KLHL40A, CCNO, UCKL1B, ILRUN, NUMBL, ANAPC16, CAPN3A, DAPK2A, SGIP1A, PTPN4A, NEDD4L, NMD3, APBB2B, TRIM35-31, FXR1, LDHA, FXR2, PCBP4, CMYA5, MYL10, NAA50, SRPK3, MYO5AA, CAMK2B1, PARVB, UBAC1, ALS2B, DAZL, MAPKAPK2A, MYH7L, DUSP22A, ABLIM1A, DUSP22B, FARSB, SYNM, BTG2, DNAJB6A, SVILA, SETD3, MYLPFB, ACY1, LRRC39, RPLP0, PDZRN3B, STON2, SH3RF1, SMG6, SSH2A, MSI2B, GYS1, UCHL1, KIF1B, ZGC:85777, SI:CH73-54F23.4, MYL12.2, PLEKHO1B, CNOT6L, RBFOX1L, MYO18AB, ARG1, MYO18AA, PTGR2, TMSB, EIF4EBP3L, CLOCKA, CACTIN, ACSBG2, KANK2, FH, SIX1B, VCLB, ARNTL2, PAK1, TTN.2, FRZB, DESMA, MAP3K3, SI:CH211-260E23.9, MDH2, SI:CH211-220I18.4, STAC3, BBOX1, PKMA, SMCR8A, TUBB4B, SMYHC2, RNF146, SI:CH211-120G10.1* [illegible] | 827 | 4405 | 18868 | 1,439856599 | 2,03E-09 | 6,78E-10 | 6,52E-10 |
| GO:0031430 | M band | Cellular Component | 8 | 0,823045 | 1,96E-07 | *SMPX, SPEGB, LRRC39, MYOM2A, MYOM1A, MYOM1B, LMOD3, SMYD1B* | 827 | 12 | 18868 | 15,20999597 | 6,05E-05 | 1,37E-05 | 1,32E-05 |
| GO:0033017 | sarcoplasmic reticulum membrane | Cellular Component | 9 | 0,925926 | 2,23E-07 | *ATP2A2A, RYR1B, KLHL41A, RYR1A, ATP2A1, TMEM38A, KLHL41B, RYR3, TRDN* | 827 | 17 | 18868 | 12,07852621 | 6,87E-05 | 1,37E-05 | 1,32E-05 |
| GO:0016460 | myosin II complex | Cellular Component | 10 | 1,028807 | 1,43E-05 | *SMYHC2, MYL12.2, MYHB, MYO18AB, MYL13, MYO18AA, MYH7BA, MYH7L, MYH14, MYLZ3* | 827 | 35 | 18868 | 6,518569701 | 0,00439237 | 7,34E-04 | 7,05E-04 |
| GO:0005829 | cytosol | Cellular Component | 77 | 7,921811 | 2,97E-05 | *IPO11, UBE3A, MTR, IPO7, DUSP16, LARP1B, ZFYVE28, PSME4B, PSME4A, MAP1LC3A, MID1IP1L, ARHGDIA, EEF2L2, PGM5, CHAC1, ACY3.1, ZGC:136908, PGM1, SI:DKEY-51E6.1, USP9, ADSL, ADKB, ARG1, PDE4D, ADKA, ACOT12, AMPD1, RBP7B, LARP4B, ALDOAA, ALDOAB, IRS2B, HPRT1, SI:CH211-195B15.8, GAPDH, ASPA, FBP2, ZGC:64002, USP13, VCP, FH, AHCY, ANAPC16, RAF1A, AK1, STRIP2, BAG3, G3BP1, CDAB, AAMP, ZC3H15, NAA50, USP24, GPD1B, RIC1, GPIB, SI:CH211-260E23.9, OSBPL5, PLEKHA5, STAC3, RAD23AA, MTHFR, USP28, ALDOCB, MLLT11, RNF146, ACOT11A, AGBL1, CASTOR2, PPP2R2BB, PFKFB4B, DUSP22A, SI:DKEY-96F10.1, ACO2, TPI1B, LRCH3, DUSP22B* | 827 | 1082 | 18868 | 1,623617869 | 0,009106642 | 0,001306889 | 0,001256 |

| | | | | | | | | | | | | | |
|---|---|---|---|---|---|---|---|---|---|---|---|---|---|
| GO:0005861 | troponin complex | Cellular Component | 8 | 0,823045 | 9,52E-05 | *TNNI1C, TNNI4A, TNNI2A.4, TNNT2E, TNNC1B, TNNI2A.1, TNNT3B, TNNT2A* | 827 | 26 | 18868 | 7,01999814 | 0,028883725 | 0,003663459 | 0,003521 |
| GO:0042383 | sarcolemma | Cellular Component | 10 | 1,028807 | 1,72E-04 | *POPDC3, RYR1B, SGCB, KCNB1, RYR1A, STAC3, DESMA, PGM5, VCLB, RYR3* | 827 | 47 | 18868 | 4,854254033 | 0,051663583 | 0,005893489 | 0,005664 |
| GO:0015629 | actin cytoskeleton | Cellular Component | 16 | 1,646091 | 1,98E-04 | *SVILA, MYO5AA, ARHGAP32B, VCLB, PARVB, MYO16, SYNPO2LA, SYNPO2LB, MYOZ3A, MYOZ2A, MYOZ2B, MYOZ1A, TPMA, MYOZ1B, ABLIM1A, CORO1CA* | 827 | 118 | 18868 | 3,093558502 | 0,059160439 | 0,006097662 | 0,00586 |
| GO:0032982 | myosin filament | Cellular Component | 7 | 0,720165 | 3,68E-04 | *SMYHC2, MYHB, MYO18AB, MYO18AA, MYH7BA, MYH7L, MYH14* | 827 | 23 | 18868 | 6,943693812 | 0,107093176 | 0,010295656 | 0,009895 |
| GO:0030016 | myofibril | Cellular Component | 6 | 0,617284 | 0,001106 | *TMOD1, LMOD2B, TMOD4, TNNT2A, TWF2B, LMOD3* | 827 | 19 | 18868 | 7,204734933 | 0,288790797 | 0,028383349 | 0,027278 |
| GO:0031941 | filamentous actin | Cellular Component | 6 | 0,617284 | 0,002246 | *PDLIM5B, PDLIM3B, LDB3B, EHBP1L1A, EHBP1L1B, SMTNL1* | 827 | 22 | 18868 | 6,222271078 | 0,499670009 | 0,053208425 | 0,051135 |
| GO:0016459 | myosin complex | Cellular Component | 10 | 1,028807 | 0,00248 | *SMYHC2, SMYHC3, MYO5AA, MYHB, MYO18AB, MYO18AA, MYH7BA, MYH7L, MYH14, MYO16* | 827 | 67 | 18868 | 3,405222978 | 0,534560246 | 0,054558852 | 0,052433 |
| GO:0031463 | Cul3-RING ubiquitin ligase complex | Cellular Component | 6 | 0,617284 | 0,006748 | *KLHL15, KLHL21, KLHL13, KLHL40B, KLHL40A, CUL3B* | 827 | 28 | 18868 | 4,888927276 | 0,875745965 | 0,132589038 | 0,127423 |
| GO:0005856 | cytoskeleton | Cellular Component | 43 | 4,423868 | 0,006888 | *DYRK4, FHOD1, PTPN4A, TUBA8L2, PTPN21, RHOBTB4, HSPB1, LRMP, KRT18A.1, VCLB, CITA, LMOD3, SSH2A, ACTB2, ROCK2A, MID1IP1L, CAPZB, SGCB, DCAF12, KLHL41B, RHOAC, MAP2K6, TMOD1, KLHL41A, ABI2A, TMOD4, CCDC135, TPM1, KLHL21, PARVB, TUBB4B, PTK2AA, TMSB, EPB41L3B, EPB41L3A, DNMBP, LMOD2B, TPMA, TWF2B, TACC2, GAPDH, KANK4, CORO1CA* | 827 | 645 | 18868 | 1,520999597 | 0,881015762 | 0,132589038 | 0,127423 |
| GO:0005911 | cell-cell junction | Cellular Component | 9 | 0,925926 | 0,00947 | *IGSF11, MPP1, EPB41L3B, USP53B, EPB41L3A, DNMBP, MPP7A, DESMA, LIMS1* | 827 | 68 | 18868 | 3,019631553 | 0,946641091 | 0,17157736 | 0,164893 |
| GO:0003779 | actin binding | Molecular Function | 50 | 5,144033 | 2,02E-13 | *ABRAB, MYHB, SVILA, ABRAA, SETD3, SSH2A, SYNPO2LA, SYNPO2LB, CAPZB, MYO18AB, TPM3, MYO18AA, PDLIM3B, TPM1, ACTN2B, TMSB, EPB41L3B, EPB41L3A, DAAM2, MYOZ3A, TPMA, CORO1CA, MICAL2A, TNNT2A, ACTN3B, VCLB, SMTNL, MYOZ2A, NRAP, MYOZ2B, XIRP1, FLNCB, MYH14, FLNA, MYO5AA, PDLIM5B, PHACTR3B, NEB, PARVB, MYO16, SMYHC2, SMYHC3, MYH7BA, LDB3B, MYH7L, CAPZA1B, MYOZ1A, MYOZ1B, ABLIM1A, TWF2B* | 771 | 337 | 17340 | 3,336835664 | 1,41E-10 | 1,41E-10 | 1,38E-10 |
| GO:0051015 | actin filament binding | Molecular Function | 36 | 3,703704 | 1,19E-09 | *MYHB, SVILA, FHOD1, ACTN3B, MYOM1A, VCLB, MYOM1B, CAPZB, SPEGB, NRAP, XIRP1, MYH14, FLNA, FLNCB, TMOD1, MYO5AA, MYO18AB, TPM3, MYO18AA, TMOD4, TNNC1B, TPM1, MYOM2A, NEB, MYO16, ACTN2B, SMYHC2, SMYHC3, MYH7BA, CAPZA1B, MYH7L, TPMA, CORO6, ABLIM1A, TWF2B, CORO1CA* | 771 | 247 | 17340 | 3,277934435 | 8,30E-07 | 4,15E-07 | 4,08E-07 |

| | | | | | | | | | | | | | |
|---|---|---|---|---|---|---|---|---|---|---|---|---|---|
| GO:0008138 | protein tyrosine/serine/threonine phosphatase activity | Molecular Function | 12 | 1,234568 | 3,50E-05 | *EPM2A, DUSP10, SI:CH211-223P8.8, SI:CH211-121A2.2, DUSP22A, SI:CH211-195B15.8, DUSP13A, DUSP27, DUSP16, DUSP22B, PTP4A3A, SSH2A* | 771 | 57 | 17340 | 4,734794184 | 0,024053012 | 0,008115528 | 0,007975 |
| GO:0005523 | tropomyosin binding | Molecular Function | 7 | 0,720165 | 6,11E-05 | *TMOD1, LMOD2B, TNNT2E, TMOD4, TNNT3B, TNNT2A, LMOD3* | 771 | 17 | 17340 | 9,260700389 | 0,04156231 | 0,010612357 | 0,010429 |
| GO:0004879 | RNA polymerase II transcription factor activity, ligand-activatec | Molecular Function | 13 | 1,337449 | 1,07E-04 | *ESR2A, RXRAA, RARAA, PPARDB, RORC, RXRGB, NR1D1, NR2F2, NR2F5, NR2F6B, NR2F1A, RORAA, ESRRGA* | 771 | 75 | 17340 | 3,898313878 | 0,07192424 | 0,012041058 | 0,011833 |
| GO:0031433 | telethonin binding | Molecular Function | 5 | 0,514403 | 1,21E-04 | *MYOZ3A, MYOZ2A, MYOZ2B, MYOZ1A, MYOZ1B* | 771 | 7 | 17340 | 16,06448027 | 0,080837652 | 0,012041058 | 0,011833 |
| GO:0051373 | FATZ binding | Molecular Function | 5 | 0,514403 | 1,21E-04 | *MYOZ3A, MYOZ2A, MYOZ2B, MYOZ1A, MYOZ1B* | 771 | 7 | 17340 | 16,06448027 | 0,080837652 | 0,012041058 | 0,011833 |
| GO:0003824 | catalytic activity | Molecular Function | 35 | 3,600823 | 1,67E-04 | *APOBEC2B, FH, TKTB, AGLA, CKMT2A, GLULB, ACACB, HADHAA, LDHA, HYAL3, ENPP6, CDAB, PFKMB, CKMB, ADSL, PYGMA, CKMA, MDH2, MOCOS, PGAM2, ZGC:172076, PKMA, ALDOAA, ALDOAB, ALDOCB, GOT2B, SUCLA2, IMPDH1A, PCYT1BA, GOT2A, PFKFB4B, SI:DKEY-96F10.1, DUS1L, TPI1B, BCAT1* | 771 | 393 | 17340 | 2,002950466 | 0,109314057 | 0,014469219 | 0,014219 |
| GO:0016301 | kinase activity | Molecular Function | 54 | 5,555556 | 2,00E-04 | *COQ8AA, MYLK2, DYRK4, CDKN1A, PRKAB1A, MAST2, CKMT2A, AKAP8L, CAMK2N1A, ROCK2A, EEF2K, ULK1B, MYLK4A, PLAUA, PIK3R3B, ADKB, GRK7A, PRKG1B, ADKA, ZGC:172076, HUNK, BMPR1AA, MAPKAPK3, PIK3CA, PRKCQ, MET, UCKL1B, DAPK2A, RAF1A, AK1, CITA, PAK1, INSRA, ERBB2, ABL1, CDKN1CA, MAP2K6, SRPK3, PDK2A, CKMB, PFKMB, CKMA, NEK6, SI:CH211-220I18.4, NEK7, CAMK2B1, PKMA, ALPK3A, PTK2AA, AKT3A, SI:DKEY-8E10.3, MAPKAPK2A, PFKFB4B, SI:DKEY-96F10.1* | 771 | 718 | 17340 | 1,691468953 | 0,129839425 | 0,015451511 | 0,015185 |
| GO:0016791 | phosphatase activity | Molecular Function | 20 | 2,057613 | 2,54E-04 | *EPM2A, SI:CH211-223P8.8, PTPN4A, PTPN21, PPTC7A, DUSP27, DUSP16, PTP4A3A, SSH2A, PDP1, PTPRNA, CTDSPL2A, DUSP10, SI:CH211-121A2.2, DUSP22A, SI:CH211-195B15.8, CTDSPLB, DUSP13A, FBP2, DUSP22B* | 771 | 173 | 17340 | 2,600031488 | 0,161930356 | 0,017663163 | 0,017358 |
| GO:0004721 | phosphoprotein phosphatase activity | Molecular Function | 17 | 1,748971 | 4,52E-04 | *EPM2A, SI:CH211-223P8.8, PTPN4A, PTPN21, PPTC7A, DUSP16, CDC25B, SSH2A, PDP1, CTDSPL2A, DUSP10, SI:CH211-121A2.2, DUSP22A, SI:CH211-195B15.8, CTDSPLB, DUSP13A, DUSP22B* | 771 | 139 | 17340 | 2,750608851 | 0,269638051 | 0,026531485 | 0,026073 |

| | | | | | | | | | | | | | |
|---|---|---|---|---|---|---|---|---|---|---|---|---|---|
| GO:0003700 | transcription factor activity, sequence-specific DNA binding | Molecular Function | 46 | 4,73251 | 4,58E-04 | *ESR2A, RXRAA, RARAA, CREB5B, HOXA9B, SIX1B, RORC, MAFAA, RXRGB, NPAS2, ARNTL2, NR2F6B, PBX3A, NFIXA, ZNF395A, FOXO3B, FOXO3A, CREMA, HLFA, PITX3, HOXC4A, NFIL3-6, JUND, ATF5A, PPARDB, TBX3A, RFX2, MAFBA, NFATC1, FOXN3, NR1D1, NR2F2, MITFA, NR2F5, POU3F1, MYCB, TBX15, NFATC3A, RFX6, MYCH, NFIC, CLOCKA, NR2F1A, RORAA, ESRRGA, ATF3* | 771 | 602 | 17340 | 1,718525796 | 0,272725119 | 0,026531485 | 0,026073 |
| GO:0016740 | transferase activity | Molecular Function | 105 | 10,80247 | 0,001036 | *COQ8AA, UGP2B, MYLK2, TKTB, UBE2D4, PRKAB1A, AGLA, CKMT2A, CAMK2N1A, UBE3A, ROCK2A, ULK1B, HHATLA, PIK3R3B, KCMF1, UBE2E1, BMPR1AA, MAPKAPK3, PRKCQ, HPRT1, ZGC:64002, UCKL1B, LIPT2, DAPK2A, AK1, UBR3, CITA, MOGAT3A, INSRA, ABL1, COX10, MAP2K6, NAA50, SRPK3, NEK6, GFPT2, NEK7, PCIF1, UBE2G1A, CAMK2B1, ALPK3A, CS, SI:DKEY-8E10.3, MAPKAPK2A, PFKFB4B, SI:DKEY-96F10.1, DYRK4, CDKN1A, SETD3, MAST2, PDZRN3B, AKAP8L, SETD7, MTR, CDC34A, SH3RF1, SMG6, DTWD2, GYS1, EEF2K, MYLK4A, NTMT1, PLAUA, DLAT, ZGC:162952, GRK7A, ADKB, PRKG1B, ADKA, ZGC:172076, HUNK, TRPT1, ASH1L, SIRT3, GOT2B, METTL22, PIK3CA, GOT2A, SI:CH211-269C21.2, BCAT1, GAPDH, MET, RAF1A, DLST, TGM2B, RNF2, PAK1, ERBB2, METTL5, GPAT3, CDKN1CA, SMYD1B, PDK2A, CKMB, PYGMA, PFKMB, CKMA, SI:CH211-220I18.4, MOCOS, PKMA, PTK2AA, AKT3A, RNF146, GATM, PCMT* | 771 | 1743 | 17340 | 1,354835685 | 0,513478159 | 0,055392321 | 0,054436 |
| GO:0004725 | protein tyrosine phosphatase activity | Molecular Function | 15 | 1,54321 | 0,001778 | *EPM2A, SI:CH211-223P8.8, PTPN4A, PTPN21, DUSP16, PTP4A3A, CDC25B, SSH2A, PTPRNA, DUSP10, SI:CH211-121A2.2, DUSP22A, SI:CH211-195B15.8, DUSP13A, DUSP22B* | 771 | 129 | 17340 | 2,61514795 | 0,709717396 | 0,088271452 | 0,086747 |

| | | | | | | | | | | | | | |
|---|---|---|---|---|---|---|---|---|---|---|---|---|---|
| GO:0000978 | RNA polymerase II core promoter proximal region sequence-sp | Molecular Function | 70 | 7,201646 | 0,002018 | *HIVEP2A, CREB5B, HOXA9B, ZBTB47B, RORC, MAFAA, RXRGB, PBX3A, ZNF648, FOXO3B, FOXO3A, CREMA, HLFA, PITX3, MYOG, TBX3A, PAX4, RFX2, SOX13, TFEB, ZBTB16A, MAFBA, ZBTB34, MITFA, POU3F1, KLF15, SOX10, EGR2B, MYCB, RFX6, MYCH, MEF2AA, KLF11B, MEF2AB, CLOCKA, DACHC, RORAA, ESRRGA, DACHD, ATF3, ESR2A, RXRAA, RARAA, SIX1B, NPAS2, ARNTL2, NR2F6B, NFIXA, HOXA10B, IRX3A, ZNF395A, HOXC4A, ZBTB18, MLXIP, JUND, SIX4B, PPARDB, NFATC1, NR1D1, NR2F2, NR2F5, KLF4, TBX15, NFATC3A, TFCP2, NFIC, BHLHE40, PROX1A, NR2F1A, MYF5* | 771 | 1095 | 17340 | 1,437734307 | 0,754294653 | 0,093480387 | 0,091866 |
| GO:0004111 | creatine kinase activity | Molecular Function | 4 | 0,411523 | 0,00413 | *CKMB, CKMA, CKMT2A, ZGC:172076* | 771 | 8 | 17340 | 11,24513619 | 0,943659776 | 0,179400167 | 0,176303 |
| GO:0051019 | mitogen-activated protein kinase binding | Molecular Function | 5 | 0,514403 | 0,005778 | *DUSP10, MAPKAPK3, MAPKAPK2A, SI:CH211-195B15.8, DUSP16* | 771 | 17 | 17340 | 6,614785992 | 0,982184439 | 0,236237342 | 0,232158 |

**Supplementary Table 5** 19 Significantly enriched Gene Ontology terms with Fisher's exact test p-value < 0.05 among genes with increasing expression relative to EOD duration (Group 5 and 6).

| Term | GO terms | Category | Count | % | P-value | Genes | List Total | Pop Hits | Pop Total | Fold Enrichment | Bonferroni | Benjamini | FDR |
|---|---|---|---|---|---|---|---|---|---|---|---|---|---|
| GO:0005975 | carbohydrate metabolic process | Biological Process | 11 | 4,564315 | 7,83E-05 | *CHST7, GNPDA2, MAN2B2, B3GAT3, GLB1, RPE, SI:DKEY-199F5.8, SPATA20, GUSB, YDJC, HK2* | 205 | 199 | 18397 | 4,960583405 | 0,044687604 | 0,045715087 | 0,045715087 |
| GO:0048675 | axon extension | Biological Process | 4 | 1,659751 | 0,010509161 | *IST1, UBAP1, PLXNB1B, GRNB* | 205 | 41 | 18397 | 8,755264723 | 0,997908403 | 1 | 1 |
| GO:0006914 | autophagy | Biological Process | 5 | 2,074689 | 0,01365593 | *BECN1, GABARAPL2, PLEKHM1, WIPI1, MCOLN1A* | 205 | 83 | 18397 | 5,406112254 | 0,999674458 | 1 | 1 |
| GO:0007032 | endosome organization | Biological Process | 3 | 1,244813 | 0,022373383 | *PLEKHF2, IST1, PI4K2A* | 205 | 21 | 18397 | 12,82020906 | 0,999998176 | 1 | 1 |
| GO:0046854 | phosphatidylinositol phosphorylation | Biological Process | 4 | 1,659751 | 0,032820251 | *PIP5K1CA, IMPA1, PIK3CG, PI4K2A* | 205 | 63 | 18397 | 5,697870693 | 0,999999997 | 1 | 1 |
| GO:0006811 | ion transport | Biological Process | 13 | 5,394191 | 0,041416232 | *SLC24A2, SLC12A4, SLC10A7, SLC31A2, KCTD12.1, KCNJ2A, SFXN5B, ATP6V0A1A, TCN2, ATP6V1D, PANX3, MCU, ATP6V1F* | 205 | 614 | 18397 | 1,900063558 | 1 | 1 | 1 |
| GO:0006665 | sphingolipid metabolic process | Biological Process | 3 | 1,244813 | 0,043438725 | *ARV1, PSAP, SFTPBB* | 205 | 30 | 18397 | 8,974146341 | 1 | 1 | 1 |
| GO:0005764 | lysosome | Cellular Component | 11 | 4,564315 | 3,03E-05 | *ASAH1B, MAN2B2, BRI3, CTSBA, SMPD1, VPS41, PSAP, PLEKHM1, MCOLN1A, SFTPBB, GUSB* | 201 | 186 | 18868 | 5,551489863 | 0,005199004 | 0,005212487 | 0,005182182 |
| GO:0005794 | Golgi apparatus | Cellular Component | 18 | 7,46888 | 3,93E-04 | *ZDHHC16B, BECN1, SLC10A7, YIPF3, B3GAT3, CHPF2, ENTPD6, SYAP1, SCOCA, FUT9A, ARV1, MGAT1B, TRAPPC6B, VPS41, SI:DKEY-199F5.8, EXTL3, ZGC:162698, PI4K2A* | 201 | 628 | 18868 | 2,690559939 | 0,065338118 | 0,033778584 | 0,033582197 |
| GO:0005768 | endosome | Cellular Component | 11 | 4,564315 | 6,34E-04 | *PLEKHF2, BECN1, RAB5B, RAB40C, VPS41, PLEKHM1, UBAP1, FLOT1B, COMMD1, LAMTOR3, PI4K2A* | 201 | 270 | 18868 | 3,824359683 | 0,103297005 | 0,03633201 | 0,036120778 |
| GO:0005829 | cytosol | Cellular Component | 21 | 8,713693 | 0,011107253 | *BECN1, GABARAPL2, USP7, RIC1, IRS1, RPE, GSR, DNAJB1A, CST14A.2, PPM1AA, WIPI1, SCOCA, PRDX6, HK2, RSPH1, EIF5, UBAP1, OSBPL1A, RAN, ZGC:162698, SPG21* | 201 | 1082 | 18868 | 1,821888708 | 0,853559593 | 0,477611894 | 0,47483508 |
| GO:0005773 | vacuole | Cellular Component | 3 | 1,244813 | 0,018737755 | *TMEM138, GLB1, LGMN* | 201 | 20 | 18868 | 14,08059701 | 0,961360155 | 0,595562503 | 0,592099931 |
| GO:0031410 | cytoplasmic vesicle | Cellular Component | 8 | 3,319502 | 0,020775436 | *RAC3A, BECN1, VMA21, TBC1D7, VPS41, FLOT1B, RHOCB, PI4K2A* | 201 | 259 | 18868 | 2,899479437 | 0,972975637 | 0,595562503 | 0,592099931 |
| GO:0005581 | collagen trimer | Cellular Component | 4 | 1,659751 | 0,024709798 | *COL8A1A, COL12A1A, COL2A1B, COL4A5* | 201 | 59 | 18868 | 6,364111645 | 0,986478523 | 0,607155038 | 0,603625067 |

| | | | | | | | | | | | | | |
|---|---|---|---|---|---|---|---|---|---|---|---|---|---|
| GO:0005783 | endoplasmic reticulum | Cellular Component | 15 | 6,224066 | 0,03295827 | *ZDHHC16B, PLEKHF2, BECN1, SLC10A7, TAPBPL, HSPBP1, AGPAT2, CYP51, ARV1, ZFYVE27, SPTLC1, TRAPPC6B, VMA21, FKBP9, EXTL3* | 201 | 763 | 18868 | 1,845425559 | 0,996862548 | 0,708602798 | 0,704483014 |
| GO:0004185 | serine-type carboxypeptidase activity | Molecular Function | 3 | 1,244813 | 6,00E-04 | *CTSA, SCPEP1, SI:CH211-122F10.4* | 176 | 4 | 17340 | 73,89204545 | 0,166179401 | 0,181682518 | 0,181682518 |
| GO:0004180 | carboxypeptidase activity | Molecular Function | 3 | 1,244813 | 0,02229738 | *CTSA, SCPEP1, SI:CH211-122F10.4* | 176 | 23 | 17340 | 12,85079051 | 0,998921912 | 1 | 1 |
| GO:0046961 | proton-transporting ATPase activity, rotational mechanism | Molecular Function | 3 | 1,244813 | 0,026095058 | *ATP6V0A1A, ATP6V1D, ATP6V1F* | 176 | 25 | 17340 | 11,82272727 | 0,999668472 | 1 | 1 |
| GO:0016798 | hydrolase activity, acting on glycosyl bonds | Molecular Function | 4 | 1,659751 | 0,047207987 | *MAN2B2, GLB1, SMPD1, GUSB* | 176 | 80 | 17340 | 4,926136364 | 0,999999567 | 1 | 1 |

**Supplementary Table 6** 41 Significantly enriched Gene Ontology terms with Fisher's exact test p-value < 0.05 among genes with decreasing expression relative to EOD duration (Group 3).

| Term | GO terms | Category | Count | % | P-value | Genes | List Total | Pop Hits | Pop Total | Fold Enrichment | Bonferroni | Benjamini | FDR |
|---|---|---|---|---|---|---|---|---|---|---|---|---|---|
| GO:0007411 | axon guidance | Biological Process | 12 | 3,71517 | 8,12E-05 | *SEMA3AB, CNTN1A, SEMA6A, SEMA4D, LAMA1, NPTNB, EPHA4A, SEMA4C, EFNB2A, NRCAMA, TRIOA, DPYSL2B* | 271 | 182 | 18397 | 4,47597421 | 0,055436912 | 0,057030482 | 0,056949 |
| GO:0030198 | extracellular matrix organization | Biological Process | 9 | 2,786378 | 7,83E-04 | *ADAMTS5, COL5A3A, CCDC80, ADAMTSL4, COL1A1A, COL2A1B, ADAMTS17, ADAMTS9, ADAMTS15A* | 271 | 133 | 18397 | 4,593763005 | 0,423029942 | 0,274874767 | 0,274483 |
| GO:0006486 | protein glycosylation | Biological Process | 9 | 2,786378 | 0,001198 | *B3GAT1A, GALNT7, NUS1, GALNT14, AARS2, ST8SIA5, FUT8A, LARGE2, ST3GAL2* | 271 | 142 | 18397 | 4,302609012 | 0,568895602 | 0,280300317 | 0,279901 |
| GO:0031103 | axon regeneration | Biological Process | 4 | 1,23839 | 0,001825 | *B3GAT1A, SEMA4D, LINGO1A, DPYSL2B* | 271 | 17 | 18397 | 15,97308444 | 0,72266857 | 0,294500541 | 0,294081 |
| GO:0007155 | cell adhesion | Biological Process | 16 | 4,95356 | 0,002098 | *DCHS1B, PCDH15B, EGFL6, LAMA1, PCDH19, CCN2B, CNTN1A, VCANB, EPHA4A, ITGA10, NPTNB, EFNB2A, NCAM1A, ITGB8, FBN2B, BCAR1* | 271 | 437 | 18397 | 2,485514283 | 0,77100365 | 0,294500541 | 0,294081 |
| GO:0016310 | phosphorylation | Biological Process | 21 | 6,501548 | 0,004751 | *UCK2A, CHKB, MARK3B, DGKB, PFKFB2B, EIF2AK3, DGKZA, NUCKS1A, CDK7, CDK5, EPHB4A, ETNK2, EPHA4A, GRK6, CKBB, SI:CH211-243J20.2, MAP3K8, MAPK14A, TRIOA, PIK3R3B, CAMK1DA* | 271 | 718 | 18397 | 1,985512237 | 0,964684651 | 0,555914712 | 0,555123 |
| GO:0008045 | motor neuron axon guidance | Biological Process | 5 | 1,547988 | 0,008075 | *SEMA3AB, LAMA1, BICD1A, NOTUM2, NTN2* | 271 | 54 | 18397 | 6,285704524 | 0,996626008 | 0,809806795 | 0,808653 |
| GO:0048843 | negative regulation of axon extension involved in axon guidance | Biological Process | 4 | 1,23839 | 0,014473 | *SEMA3AB, SEMA6A, SEMA4D, SEMA4C* | 271 | 35 | 18397 | 7,758355298 | 0,999964072 | 1 | 1 |
| GO:0007420 | brain development | Biological Process | 9 | 2,786378 | 0,024678 | *ADGRL2A, SCGN, CNTN1A, SEMA4D, LAMA1, TAOK2A, NRCAMA, PCDH19, DPYSL2B* | 271 | 238 | 18397 | 2,567102856 | 0,999999976 | 1 | 1 |
| GO:0050919 | negative chemotaxis | Biological Process | 4 | 1,23839 | 0,025058 | *SEMA3AB, SEMA6A, SEMA4D, SEMA4C* | 271 | 43 | 18397 | 6,314940359 | 0,999999982 | 1 | 1 |
| GO:0006468 | protein phosphorylation | Biological Process | 19 | 5,882353 | 0,025768 | *ADAM10B, MARK3B, TAOK2A, EIF2AK3, GUCY2F, CDK7, BRSK2B, CDK5, EPHB4A, EPHA4A, GRK6, MAP3K8, MAPK14A, TRIOA, FAM20CB, NRBP2B, CAMK1DA, SI:CH211-1I11.3, MARK1* | 271 | 741 | 18397 | 1,740656637 | 0,999999989 | 1 | 1 |
| GO:0035475 | angioblast cell migration involved in selective angioblast sprouting | Biological Process | 2 | 0,619195 | 0,029138 | *EPHB4A, EFNB2A* | 271 | 2 | 18397 | 67,88560886 | 0,999999999 | 1 | 1 |
| GO:0033334 | fin morphogenesis | Biological Process | 3 | 0,928793 | 0,031137 | *FRAS1, COL1A1A, COL2A1B* | 271 | 19 | 18397 | 10,71878035 | 1 | 1 | 1 |
| GO:0090630 | activation of GTPase activity | Biological Process | 5 | 1,547988 | 0,033837 | *ARHGAP22, SI:CH211-288D18.1, SI:DKEY-191M6.4, SIPA1L1, AGAP3* | 271 | 83 | 18397 | 4,089494509 | 1 | 1 | 1 |
| GO:0048013 | ephrin receptor signaling pathway | Biological Process | 3 | 0,928793 | 0,037515 | *EPHB4A, EFNB2A, ANKS1AB* | 271 | 21 | 18397 | 9,697944122 | 1 | 1 | 1 |
| GO:0071526 | semaphorin-plexin signaling pathway | Biological Process | 4 | 1,23839 | 0,038847 | *SEMA3AB, SEMA6A, SEMA4D, SEMA4C* | 271 | 51 | 18397 | 5,324361479 | 1 | 1 | 1 |
| GO:0003404 | optic vesicle morphogenesis | Biological Process | 2 | 0,619195 | 0,043388 | *EPHA4A, EFNB2A* | 271 | 3 | 18397 | 45,25707257 | 1 | 1 | 1 |
| GO:0001756 | somitogenesis | Biological Process | 5 | 1,547988 | 0,046537 | *CCDC80, MEF2AA, EFNB2A, MAPK14A, DLC* | 271 | 92 | 18397 | 3,689435264 | 1 | 1 | 1 |
| GO:0030903 | notochord development | Biological Process | 4 | 1,23839 | 0,046918 | *COL5A3A, EGFL6, LAMA1, COL2A1B* | 271 | 55 | 18397 | 4,93713519 | 1 | 1 | 1 |
| GO:0031012 | extracellular matrix | Cellular Component | 13 | 4,024768 | 1,29E-04 | *COLEC12, COL5A3A, CCN2B, ADAMTS5, VCANB, ADAMTSL4, COL1A1A, ADAMTS17, COL2A1B, FBN2B, ADAMTS9, LINGO1A, ADAMTS15A* | 287 | 218 | 18868 | 3,920404053 | 0,023360005 | 0,023635648 | 0,023636 |
| GO:0005794 | Golgi apparatus | Cellular Component | 21 | 6,501548 | 0,001505 | *GALNT7, ARHGAP32A, ADAM10B, GALNT14, AARS2, CLASP1A, SACM1LB, FUT8A, ZDHHC23B, B4GALNT3A, ZDHHC14, B3GAT1A, ST8SIA5, ZGC:162200, FAM20CB, BICD1A, LARGE2, GRINAA, ERGIC1, ST3GAL2, PLA2G4AB* | 287 | 628 | 18868 | 2,198384341 | 0,240907021 | 0,137711768 | 0,137712 |
| GO:0016020 | membrane | Cellular Component | 129 | 39,93808 | 0,008157 | *ADGRL2A, GALNT14, IFITM1, DGKB, RHOT1B, OLFCS1, XYLT1, ACSL4A, TMEM263, FADS2, SI:CH211-1E14.1, TNFSF11, EHD1B, SI:DKEY-174M14.3, LARGE2, INPP4AB, SYPL2B, ADAM10B, ENTPD2A.1, SEMA6A, NUP210, ATP6AP1A, GPSM2L, FUT8A, TRIM101, SLC5A9, ADAM15, ADGRB2, EPHA4A, MADD, TMCC1B, FAM20CB, GRAMD1BB, OSBPL3B, DIPK1B, WAIF2, AARS2, RPN2, SDC3, MTMR9, SACM1LB, PCDH19, RNF145B, HSD11B2, TMEM248, VPS11, ST8SIA5, GSG1L, SCN3B, ST3GAL2, LINGO1A, NUS1, TMEM132E, ANKRD22, TRPV1, SLC25A25B, FLRT1B, EPHB4A, NPTNB, LRFN1, ITGA10, SPIRE2, TSPAN5A, NCAM1A, SDK1A, LRCH4, TMEM19, MCTP1A, MACF1A, PIGS, KCNK6, TRPC6A, NRCAMA, VSIG10, B4GALNT3A, SPRED2A, SI:CH1073-291C23.2, DGKZA, CATIP, SI:DKEY-112M2.1, CPT2, EFNB2A, ITGB8, GAL3ST4, TMEM119B, DCHS1B, PCDH15B, EGFL6, ANO6, MMEL1, SLC39A14, ZDHHC14, RRBP1A, B3GAT1A, KCNK4A, RAP2B, ECRG4A, TMEM218, BIRC6, DLC, NAT8L, GRINAA, TNFRSF21, ERGIC1, COLEC12, GRAMD1C, LAMA1, PIP4P1A, GUCY2F, ADCY7, PPP1R3AA, IGSF21A, CNTN1A, SI:CH211-278J3.3, GDPD4A, KCNQ5A, KCNQ5B, GALNT7, SLC35A1, SEMA4C, ZDHHC23B, MGAT4C, SPCS2, JAGN1B, FRAS1, GDPD5B, EPPK1, SI:CH73-267C23.10, GPR146* | 287 | 7112 | 18868 | 1,192454819 | 0,776603689 | 0,487709159 | 0,487709 |
| GO:0005576 | extracellular region | Cellular Component | 27 | 8,359133 | 0,01066 | *LAMA1, IGFBP6B, CXCL19, OLFM3A, FGF2, HTRA1B, NTN2, CXCL18A.1, SEMA3AB, ADAMTS5, GLIPR2, ADAMTSL4, COL1A1A, ADAMTS17, ENPP2, FBN2B, NOTUM2, ADAMTS9, MARK1, ADAMTS15A, COL5A3A, CCN2B, VCANB, CCDC80, DKK3A, ECRG4A, COL2A1B* | 287 | 1059 | 18868 | 1,676145729 | 0,859325897 | 0,487709159 | 0,487709 |
| GO:0043231 | intracellular membrane-bounded organelle | Cellular Component | 11 | 3,405573 | 0,024173 | *EPN3A, HSD11B2, SPCS2, PHLPP1, HIP1RB, PDXDC1, DHRS9, EHD1B, NAT8L, RHOBTB1, OSBPL3B* | 287 | 319 | 18868 | 2,266971044 | 0,98864384 | 0,884730212 | 0,88473 |
| GO:0000139 | Golgi membrane | Cellular Component | 11 | 3,405573 | 0,03016 | *B3GAT1A, GALNT7, SLC35A1, GALNT14, AARS2, XYLT1, SACM1LB, ZDHHC23B, LARGE2, ERGIC1, ZDHHC14* | 287 | 331 | 18868 | 2,184784783 | 0,996317753 | 0,886341935 | 0,886342 |
| GO:0015629 | actin cytoskeleton | Cellular Component | 6 | 1,857585 | 0,033904 | *MTSS1LA, ARHGAP32A, CATIP, HAX1, MYO5B, MACF1A* | 287 | 118 | 18868 | 3,342821709 | 0,998185616 | 0,886341935 | 0,886342 |
| GO:0005930 | axoneme | Cellular Component | 4 | 1,23839 | 0,042102 | *HYDIN, CFAP36, CFAP206, DNALI1* | 287 | 51 | 18868 | 5,156247865 | 0,999618585 | 0,91141975 | 0,91142 |

| | | | | | | | | | | | | | |
|---|---|---|---|---|---|---|---|---|---|---|---|---|---|
| GO:0005938 | cell cortex | Cellular Component | 5 | 1,547988 | 0,044824 | *ARHGAP32A, SPIRE2, GPSM2L, RHOBTB1, MACF1A* | 287 | 88 | 18868 | 3,735350016 | 0,999773395 | 0,91141975 | 0,91142 |
| GO:0005509 | calcium ion binding | Molecular Function | 24 | 7,430341 | 4,28E-04 | *DCHS1B, SNED1, PCDH15B, DIPK1B, CALM1A, EGFL6, DGKB, RHOT1B, PDCD6, ACTN1, PCDH19, SLC25A25B, EDIL3A, SCGN, VCANB, PPP3R1B, EFHD2, ENPP2, DLC, FBN2B, EHD1B, MCTP1A, MACF1A, PLA2G4AB* | 250 | 739 | 17340 | 2,25255751 | 0,139265272 | 0,149936794 | 0,149937 |
| GO:0004222 | metalloendopeptidase activity | Molecular Function | 8 | 2,47678 | 0,001912 | *ADAMTS5, ADAM10B, ADAM15, ADAMTSL4, ADAMTS17, MMEL1, ADAMTS9, ADAMTS15A* | 250 | 122 | 17340 | 4,548196721 | 0,488191854 | 0,323477718 | 0,323478 |
| GO:0016740 | transferase activity | Molecular Function | 40 | 12,3839 | 0,003459 | *UBE2NB, UGP2B, UCK2A, GALNT14, AARS2, DGKB, PFKFB2B, XYLT1, B4GALNT3A, DGKZA, SI:CH211-278J3.3, CPT2, GRK6, ST8SIA5, SI:CH211-243J20.2, MAP3K8, TRIOA, LARGE2, PIK3R3B, ST3GAL2, GALNT7, NUS1, CHKB, MARK3B, EIF2AK3, FUT8A, ZDHHC23B, ZDHHC14, MGAT4C, B3GAT1A, NUCKS1A, CDK7, CDK5, ETNK2, EPHB4A, EPHA4A, CKBB, MAPK14A, NAT8L, CAMK1DA* | 250 | 1743 | 17340 | 1,591738382 | 0,702598533 | 0,323477718 | 0,323478 |
| GO:0016301 | kinase activity | Molecular Function | 21 | 6,501548 | 0,003697 | *UCK2A, CHKB, MARK3B, DGKB, PFKFB2B, EIF2AK3, DGKZA, NUCKS1A, CDK7, CDK5, EPHB4A, ETNK2, EPHA4A, GRK6, CKBB, SI:CH211-243J20.2, MAP3K8, MAPK14A, TRIOA, PIK3R3B, CAMK1DA* | 250 | 718 | 17340 | 2,028635097 | 0,726460298 | 0,323477718 | 0,323478 |
| GO:0030215 | semaphorin receptor binding | Molecular Function | 4 | 1,23839 | 0,01261 | *SEMA3AB, SEMA6A, SEMA4D, SEMA4C* | 250 | 34 | 17340 | 8,16 | 0,988223357 | 0,778839411 | 0,778839 |
| GO:0045499 | chemorepellent activity | Molecular Function | 4 | 1,23839 | 0,014735 | *SEMA3AB, SEMA6A, SEMA4D, SEMA4C* | 250 | 36 | 17340 | 7,706666667 | 0,994458678 | 0,778839411 | 0,778839 |
| GO:0016757 | transferase activity, transferring glycosyl groups | Molecular Function | 10 | 3,095975 | 0,019346 | *B3GAT1A, GALNT7, MGAT4C, GALNT14, AARS2, ST8SIA5, XYLT1, FUT8A, LARGE2, ST3GAL2* | 250 | 278 | 17340 | 2,494964029 | 0,998927326 | 0,778839411 | 0,778839 |
| GO:0005178 | integrin binding | Molecular Function | 5 | 1,547988 | 0,01995 | *CCN2B, EGFL6, ADAM15, ITGA10, ITGB8* | 250 | 72 | 17340 | 4,816666667 | 0,999135338 | 0,778839411 | 0,778839 |
| GO:0050321 | tau-protein kinase activity | Molecular Function | 3 | 0,928793 | 0,021586 | *BRSK2B, MARK3B, MARK1* | 250 | 16 | 17340 | 13,005 | 0,999518189 | 0,778839411 | 0,778839 |
| GO:0005524 | ATP binding | Molecular Function | 35 | 10,83591 | 0,022253 | *UBE2NB, UCK2A, AARS2, DGKB, PFKFB2B, TAOK2A, GUCY2F, SMC4, ADCY7, DGKZA, SMCHD1, SI:CH211-257P13.3, GRK6, DHX15, SI:CH211-243J20.2, MAP3K8, TRIOA, EHD1B, NRBP2B, SI:CH211-1I11.3, MARK1, ACTR3, MARK3B, ENTPD2A.1, STARD9, EIF2AK3, CDK7, BRSK2B, CDK5, EPHB4A, EPHA4A, MYO5B, CKBB, MAPK14A, CAMK1DA* | 250 | 1662 | 17340 | 1,460649819 | 0,999620387 | 0,778839411 | 0,778839 |
| GO:0016758 | transferase activity, transferring hexosyl groups | Molecular Function | 5 | 1,547988 | 0,032802 | *GALNT7, GALNT14, AARS2, LARGE2, B4GALNT3A* | 250 | 84 | 17340 | 4,128571429 | 0,99999148 | 1 | 1 |
| GO:0005085 | guanyl-nucleotide exchange factor activity | Molecular Function | 8 | 2,47678 | 0,03611 | *PREX2, BCAR3, RABGEF1, CCDC88C, MADD, ARHGEF9B, TRIOA, RAPGEFL1* | 250 | 215 | 17340 | 2,580837209 | 0,999997432 | 1 | 1 |
| GO:0005201 | extracellular matrix structural constituent | Molecular Function | 5 | 1,547988 | 0,044995 | *COL5A3A, LAMA1, COL1A1A, COL2A1B, FBN2B* | 250 | 93 | 17340 | 3,729032258 | 0,9999999 | 1 | 1 |

**Supplementary Table 7** Imbalanced expressed alleles and their *C. compressirostris* allele proportion for all five replicates of any hybrid cohort.

| Hybrid cohort | Tissue | SNPs ID | Gene ID in Annotation | Proportion 1 | Proportion 2 | Proportion 3 | Proportion 4 | Proportion 5 | Average Proportion | Gene | Hightlights of Predicted Function | Gene Description |
|---|---|---|---|---|---|---|---|---|---|---|---|---|
| *com x rhy* | EO | 1665681 | maker-ptg000267l-snap-gene-2.67-mRNA-1 | 0,64 | 0,63 | 0,64 | 0,7 | 0,67 | 0,66 | *ANKRD12* | ankyrin repeats-containing cofactor | ankyrin repeat domain 12 |
| *com x rhy* | EO | 681658 | maker-ptg000082l-snap-gene-20.7-mRNA-1 | 0,64 | 0,61 | 0,74 | 0,68 | 0,73 | 0,68 | *ARL13b* | cilium-specific protein | ADP ribosylation factor like GTPase 13B |
| *com x rhy* | EO | 2982031 | maker-ptg000598l-snap-gene-2.10-mRNA-1 | 0,65 | 0,79 | 0,81 | 0,68 | 0,67 | 0,72 | *CDH15* | calcium-dependent cell adhesion protein | cadherin 15 |
| *com x rhy* | EO | 1098949 | maker-ptg000160l-snap-gene-11.44-mRNA-1 | 0,86 | 0,82 | 0,79 | 0,81 | 0,79 | 0,82 | *CHRND* | opening of an ion-conducting channel across the plasma membrane. | cholinergic receptor nicotinic delta subunit |
| *com x rhy* | EO | 4997264 | maker-ptg001536l-augustus-gene-4.34-mRNA-1 | 0,61 | 0,61 | 0,79 | 0,6 | 0,64 | 0,65 | *COL6a3* | cell-binding protein | collagen type VI alpha 3 chain |
| *com x rhy* | EO | 4997265 | maker-ptg001536l-augustus-gene-4.34-mRNA-1 | 0,61 | 0,62 | 0,8 | 0,6 | 0,64 | 0,65 | *COL6a3* | cell-binding protein | collagen type VI alpha 3 chain |
| *com x rhy* | EO | 4531902 | maker-ptg001236l-augustus-gene-26.64-mRNA-1 | 0,71 | 0,75 | 0,79 | 0,74 | 0,66 | 0,73 | *CRELD1* | epidermal growth factor-related proteins | cysteine rich with EGF like domains 1 |
| *com x rhy* | EO | 4531904 | maker-ptg001236l-augustus-gene-26.64-mRNA-1 | 0,81 | 0,6 | 0,65 | 0,89 | 0,7 | | | | |
| *com x rhy* | EO | 1396998 | maker-ptg000216l-augustus-gene-4.4-mRNA-1 | 0,67 | 0,6 | 0,64 | 0,61 | 0,6 | 0,62 | *DAG1* | laminin and basement membrane assembly | dystroglycan 1 |
| *com x rhy* | EO | 4392819 | snap_masked-ptg001165l-processed-gene-0.145-mRNA-1 | 0,65 | 0,74 | 0,67 | 0,77 | 0,74 | 0,71 | *DCAF6* | ligand-dependent coactivator of nuclear receptors | DDB1 and CUL4 associated factor 6 |
| *com x rhy* | EO | 837824 | maker-ptg000110l-snap-gene-11.31-mRNA-1 | 0,67 | 0,61 | 0,71 | 0,61 | 0,61 | 0,64 | *DST* | cytoskeletal linker protein | dystonin |
| *com x rhy* | EO | 3404159 | maker-ptg000729l-snap-gene-5.79-mRNA-1 | 0,76 | 0,8 | 0,7 | 0,69 | 0,84 | 0,74 | *ENPP2* | phosphodiesterase | ectonucleotide pyrophosphatase/phosphodiesterase 2 |
| *com x rhy* | EO | 3404158 | maker-ptg000729l-snap-gene-5.79-mRNA-1 | 0,73 | 0,67 | 0,68 | 0,7 | 0,84 | | | | |
| *com x rhy* | EO | 3813376 | maker-ptg000876l-snap-gene-1.7-mRNA-1 | 0,66 | 0,67 | 0,71 | 0,63 | 0,6 | 0,65 | *HOXC11a* | multicellular organism development and regulation of transcription | homeobox C11a |
| *com x rhy* | EO | 1651431 | maker-ptg000265l-est_gff_est2genome-gene-6.33-mRNA-1 | 0,2 | 0,18 | 0,13 | 0,14 | 0,12 | 0,16 | *KCNJ2* | allow potassium to flow into a cell rather than out of a cell, probably participates in establishing action potential waveform | inward rectifier potassium channel 2 |
| *com x rhy* | EO | 768530 | snap_masked-ptg000100l-processed-gene-4.104-mRNA-1 | 0,81 | 0,69 | 0,77 | 0,88 | 0,77 | 0,71 | *OBSCN* | structural component of striated muscles which plays a role in myofibrillogenesis | obscurin, cytoskeletal calmodulin and titin-interacting RhoGEF |
| *com x rhy* | EO | 768526 | snap_masked-ptg000100l-processed-gene-4.104-mRNA-1 | 0,62 | 0,62 | 0,6 | 0,67 | 0,68 | | | | |
| *com x rhy* | EO | 1624861 | maker-ptg000254l-augustus-gene-4.109-mRNA-1 | 0,69 | 0,9 | 0,68 | 0,9 | 0,73 | 0,78 | *PALMD* | regulation of cell shape | palmdelphin |
| *com x rhy* | EO | 2766503 | maker-ptg000548l-snap-gene-0.90-mRNA-1 | 0,62 | 0,78 | 0,63 | 0,67 | 0,76 | 0,69 | *PNKD* | regulation of myofibrillogenesis | paroxysmal nonkinesigenic dyskinesia |
| *com x rhy* | EO | 4446575 | maker-ptg001188l-snap-gene-6.4-mRNA-1 | 0,66 | 0,63 | 0,61 | 0,69 | 0,69 | 0,66 | *SCN4aa* | voltage-gated sodium channel | sodium channel protein type 4 subunit alpha A |
| *com x rhy* | EO | 4446574 | maker-ptg001188l-snap-gene-6.4-mRNA-1 | 0,65 | 0,68 | 0,67 | 0,66 | 0,63 | | | | |
| *com x tsh* | EO | 2670824 | maker-ptg000509l-augustus-gene-3.41-mRNA-1 | 0,91 | 0,85 | 0,78 | 0,92 | 0,86 | 0,86 | *TSPAN7b* | integral component of plasma membrane | tetraspanin 7b |
| *com x tsh* | EO | 5058039 | maker-ptg001586l-snap-gene-1.60-mRNA-1 | 0,94 | 0,71 | 0,82 | 0,9 | 0,9 | 0,84 | *UACA* | modulates isoactin dynamics to regulate the morphological alterations required for cell growth and motility | uveal autoantigen with coiled-coil domains and ankyrin repeats |
| *com x tsh* | EO | 5058040 | maker-ptg001586l-snap-gene-1.60-mRNA-1 | 0,91 | 0,68 | 0,77 | 0,9 | 0,87 | | | | |
| *com x rhy* | SM | 3177368 | maker-ptg000959l-snap-gene-7.19-mRNA-1 | 0,72 | 0,6 | 0,67 | 0,67 | 0,68 | 0,67 | *PLPP3* | membrane glycoprotein at the cell plasma membrane | phospholipid phosphatase 3 |
| *com x rhy* | SM | 305826 | maker-ptg000051l-snap-gene-6.91-mRNA-1 | 0,73 | 0,67 | 0,61 | 0,68 | 0,64 | 0,67 | *HSP90aa1* | inducible molecular chaperone | heat shock protein 90 alpha family class a member 1 |
| *com x rhy* | SM | 3786101 | maker-ptg001394l-snap-gene-3.24-mRNA-1 | 0,66 | 0,7 | 0,65 | 0,8 | 0,6 | 0,68 | *MYO18a* | unconventional myosin | myosin XVIIIA |
| *com x rhy* | SM | 4016401 | maker-ptg001619l-snap-gene-1.135-mRNA-1 | 0,65 | 0,75 | 0,79 | 0,64 | 0,72 | 0,71 | *TRIM63* | muscle-specific rING finger protein | E3 ubiquitin-protein ligase TRIM63 |
| *com x rhy* | SM | 862477 | maker-ptg000160l-snap-gene-11.44-mRNA-1 | 0,8 | 0,8 | 0,7 | 0,69 | 0,83 | 0,76 | *CHRND* | opening of an ion-conducting channel across the plasma membrane | cholinergic receptor nicotinic delta subunit |
| *com x rhy* | SM | 606006 | snap_masked-ptg000100l-processed-gene-4.104-mRNA-1 | 0,84 | 0,7 | 0,83 | 0,84 | 0,75 | 0,79 | *OBSCN* | structural component of striated muscles which plays a role in myofibrillogenesis | obscurin, cytoskeletal calmodulin and titin-interacting RhoGEF |
| *com x rhy* | SM | 3426233 | maker-ptg001156l-augustus-gene-8.0-mRNA-1 | 0,91 | 0,9 | 0,92 | 0,95 | 0,92 | 0,92 | *CASQ1a* | skeletal muscle specific member of the calsequestrin protein | calsequestrin-1a |
| *com x tsh* | SM | 1814167 | maker-ptg000413l-snap-gene-5.117-mRNA-1 | 0,77 | 0,86 | 0,73 | 0,78 | 0,81 | 0,79 | *XIRP1* | protect actin filaments during depolymerization | xin actin-binding repeat-containing protein 1 |

**Supplementary Table 8** Sequence information of *KCNJ2* transcripts in all pure-bred individuals

| Species | Sequence |
|---|---|
| com1 | ATGAATGTCCAAAACTGTTCCTCAGAAGGTTCTTCCAAAAACTTTTCCAAGGCAGCAGAAGTGAAGCCCTCATCAGCAGAA<br>GCGATGGGTAGTGTGCGGGCCAGCCGCTACAGCGTCGTGTCCTCCAAAGTAGATGGCCTCAAGTTGGCCACTGTGGCCGT<br>GTCCAATGGCCACAGCAATGGTGATGGCAAGGTGAACATGTGGCAGCCGGTGCCATGTCGTTTCGTCAAGAAGGATGGA<br>CACTGCAACGTGCACATCATCAACATGAGCGAGAAAGGCCAGCGCTACATAGCCGACATCTTCACCACCTGCGTGGACATC<br>CGTTGGCGATGGATGATAATCATCTTCTGCTTGACTTTTGTGCTTTCATGGTTGTTCTTTGGCTATGTGTTCTGGCTGGTGG<br>CCTTCTTCTATGGTGACTTGGGGAATAGCTCCCAGCAGTGTGTCTCCAATGTCAACAGCTTCATGGCAGCCTTCCTCTTCTCT<br>GTGGAGACGCAGACCACTATTGGCTATGGTTACCACCATGTGACAGAAGAGTGCCCCATCGCTGTCTTTATGGTGGTTTTC<br>CAGTGCATTGTTGGCTGCATCATCGACGCCTTCATCATTGGTGCCGTCATGGCCAAGATGGCCAAGCCCACGAAGCGCAAT<br>GAAACCCTGGTGTTTAGCCACAACGCTACAATAGCAATGCGGGACGGCAAGCTATGCCTGATGTGGCGAGTTGGCAACCT<br>ACGCAAAAGCCACCTGGTGGAGGCCCACGTGAGGGCTCAGCTACTCAAGTCCCGGACCACCGCCGAGGGGGAGTTTATC<br>CCCCTAGACCACGTAGATATTGATGTGGGCTTTGACACTGGCGTAGACCGGATCTTCCTTGTTTCCCCCATCACCATTGTCC<br>ATGAGATCAACGAGGACAGTCCCTTCTATGATATGAGCAAGCAGGATTTTGAGACTGCTGGATTTGAGATTGTGGTCATCC<br>TGGAGGGCATGGTAGAAGCCACAGCCATGACAACCCAGTGTCGCAGTTCCTACCTGGCAGGGGAGATCCTCTGGGGACA<br>CTGCTTCGAGCCTGTACTCTTTGAGGAGAAGAACTACTACAAGGTCGACTACTCTCATTTCCACAAAACCTACGAGGTGCC<br>GAGCACTCCGCTATGTAGTGCGCGGGAGCTTGCTGAAAAGAAGGATAATGAGTCCAGCTCTAACTCTTTTTGCTATGAGA<br>ATGAAGTGGCGATGATGGACAAAGAGGAGACGGAGGACAAAAGCGAGTGCAGCAATGATGGGAGCAGTTCACAAAAGG<br>CTTCAGAGTTGGGGCGCAATCTCTTCATGACGTTTAGACGAGAATCTGAGATTTGA |

com2

ATGAATGTCCAAAACTGTTCCTCAGAAGGTTCTTCCAAAAACTTTTCCAAGGCAGCAGAAGTGAAGCCCTCATCAGCAGAA
GCGATGGGTAGTGTGCGGGCCAGCCGCTACAGCGTCGTGTCCTCCAAAGTAGATGGCCTCAAGTTGGCCACTGTGGCCGT
GTCCAATGGCCACAGCAATGGTGATGGCAAGGTGAACATGTGGCAGCCGGTGCCATGTCGTTTCGTCAAGAAGGATGGA
CACTGCAACGTGCACATCATCAACATGAGCGAGAAAGGCCAGCGCTACATAGCCGACATCTTCACCACCTGCGTGGACATC
CGTTGGCGATGGATGATAATCATCTTCTGCTTGACTTTTGTGCTTTCATGGTTGTTCTTTGGCTATGTGTTCTGGCTGGTGG
CCTTCTTCTATGGTGACTTGGGGAATAGCTCCCAGCAGTGTGTCTCCAATGTCAACAGCTTCATGGCAGCCTTCCTCTTCTCT
GTGGAGACGCAGACCACTATTGGCTATGGTTACCACCATGTGACAGAAGAGTGCCCCATCGCTGTCTTTATGGTGGTTTTC
CAGTGCATTGTTGGCTGCATCATCGACGCCTTCATCATTGGTGCCGTCATGGCCAAGATGGCCAAGCCCACGAAGCGCAAT
GAAACCCTGGTGTTTAGCCACAACGCTACAATAGCAATGCGGGACGGCAAGCTATGCCTGATGTGGCGAGTTGGCAACCT
ACGCAAAAGCCACCTGGTGGAGGCCCACGTGAGGGCTCAGCTACTCAAGTCCCGGACCACCGCCGAGGGGGAGTTTATC
CCCCTAGACCACGTAGATATTGATGTGGGCTTTGACACTGGCGTAGACCGGATCTTCCTTGTTTCCCCCATCACCATTGTCC
ATGAGATCAACGAGGACAGTCCCTTCTATGATATGAGCAAGCAGGATTTTGAGACTGCTGGATTTGAGATTGTGGTCATCC
TGGAGGGCATGGTAGAAGCCACAGCCATGACAACCCAGTGTCGCAGTTCCTACCTGGCAGGGGAGATCCTCTGGGGACA
CTGCTTCGAGCCTGTACTCTTTGAGGAGAAGAACTACTACAAGGTCGACTACTCTCATTTCCACAAAACCTACGAGGTGCC
GAGCACTCCGCTATGTAGTGCGCGGGAGCTTGCTGAAAAGAAGGATAATGAGTCCAGCTCTAACTCTTTTTGCTATGAGA
ATGAAGTGGCGATGATGGACAAAGAGGAGACGGAGGACAAAAGCGAGTGCAGCAATGATGGGAGCAGTTCACAAAAGG
CTTCAGAGTTGGGGCGCAATCTCTTCATGACGTTTAGACGAGAATCTGAGATTTGA

| | |
|---|---|
| com3 | ATGAATGTCCAAAACTGTTCCTCAGAAGGTTCTTCCAAAAACTTTTCCAAGGCAGCAGAAGTGAAGCCCTCATCAGCAGAA<br>GCGATGGGTAGTGTGCGGGCCAGCCGCTACAGCGTCGTGTCCTCCAAAGTAGATGGCCTCAAGTTGGCCACTGTGGCCGT<br>GTCCAATGGCCACAGCAATGGTGATGGCAAGGTGAACATGTGGCAGCCGGTGCCATGTCGTTTCGTCAAGAAGGATGGA<br>CACTGCAACGTGCACATCATCAACATGAGCGAGAAAGGCCAGCGCTACATAGCCGACATCTTCACCACCTGCGTGGACATC<br>CGTTGGCGATGGATGATAATCATCTTCTGCTTGACTTTTGTGCTTTCATGGTTGTTCTTTGGCTATGTGTTCTGGCTGGTGG<br>CCTTCTTCTATGGTGACTTGGGGAATAGCTCCCAGCAGTGTGTCTCCAATGTCAACAGCTTCATGGCAGCCTTCCTCTTCTCT<br>GTGGAGACGCAGACCACTATTGGCTATGGTTACCACCATGTGACAGAAGAGTGCCCCATCGCTGTCTTTATGGTGGTTTTC<br>CAGTGCATTGTTGGCTGCATCATCGACGCCTTCATCATTGGTGCCGTCATGGCCAAGATGGCCAAGCCCACGAAGCGCAAT<br>GAAACCCTGGTGTTTAGCCACAACGCTACAATAGCAATGCGGGACGGCAAGCTATGCCTGATGTGGCGAGTTGGCAACCT<br>ACGCAAAAGCCACCTGGTGGAGGCCCACGTGAGGGCTCAGCTACTCAAGTCCCGGACCACCGCCGAGGGGGAGTTTATC<br>CCCCTAGACCACGTAGATATTGATGTGGGCTTTGACACTGGCGTAGACCGGATCTTCCTTGTTTCCCCCATCACCATTGTCC<br>ATGAGATCAACGAGGACAGTCCCTTCTATGATATGAGCAAGCAGGATTTTGAGACTGCTGGATTTGAGATTGTGGTCATCC<br>TGGAGGGCATGGTAGAAGCCACAGCCATGACAACCCAGTGTCGCAGTTCCTACCTGGCAGGGGAGATCCTCTGGGGACA<br>CTGCTTCGAGCCTGTACTCTTTGAGGAGAAGAACTACTACAAGGTCGACTACTCTCATTTCCACAAAACCTACGAGGTGCC<br>GAGCACTCCGCTATGTAGTGCGCGGGAGCTTGCTGAAAAGAAGGATAATGAGTCCAGCTCTAACTCTTTTTGCTATGAGA<br>ATGAAGTGGCGATGATGGACAAAGAGGAGACGGAGGACAAAAGCGAGTGCAGCAATGATGGGAGCAGTTCACAAAAGG<br>CTTCAGAGTTGGGGCGCAATCTCTTCATGACGTTTAGACGAGAATCTGAGATTTGA |

com4 ATGAATGTCCAAAACTGTTCCTCAGAAGGTTCTTCCAAAAACTTTTCCAAGGCAGCAGAAGTGAAGCCCTCATCAGCAGAA
GCGATGGGTAGTGTGCGGGCCAGCCGCTACAGCGTCGTGTCCTCCAAAGTAGATGGCCTCAAGTTGGCCACTGTGGCCGT
GTCCAATGGCCACAGCAATGGTGATGGCAAGGTGAACATGTGGCAGCCGGTGCCATGTCGTTTCGTCAAGAAGGATGGA
CACTGCAACGTGCACATCATCAACATGAGCGAGAAAGGCCAGCGCTACATAGCCGACATCTTCACCACCTGCGTGGACATC
CGTTGGCGATGGATGATAATCATCTTCTGCTTGACTTTTGTGCTTTCATGGTTGTTCTTTGGCTATGTGTTCTGGCTGGTGG
CCTTCTTCTATGGTGACTTGGGGAATAGCTCCCAGCAGTGTGTCTCCAATGTCAACAGCTTCATGGCAGCCTTCCTCTTCTCT
GTGGAGACGCAGACCACTATTGGCTATGGTTACCACCATGTGACAGAAGAGTGCCCCATCGCTGTCTTTATGGTGGTTTTC
CAGTGCATTGTTGGCTGCATCATCGACGCCTTCATCATTGGTGCCGTCATGGCCAAGATGGCCAAGCCCACGAAGCGCAAT
GAAACCCTGGTGTTTAGCCACAACGCTACAATAGCAATGCGGGACGGCAAGCTATGCCTGATGTGGCGAGTTGGCAACCT
ACGCAAAAGCCACCTGGTGGAGGCCCACGTGAGGGCTCAGCTACTCAAGTCCCGGACCACCGCCGAGGGGGAGTTTATC
CCCCTAGACCACGTAGATATTGATGTGGGCTTTGACACTGGCGTAGACCGGATCTTCCTTGTTTCCCCCATCACCATTGTCC
ATGAGATCAACGAGGACAGTCCCTTCTATGATATGAGCAAGCAGGATTTTGAGACTGCTGGATTTGAGATTGTGGTCATCC
TGGAGGGCATGGTAGAAGCCACAGCCATGACAACCCAGTGTCGCAGTTCCTACCTGGCAGGGGAGATCCTCTGGGGACA
CTGCTTCGAGCCTGTACTCTTTGAGGAGAAGAACTACTACAAGGTCGACTACTCTCATTTCCACAAAACCTACGAGGTGCC
GAGCACTCCGCTATGTAGTGCGCGGGAGCTTGCTGAAAAGAAGGATAATGAGTCCAGCTCTAACTCTTTTTGCTATGAGA
ATGAAGTGGCGATGATGGACAAAGAGGAGACGGAGGACAAAAGCGAGTGCAGCAATGATGGGAGCAGTTCACAAAAGG
CTTCAGAGTTGGGGCGCAATCTCTTCATGACGTTTAGACGAGAATCTGAGATTTGA

com5 ATGAATGTCCAAAACTGTTCCTCAGAAGGTTCTTCCAAAAACTTTTCCAAGGCAGCAGAAGTGAAGCCCTCATCAGCAGAA
GCGATGGGTAGTGTGCGGGCCAGCCGCTACAGCGTCGTGTCCTCCAAAGTAGATGGCCTCAAGTTGGCCACTGTGGCCGT
GTCCAATGGCCACAGCAATGGTGATGGCAAGGTGAACATGTGGCAGCCGGTGCCATGTCGTTTCGTCAAGAAGGATGGA
CACTGCAACGTGCACATCATCAACATGAGCGAGAAAGGCCAGCGCTACATAGCCGACATCTTCACCACCTGCGTGGACATC
CGTTGGCGATGGATGATAATCATCTTCTGCTTGACTTTTGTGCTTTCATGGTTGTTCTTTGGCTATGTGTTCTGGCTGGTGG
CCTTCTTCTATGGTGACTTGGGGAATAGCTCCCAGCAGTGTGTCTCCAATGTCAACAGCTTCATGGCAGCCTTCCTCTTCTCT
GTGGAGACGCAGACCACTATTGGCTATGGTTACCACCATGTGACAGAAGAGTGCCCCATCGCTGTCTTTATGGTGGTTTTC
CAGTGCATTGTTGGCTGCATCATCGACGCCTTCATCATTGGTGCCGTCATGGCCAAGATGGCCAAGCCCACGAAGCGCAAT
GAAACCCTGGTGTTTAGCCACAACGCTACAATAGCAATGCGGGACGGCAAGCTATGCCTGATGTGGCGAGTTGGCAACCT
ACGCAAAAGCCACCTGGTGGAGGCCCACGTGAGGGCTCAGCTACTCAAGTCCCGGACCACCGCCGAGGGGGAGTTTATC
CCCCTAGACCACGTAGATATTGATGTGGGCTTTGACACTGGCGTAGACCGGATCTTCCTTGTTTCCCCCATCACCATTGTCC
ATGAGATCAACGAGGACAGTCCCTTCTATGATATGAGCAAGCAGGATTTTGAGACTGCTGGATTTGAGATTGTGGTCATCC
TGGAGGGCATGGTAGAAGCCACAGCCATGACAACCCAGTGTCGCAGTTCCTACCTGGCAGGGGAGATCCTCTGGGGACA
CTGCTTCGAGCCTGTACTCTTTGAGGAGAAGAACTACTACAAGGTCGACTACTCTCATTTCCACAAAACCTACGAGGTGCC
GAGCACTCCGCTATGTAGTGCGCGGGAGCTTGCTGAAAAGAAGGATAATGAGTCCAGCTCTAACTCTTTTTGCTATGAGA
ATGAAGTGGCGATGATGGACAAAGAGGAGACGGAGGACAAAAGCGAGTGCAGCAATGATGGGAGCAGTTCACAAAAGG
CTTCAGAGTTGGGGCGCAATCTCTTCATGACGTTTAGACGAGAATCTGAGATTTGA

tsh1

ATGAATGTCCAAAACTGTTCCTCAGAAGGTTCTTCCAAAAACTTTTCCAAGGCAGCAGAAGTGAAGCCCTCATCAGCAGAA
GCGATGGGTAGTGTGCGGGCCAGCCGCTACAGCGTCGTGTCCTCCAAAGTAGATGGCCTCAAGTTGGCCACTGTGGCCGT
GTCCAATGGCCACAGCAATGGTGATGGCAAGGTGAACATGTGGCAGCCGGTGCCATGTCGTTTCGTCAAGAAGGATGGA
CACTGCAACGTGCACATCATCAACATGAGCGAGAAAGGCCAGCGCTACATAGCCGACATCTTCACCACCTGCGTGGACATC
CGTTGGCGATGGATGATAATCATCTTCTGCTTGACTTTTGTGCTTTCATGGTTGTTCTTTGGCTATGTGTTCTGGCTGGTGG
CCTTCTTCTATGGTGACTTGGGGAATAGCTCCCAGCAGTGTGTCTCCAATGTCAACAGCTTCATGGCAGCCTTCCTCTTCTCT
GTGGAGACGCAGACCACTATTGGCTATGGTTACCACCATGTGACAGAAGAGTGCCCCATCGCTGTCTTTATGGTGGTTTTC
CAGTGCATTGTTGGCTGCATCATCGACGCCTTCATCATTGGTGCCGTCATGGCCAAGATGGCCAAGCCCACGAAGCGCAAT
GAAACCCTGGTGTTTAGCCACAACGCTACAATAGCAATGCGGGACGGCAAGCTATGCCTGATGTGGCGAGTTGGCAACCT
ACGCAAAAGCCACCTGGTGGAGGCCCACGTGAGGGCTCAGCTACTCAAGTCCCGGACCACCGCCGAGGGGGAGTTTATC
CCCCTAGACCACGTAGATATTGATGTGGGCTTTGACACTGGCGTAGACCGGATCTTCCTTGTTTCCCCCATCACCATTGTCC
ATGAGATCAACGAGGACAGTCCCTTCTATGATATGAGCAAGCAGGATTTTGAGACTGCTGGATTTGAGATTGTGGTCATCC
TGGAGGGCATGGTAGAAGCCACAGCCATGACAACCCAGTGTCGCAGTTCCTACCTGGCAGGGGAGATCCTCTGGGGACA
CTGCTTCGAGCCTGTACTCTTTGAGGAGAAGAACTACTACAAGGTCGACTACTCTCATTTCCACAAAACCTACGAGGTGCC
GAGCACTCCGCTATGTAGTGCGCGGGAGCTTGCTGAAAAGAAGGATAATGAGTCCAGCTCTAACTCTTTTTGCTATGAGA
ATGAAGTGGCGATGATGGACAAAGAGGAGACGGAGGACAAAAGCGAGTGCAGCAATGATGGGAGCAGTTCACAAAAGG
CTTCAGAGTTGGGGCGCAATCTCTTCATGACGTTTAGACGAGAATCTGAGATTTGA

tsh2

ATGAATGTCCAAAACTGTTCCTCAGAAGGTTCTTCCAAAAACTTTTCCAAGGCAGCAGAAGTGAAGCCCTCATCAGCAGAA
GCGATGGGTAGTGTGCGGGCCAGCCGCTACAGCGTCGTGTCCTCCAAAGTAGATGGCCTCAAGTTGGCCACTGTGGCCGT
GTCCAATGGCCACAGCAATGGTGATGGCAAGGTGAACATGTGGCAGCCGGTGCCATGTCGTTTCGTCAAGAAGGATGGA
CACTGCAACGTGCACATCATCAACATGAGCGAGAAAGGCCAGCGCTACATAGCCGACATCTTCACCACCTGCGTGGACATC
CGTTGGCGATGGATGATAATCATCTTCTGCTTGACTTTTGTGCTTTCATGGTTGTTCTTTGGCTATGTGTTCTGGCTGGTGG
CCTTCTTCTATGGTGACTTGGGGAATAGCTCCCAGCAGTGTGTCTCCAATGTCAACAGCTTCATGGCAGCCTTCCTCTTCTCT
GTGGAGACGCAGACCACTATTGGCTATGGTTACCACCATGTGACAGAAGAGTGCCCCATCGCTGTCTTTATGGTGGTTTTC
CAGTGCATTGTTGGCTGCATCATCGACGCCTTCATCATTGGTGCCGTCATGGCCAAGATGGCCAAGCCCACGAAGCGCAAT
GAAACCCTGGTGTTTAGCCACAACGCTACAATAGCAATGCGGGACGGCAAGCTATGCCTGATGTGGCGAGTTGGCAACCT
ACGCAAAAGCCACCTGGTGGAGGCCCACGTGAGGGCTCAGCTACTCAAGTCCCGGACCACCGCCGAGGGGGAGTTTATC
CCCCTAGACCACGTAGATATTGATGTGGGCTTTGACACTGGCGTAGACCGGATCTTCCTTGTTTCCCCCATCACCATTGTCC
ATGAGATCAACGAGGACAGTCCCTTCTATGATATGAGCAAGCAGGATTTTGAGACTGCTGGATTTGAGATTGTGGTCATCC
TGGAGGGCATGGTAGAAGCCACAGCCATGACAACCCAGTGTCGCAGTTCCTACCTGGCAGGGGAGATCCTCTGGGGACA
CTGCTTCGAGCCTGTACTCTTTGAGGAGAAGAACTACTACAAGGTCGACTACTCTCATTTCCACAAAACCTACGAGGTGCC
GAGCACTCCGCTATGTAGTGCGCGGGAGCTTGCTGAAAAGAAGGATAATGAGTCCAGCTCTAACTCTTTTTGCTATGAGA
ATGAAGTGGCGATGATGGACAAAGAGGAGACGGAGGACAAAAGCGAGTGCAGCAATGATGGGAGCAGTTCACAAAAGG
CTTCAGAGTTGGGGCGCAATCTCTTCATGACGTTTAGACGAGAATCTGAGATTTGA

| | |
|---|---|
| tsh3 | ATGAATGTCCAAAACTGTTCCTCAGAAGGTTCTTCCAAAAACTTTTCCAAGGCAGCAGAAGTGAAGCCCTCATCAGCAGAA<br>GCGATGGGTAGTGTGCGGGCCAGCCGCTACAGCGTCGTGTCCTCCAAAGTAGATGGCCTCAAGTTGGCCACTGTGGCCGT<br>GTCCAATGGCCACAGCAATGGTGATGGCAAGGTGAACATGTGGCAGCCGGTGCCATGTCGTTTCGTCAAGAAGGATGGA<br>CACTGCAACGTGCACATCATCAACATGAGCGAGAAAGGCCAGCGCTACATAGCCGACATCTTCACCACCTGCGTGGACATC<br>CGTTGGCGATGGATGATAATCATCTTCTGCTTGACTTTTGTGCTTTCATGGTTGTTCTTTGGCTATGTGTTCTGGCTGGTGG<br>CCTTCTTCTATGGTGACTTGGGGAATAGCTCCCAGCAGTGTGTCTCCAATGTCAACAGCTTCATGGCAGCCTTCCTCTTCTCT<br>GTGGAGACGCAGACCACTATTGGCTATGGTTACCACCATGTGACAGAAGAGTGCCCCATCGCTGTCTTTATGGTGGTTTTC<br>CAGTGCATTGTTGGCTGCATCATCGACGCCTTCATCATTGGTGCCGTCATGGCCAAGATGGCCAAGCCCACGAAGCGCAAT<br>GAAACCCTGGTGTTTAGCCACAACGCTACAATAGCAATGCGGGACGGCAAGCTATGCCTGATGTGGCGAGTTGGCAACCT<br>ACGCAAAAGCCACCTGGTGGAGGCCCACGTGAGGGCTCAGCTACTCAAGTCCCGGACCACCGCCGAGGGGGAGTTTATC<br>CCCCTAGACCACGTAGATATTGATGTGGGCTTTGACACTGGCGTAGACCGGATCTTCCTTGTTTCCCCCATCACCATTGTCC<br>ATGAGATCAACGAGGACAGTCCCTTCTATGATATGAGCAAGCAGGATTTTGAGACTGCTGGATTTGAGATTGTGGTCATCC<br>TGGAGGGCATGGTAGAAGCCACAGCCATGACAACCCAGTGTCGCAGTTCCTACCTGGCAGGGGAGATCCTCTGGGGACA<br>CTGCTTCGAGCCTGTACTCTTTGAGGAGAAGAACTACTACAAGGTCGACTACTCTCATTTCCACAAAACCTACGAGGTGCC<br>GAGCACTCCGCTATGTAGTGCGCGGGAGCTTGCTGAAAAGAAGGATAATGAGTCCAGCTCTAACTCTTTTTGCTATGAGA<br>ATGAAGTGGCGATGATGGACAAAGAGGAGACGGAGGACAAAAGCGAGTGCAGCAATGATGGGAGCAGTTCACAAAAGG<br>CTTCAGAGTTGGGGCGCAATCTCTTCATGACGTTTAGACGAGAATCTGAGATTTGA |

rhy1

ATGAATGTCCAAAACTGTTCCTCAGAAGGTTCTTCCAAAAACTTTTCCAAGGCAGCAGAAGTGAAGCCCTCATCAGCAGAA
GCGATGGGTAGTGTGCGGGCCAGCCGCTACAGCGTCGTGTCCTCCAAAGTAGATGGCCTCAAGTTGGCCACTGTGGCCGT
GTCCAATGGCCACAGCAGTGGTGATGGCAAGGTGAACATGTGGCAGCCGGTGCCATGTCGTTTCGTCAAGAAGGATGGA
CACTGCAACGTGCACATCATCAACATGAGCGAGAAAGGCCAGCGCTACATAGCCGACATCTTCACCACCTGCGTGGACATC
CGTTGGCGATGGATGATAATCATCTTCTGCTTGACTTTTGTGCTTTCATGGTTGTTCTTTGGCTATGTGTTCTGGCTGGTGG
CCTTCTTCTATGGTGACTTGGGGAATAGCTCCCAGCAGTGTGTCTCCAATGTCAACAGCTTCATGGCAGCCTTCCTCTTCTCT
GTGGAAACGCAGACCACTATTGGCTATGGTTACCACCATGTGACAGAAGAGTGCCCCATCGCTGTCTTTATGGTGGTTTTC
CAGTGCATTGTTGGCTGCATCATCAACGCCTTCATCATTGGTGCCGTCATGGCCAAGATGGCCAAGCCCACGAAGCGCAAT
GAAACCCTGGTGTTTAGCCACAACGCTACAATAGCAATGCGGGACGGTAAGCTATGCCTGATGTGGCGAGTTGGCAACCT
ACGCAAAAGCCACCTGGTGGAGGCCCACGTGAGGGCTCAGCTACTCAAGTCCCGGACCACCGCCGAGGGGGAGTTTATC
CCCCTAGACCACGTAGATATTGATGTGGGCTTTGACACTGGCGTAGACCGGATCTTCCTTGTTTCCCCCATCACCATTGTCC
ATGAGATCAACGAGGACAGTCCCTTCTATGATATGAGCAAGCAGGATTTTGAGACTGCTGGATTTGAGATTGTGGTCATCC
TGGAGGGCATGGTAGAAGCCACAGCCATGACAACCCAGTGTCGCAGTTCCTACCTGGCAGGGGAGATCCTCTGGGGACA
CTGCTTCGAGCCTGTACTCTTTGAGGAGAAGAACTACTACAAGGTCGACTACTCTCATTTCCACAAAACCTACGAGGTGCC
GAGCACTCCGCTATGTAGTGCGCGGGAGCTTGCTGAAAAGAAGGATAATGAGTCCAGCTCTAACTCTTTTTGCTATGAGA
ATGAAGTGGCGATGATGGACAAAGAGGAGACGGAGGACAAAAGCGAGTGCAGCAATGATGGGAGCAGTTCACAAAAGG
CTTCAGAGTTGGGGCGCAATCTCTTCATGACGTTTAGACGAGAATCTGAGATTTGA

rhy2

ATGAATGTCCAAAACTGTTCCTCAGAAGGTTCTTCCAAAAACTTTTCCAAGGCAGCAGAAGTGAAGCCCTCATCAGCAGAA
GCGATGGGTAGTGTGCGGGCCAGCCGCTACAGCGTCGTGTCCTCCAAAGTAGATGGCCTCAAGTTGGCCACTGTGGCCGT
GTCCAATGGCCACAGCAGTGGTGATGGCAAGGTGAACATGTGGCAGCCGGTGCCATGTCGTTTCGTCAAGAAGGATGGA
CACTGCAACGTGCACATCATCAACATGAGCGAGAAAGGCCAGCGCTACATAGCCGACATCTTCACCACCTGCGTGGACATC
CGTTGGCGATGGATGATAATCATCTTCTGCTTGACTTTTGTGCTTTCATGGTTGTTCTTTGGCTATGTGTTCTGGCTGGTGG
CCTTCTTCTATGGTGACTTGGGGAATAGCTCCCAGCAGTGTGTCTCCAATGTCAACAGCTTCATGGCAGCCTTCCTCTTCTCT
GTGGANACGCAGACCACTATTGGCTATGGTTACCACCATGTGACAGAAGAGTGCCCCATCGCTGTCTTTATGGTGGTTTTC
CAGTGCATTGTTGGCTGCATCATCAACGCCTTCATCATTGGTGCCGTCATGGCCAAGATGGCCAAGCCCACGAAGCGCAAT
GAAACCCTGGTGTTTAGCCACAACGCTACAATAGCAATGCGGGACGGTAAGCTATGCCTGATGTGGCGAGTTGGCAACCT
ACGCAAAAGCCACCTGGTGGAGGCCCACGTGAGGGCTCAGCTACTCAAGTCCCGGACCACCGCCGAGGGGGAGTTTATC
CCCCTAGACCACGTAGATATTGATGTGGGCTTTGACACTGGCGTAGACCGGATCTTCCTTGTTTCCCCCATCACCATTGTCC
ATGAGATCAACGAGGACAGTCCCTTCTATGATATGAGCAAGCAGGATTTTGAGACTGCTGGATTTGAGATTGTGGTCATCC
TGGAGGGCATGGTAGAAGCCACAGCCATGACAACCCAGTGTCGCAGTTCCTACCTGGCAGGGGAGATCCTCTGGGGACA
CTGCTTCGAGCCTGTACTCTTTGAGGAGAAGAACTACTACAAGGTCGACTACTCTCATTTCCACAAAACCTACGAGGTGCC
GAGCACTCCGCTATGTAGTGCGCGGGAGCTTGCTGAAAAGAAGGATAATGAGTCCAGCTCTAACTCTTTTTGCTATGAGA
ATGAAGTGGCGATGATGGACAAAGAGGAGACGGAGGACAAAAGCGAGTGCAGCAATGATGGGAGCAGTTCACAAAAGG
CTTCAGAGTTGGGGCGCAATCTCTTCATGACGTTTAGACGAGAATCTGAGATTTGA

rhy3 ATGAATGTCCAAAACTGTTCCTCAGAAGGTTCTTCCAAAAACTTTTCCAAGGCAGCAGAAGTGAAGCCCTCATCAGCAGAA
GCGATGGGTAGTGTGCGGGCCAGCCGCTACAGCGTCGTGTCCTCCAAAGTAGATGGCCTCAAGTTGGCCACTGTGGCCGT
GTCCAATGGCCACAGCAGTGGTGATGGCAAGGTGAACATGTGGCAGCCGGTGCCATGTCGTTTCGTCAAGAAGGATGGA
CACTGCAACGTGCACATCATCAACATGAGCGAGAAAGGCCAGCGCTACATAGCCGACATCTTCACCACCTGCGTGGACATC
CGTTGGCGATGGATGATAATCATCTTCTGCTTGACTTTTGTGCTTTCATGGTTGTTCTTTGGCTATGTGTTCTGGCTGGTGG
CCTTCTTCTATGGTGACTTGGGGAATAGCTCCCAGCAGTGTGTCTCCAATGTCAACAGCTTCATGGCAGCCTTCCTCTTCTCT
GTGGANACGCAGACCACTATTGGCTATGGTTACCACCATGTGACAGAAGAGTGCCCCATCGCTGTCTTTATGGTGGTTTTC
CAGTGCATTGTTGGCTGCATCATCAACGCCTTCATCATTGGTGCCGTCATGGCCAAGATGGCCAAGCCCACGAAGCGCAAT
GAAACCCTGGTGTTTAGCCACAACGCTACAATAGCAATGCGGGACGGTAAGCTATGCCTGATGTGGCGAGTTGGCAACCT
ACGCAAAAGCCACCTGGTGGAGGCCCACGTGAGGGCTCAGCTACTCAAGTCCCGGACCACCGCCGAGGGGGAGTTTATC
CCCCTAGACCACGTAGATATTGATGTGGGCTTTGACACTGGCGTAGACCGGATCTTCCTTGTTTCCCCCATCACCATTGTCC
ATGAGATCAACGAGGACAGTCCCTTCTATGATATGAGCAAGCAGGATTTTGAGACTGCTGGATTTGAGATTGTGGTCATCC
TGGAGGGCATGGTAGAAGCCACAGCCATGACAACCCAGTGTCGCAGTTCCTACCTGGCAGGGGAGATCCTCTGGGGACA
CTGCTTCGAGCCTGTACTCTTTGAGGAGAAGAACTACTACAAGGTCGACTACTCTCATTTCCACAAAACCTACGAGGTGCC
GAGCACTCCGCTATGTAGTGCGCGGGAGCTTGCTGAAAAGAAGGATAATGAGTCCAGCTCTAACTCTTTTTGCTATGAGA
ATGAAGTGGCGATGATGGACAAAGAGGAGACGGAGGACAAAAGCGAGTGCAGCAATGATGGGAGCAGTTCACAAAAGG
CTTCAGAGTTGGGGCGCAATCTCTTCATGACGTTTAGACGAGAATCTGAGATTTGA

rhy4 ATGAATGTCCAAAACTGTTCCTCAGAAGGTTCTTCCAAAAACTTTTCCAAGGCAGCAGAAGTGAAGCCCTCATCAGCAGAA
GCGATGGGTAGTGTGCGGGCCAGCCGCTACAGCGTCGTGTCCTCCAAAGTAGATGGCCTCAAGTTGGCCACTGTGGCCGT
GTCCAATGGCCACAGCAGTGGTGATGGCAAGGTGAACATGTGGCAGCCGGTGCCATGTCGTTTCGTCAAGAAGGATGGA
CACTGCAACGTGCACATCATCAACATGAGCGAGAAAGGCCAGCGCTACATAGCCGACATCTTCACCACCTGCGTGGACATC
CGTTGGCGATGGATGATAATCATCTTCTGCTTGACTTTTGTGCTTTCATGGTTGTTCTTTGGCTATGTGTTCTGGCTGGTGG
CCTTCTTCTATGGTGACTTGGGGAATAGCTCCCAGCAGTGTGTCTCCAATGTCAACAGCTTCATGGCAGCCTTCCTCTTCTCT
GTGGAAACGCAGACCACTATTGGCTATGGTTACCACCATGTGACAGAAGAGTGCCCCATCGCTGTCTTTATGGTGGTTTTC
CAGTGCATTGTTGGCTGCATCATCAACGCCTTCATCATTGGTGCCGTCATGGCCAAGATGGCCAAGCCCACGAAGCGCAAT
GAAACCCTGGTGTTTAGCCACAACGCTACAATAGCAATGCGGGACGGTAAGCTATGCCTGATGTGGCGAGTTGGCAACCT
ACGCAAAAGCCACCTGGTGGAGGCCCACGTGAGGGCTCAGCTACTCAAGTCCCGGACCACCGCCGAGGGGGAGTTTATC
CCCCTAGACCACGTAGATATTGATGTGGGCTTTGACACTGGCGTAGACCGGATCTTCCTTGTTTCCCCCATCACCATTGTCC
ATGAGATCAACGAGGACAGTCCCTTCTATGATATGAGCAAGCAGGATTTTGAGACTGCTGGATTTGAGATTGTGGTCATCC
TGGAGGGCATGGTAGAAGCCACAGCCATGACAACCCAGTGTCGCAGTTCCTACCTGGCAGGGGAGATCCTCTGGGGACA
CTGCTTCGAGCCTGTACTCTTTGAGGAGAAGAACTACTACAAGGTCGACTACTCTCATTTCCACAAAACCTACGAGGTGCC
GAGCACTCCGCTATGTAGTGCGCGGGAGCTTGCTGAAAAGAAGGATAATGAGTCCAGCTCTAACTCTTTTTGCTATGAGA
ATGAAGTGGCGATGATGGACAAAGAGGAGACGGAGGACAAAAGCGAGTGCAGCAATGATGGGAGCAGTTCACAAAAGG
CTTCAGAGTTGGGGCGCAATCTCTTCATGACGTTTAGACGAGAATCTGAGATTTGA

rhy5 ATGAATGTCCAAAACTGTTCCTCAGAAGGTTCTTCCAAAAACTTTTCCAAGGCAGCAGAAGTGAAGCCCTCATCAGCAGAA
GCGATGGGTAGTGTGCGGGCCAGCCGCTACAGCGTCGTGTCCTCCAAAGTAGATGGCCTCAAGTTGGCCACTGTGGCCGT
GTCCAATGGCCACAGCAGTGGTGATGGCAAGGTGAACATGTGGCAGCCGGTGCCATGTCGTTTCGTCAAGAAGGATGGA
CACTGCAACGTGCACATCATCAACATGAGCGAGAAAGGCCAGCGCTACATAGCCGACATCTTCACCACCTGCGTGGACATC
CGTTGGCGATGGATGATAATCATCTTCTGCTTGACTTTTGTGCTTTCATGGTTGTTCTTTGGCTATGTGTTCTGGCTGGTGG
CCTTCTTCTATGGTGACTTGGGGAATAGCTCCCAGCAGTGTGTCTCCAATGTCAACAGCTTCATGGCAGCCTTCCTCTTCTCT
GTGGANACGCAGACCACTATTGGCTATGGTTACCACCATGTGACAGAAGAGTGCCCCATCGCTGTCTTTATGGTGGTTTTC
CAGTGCATTGTTGGCTGCATCATCAACGCCTTCATCATTGGTGCCGTCATGGCCAAGATGGCCAAGCCCACGAAGCGCAAT
GAAACCCTGGTGTTTAGCCACAACGCTACAATAGCAATGCGGGACGGTAAGCTATGCCTGATGTGGCGAGTTGGCAACCT
ACGCAAAAGCCACCTGGTGGAGGCCCACGTGAGGGCTCAGCTACTCAAGTCCCGGACCACCGCCGAGGGGGAGTTTATC
CCCCTAGACCACGTAGATATTGATGTGGGCTTTGACACTGGCGTAGACCGGATCTTCCTTGTTTCCCCCATCACCATTGTCC
ATGAGATCAACGAGGACAGTCCCTTCTATGATATGAGCAAGCAGGATTTTGAGACTGCTGGATTTGAGATTGTGGTCATCC
TGGAGGGCATGGTAGAAGCCACAGCCATGACAACCCAGTGTCGCAGTTCCTACCTGGCAGGGGAGATCCTCTGGGGACA
CTGCTTCGAGCCTGTACTCTTTGAGGAGAAGAACTACTACAAGGTCGACTACTCTCATTTCCACAAAACCTACGAGGTGCC
GAGCACTCCGCTATGTAGTGCGCGGGAGCTTGCTGAAAAGAAGGATAATGAGTCCAGCTCTAACTCTTTTTGCTATGAGA
ATGAAGTGGCGATGATGGACAAAGAGGAGACGGAGGACAAAAGCGAGTGCAGCAATGATGGGAGCAGTTCACAAAAGG
CTTCAGAGTTGGGGCGCAATCTCTTCATGACGTTTAGACGAGAATCTGAGATTTGA

**Supplementary Table 9** Prediction of impact of two amino acid substitutions inferred from KCNJ2 transcripts of *com, tsh* and *rhy* .

| Site(aa) | *com* | *tsh* | *rhy* | Score | Sensitivity | Specificity | Prediction |
|---|---|---|---|---|---|---|---|
| 60 | N | N | S | 0,223 | 0,91 | 0,88 | Benign |
| 198 | D | D | N | 0,983 | 0,74 | 0,96 | Probably damaging |
